# Tryptophan Chemistry Driven by a Widespread Cytochrome P422 Enzyme Family

**DOI:** 10.64898/2025.12.11.692455

**Authors:** Wencheng Ma, Qian Wang, Qingyu Yang, Zhaojie Teng, Xiao Han, Moli Sang, Qiuyu Li, Ruili Wang, Peiyuan Feng, Jingyi Zhong, Yan Zhang, Yifeng Wei, Li Jiang, F. Peter Guengerich, Wei Zhang

**Affiliations:** Laboratory of Experimental Marine Biology, Institute of Oceanology, Chinese Academy of Sciences, Qingdao, 266000, China; State Key Laboratory of Microbial Technology, Shandong University, Qingdao, Shandong, 266237, China; Laboratory for Marine Biology and Biotechnology, Qingdao Marine Science and Technology Center, Qingdao, Shandong, 266237, China; Key Laboratory of Marine Drugs, Chinese Ministry of Education, School of Medicine and Pharmacy, Ocean University of China, Qingdao 266003, China; State Key Laboratory of Microbial Metabolism, School of Life Sciences and Biotechnology, Shanghai Jiao Tong University, Shanghai 200240, China; Singapore Institute of Food and Biotechnology Innovation (SIFBI), Agency for Science, Technology and Research (A*STAR), 31 Biopolis Way, Nanos, Singapore 138669, Singapore; New Cornerstone Science Laboratory, School of Pharmaceutical Science and Technology, Tianjin University, Tianjin 300072, China; Department of Biochemistry, Vanderbilt University School of Medicine, Nashville, Tennessee 37232-0146

**Author notes:** To whom correspondence should be addressed (W.Z.). These authors contribute equally to this work (Wencheng Ma, Qian Wang, Qingyu Yang).

## Abstract

Tryptophan serves as a versatile biosynthetic precursor across living organisms. While heme-binding proteins (HBPs) mediate key reactions in tryptophan transformation, the full diversity of HBPs remains largely unexplored. Here, we developed the novel Cofactor-Integrative Structural Inspector (CISSspector) to systematically identify HBPs in the extensive extant microbial genomic sequence database, which revealed several uncharacterized HBP families. We experimentally characterized one of the most prominent families, the cytochrome P422 (formerly DUF6875) family, distributed throughout the prokaryotes and eukaryotes. Strikingly, we discovered that this enzyme family orchestrates four chemically distinct and biochemically unprecedented transformations, with regioselectivity, including N1-, C6-, and C7-hydroxylations and intramolecular C–S bond formations. Notably, the discovery of enzymes capable of Trp N1- and C7-hydroxylation addresses a long-standing gap in the natural enzyme arsenal. Structural analysis of the representative cytochrome P422 enzyme Mc170 revealed a structurally unique HBP fold in which conserved residues form a substrate “clamp” that positions the tryptophan indole ring for selective modification. Our work unveils a hidden enzymatic repertoire of HBPs, expands the known landscape of tryptophan metabolism, and establishes an artificial intelligence-augmented framework for discovering cryptic enzymes with broad implications for synthetic biology and natural product discovery.

## Introduction

Tryptophan, the most structurally complex canonical amino acid, serves as a universal biosynthetic precursor across all domains of life^1^. Currently, Trp metabolism is known to be carried out by a plethora of enzymes through distinct biochemical routes, with the canonical metabolic model comprising at least three major pathways. In mammals, Trp catabolism primarily proceeds through the kynurenine pathway, producing neuroactive and immunomodulatory metabolites^2^, while the serotonin-melatonin pathway regulates neuroendocrine functions via initial Trp hydroxylation^2^. In plants and microbes, Trp is transformed into hormones, antibiotics, and virulence factors through diverse chemical reactions including hydroxylation, decarboxylation, and elimination^3^. These chemical reactions are often catalyzed by heme-binding proteins (HBPs) such as histidine-ligated Trp 2,3-dioxygenases (TDOs)^4^ and cysteine-ligated cytochrome P450s^5^ (Fig. 1a and b). The subsequent discovery of histidine-ligated aromatic oxygenases further expanded the metabolic pathways of Trp derivatives^6^. These pathways highlight the intricate and diverse nature of known enzyme systems dedicated to mediating Trp transformations.

**Fig. 1.**
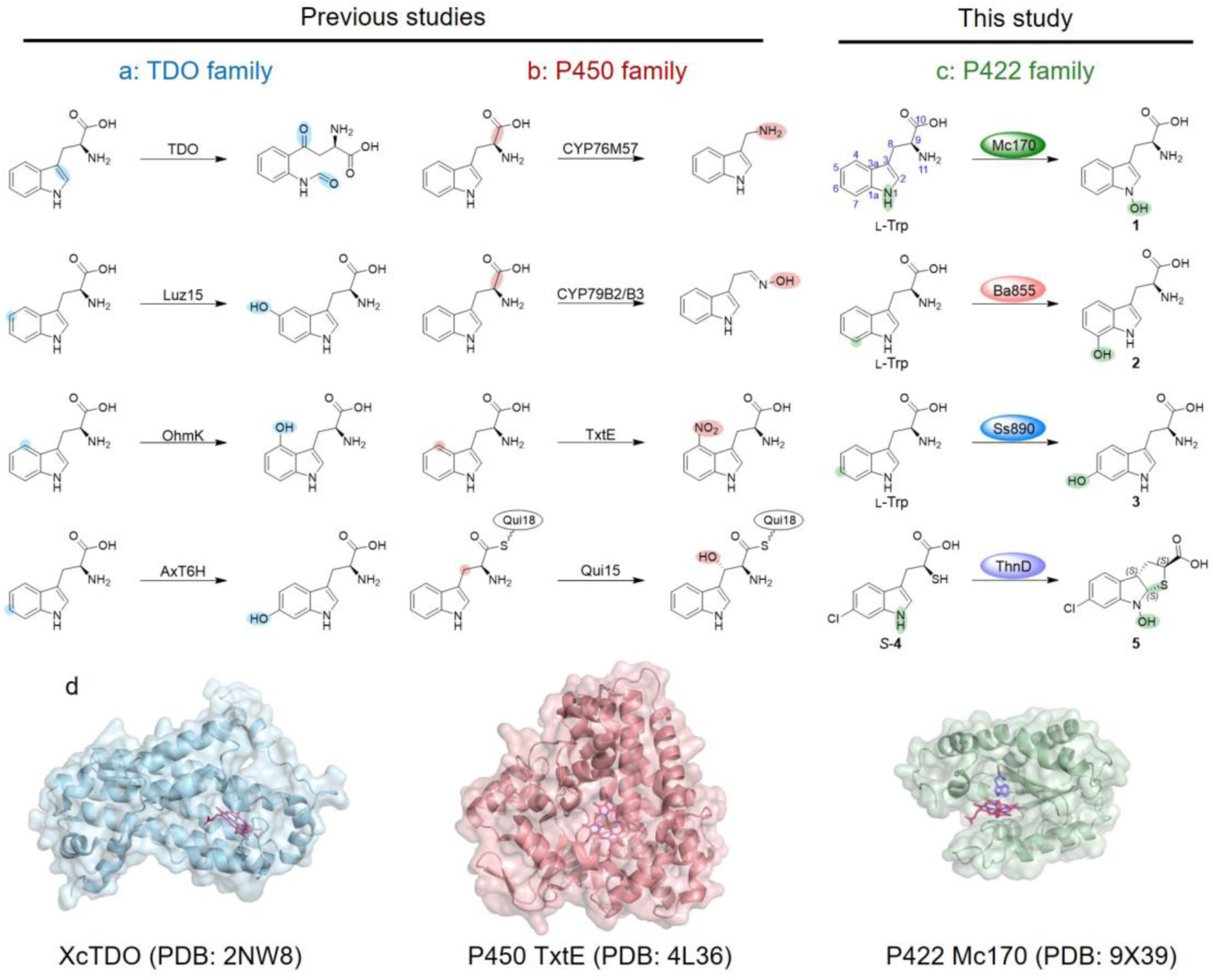
Trp metabolic pathways and overall crystal structures. **a**, Metabolic pathways catalyzed by TDOs. **b**, Metabolic pathways mediated by P450s. **c**, The novel metabolic pathways performed by members of the cytochrome P422 family (this study). **d**, Overall crystal structures of XcTDO (PDB: 2NW8), P450 TxtE (PDB: 4L36), and P422 Mc170 (PDB: 9X39).

During our study of microbial Trp-derived natural product biosynthesis, we found that the full extent of enzymatic machinery dedicated to tryptophan modification remains largely uncharted. This knowledge gap is particularly pronounced in genetically divergent microorganisms from underexplored environments, whose biosynthetic potential has been scarcely investigated due to cultivation limitations and analytical challenges. The discovery of novel Trp-modifying enzymes from these microbial reservoirs has been fundamentally constrained by the limitations of conventional bioinformatic approaches. State-of-the-art prediction tools such as CLEAN^7^ and EC-Transformer^8^ depend on sequence homology and known catalytic motifs, rendering them ineffective for identifying novel enzyme families that share neither sequence similarity nor structural features with characterized proteins (“dark-matter” enzymes). This methodological gap is especially problematic for HBPs, where the same cofactor can be coordinated through diverse structural solutions that evade detection by sequence-based methods. Consequently, a vast repository of catalytic diversity dedicated to Trp functionalization remains hidden within microbial genomes, requiring innovative strategies that transcend sequence homology to unlock nature’s full enzymatic repertoire for this crucial biosynthetic precursor.

In this work, we first developed the Cofactor-Integrative Structural Inspector (CISSspector), a hybrid approach combining deep learning-aided structural prediction with genomic mining to systematically identify non-canonical HBPs. In further screening of 100 deep-sea actinomycete genomes, CISSspector uncovered 23 previously unrecognized potential HBP families, most lacking functional annotation. Strikingly, the DUF6875 family, which is a thiolate-ligated hemoprotein family, co-localized with putative Trp metabolic genes despite exhibiting no sequence or structural homology to known HBPs (Fig. 1d). Following the nomenclature of the pioneering cytochrome P450 and P411^9^, we redesignate the DUF6875 family as the cytochrome P422 family in this work.

Selective indole C–H hydroxylation presents a persistent challenge in synthetic chemistry. Although metal-free, site-selective functionalizations have been developed, they largely depend on the installation and removal of directing groups^10^. Our functional characterization of the cytochrome P422 enzyme family reveals four distinct transformations in tryptophan metabolism: regioselective N1-, C6-, and C7-hydroxylations, and intramolecular C–S bond formation (Fig. 1c). Notably, the discovery of dedicated N1- and C7-hydroxylases addresses a long-standing gap in biology, making these otherwise challenging hydroxyindole motifs readily accessible through biocatalysis.

Remarkably, these transformations occur despite cytochrome P422 lacking sequence or structural homology to any canonical HBPs, challenging existing paradigms of enzyme evolution. The discovery of such atypical chemistry demonstrates how integrating structural genomics with functional analysis can uncover hidden enzymatic diversity that eludes traditional prediction methods. Beyond advancing our understanding of Trp metabolism, these findings open new avenues for exploring unconventional biocatalytic reactions with potential applications in synthetic biology, metabolic engineering, and the discovery of pharmacologically relevant alkaloid scaffolds.

### 1 Development and validation of CISSspector

We present the Cofactor-Integrative Structural Inspector (CISSspector), a computational workflow designed to identify HBPs through the integration of predicted protein structures and confidence metrics, offering a more robust screening approach than sequence-based methods alone (Fig. 2a). The Boltz-1 model was utilized for all protein-ligand complex predictions^11^, with its confidence scores serving as key inputs for subsequent machine learning, and the inference efficiency was further optimized (see Inference Optimization in Methods). Models were trained on a curated dataset A2 containing 400 protein sequences (200 HBPs and 200 non-HBPs), and then tested on an independently sampled dataset A1 of 100 sequences and a negative dataset B of 1000 sequences sourced from reviewed UniProtKB entries^12^, both after homology reduction (see Dataset Preparation in Methods and Extended Data Fig. 1). Key structural features, including the number of heme-surrounding residues and count of coordinating residues, were extracted from predicted structures using a custom script tailored for heme-binding characterization and were subsequently integrated with confidence scores including complex_iplddt and iPTM to form sample features (see Feature Selection in Methods).

**Fig. 2.**
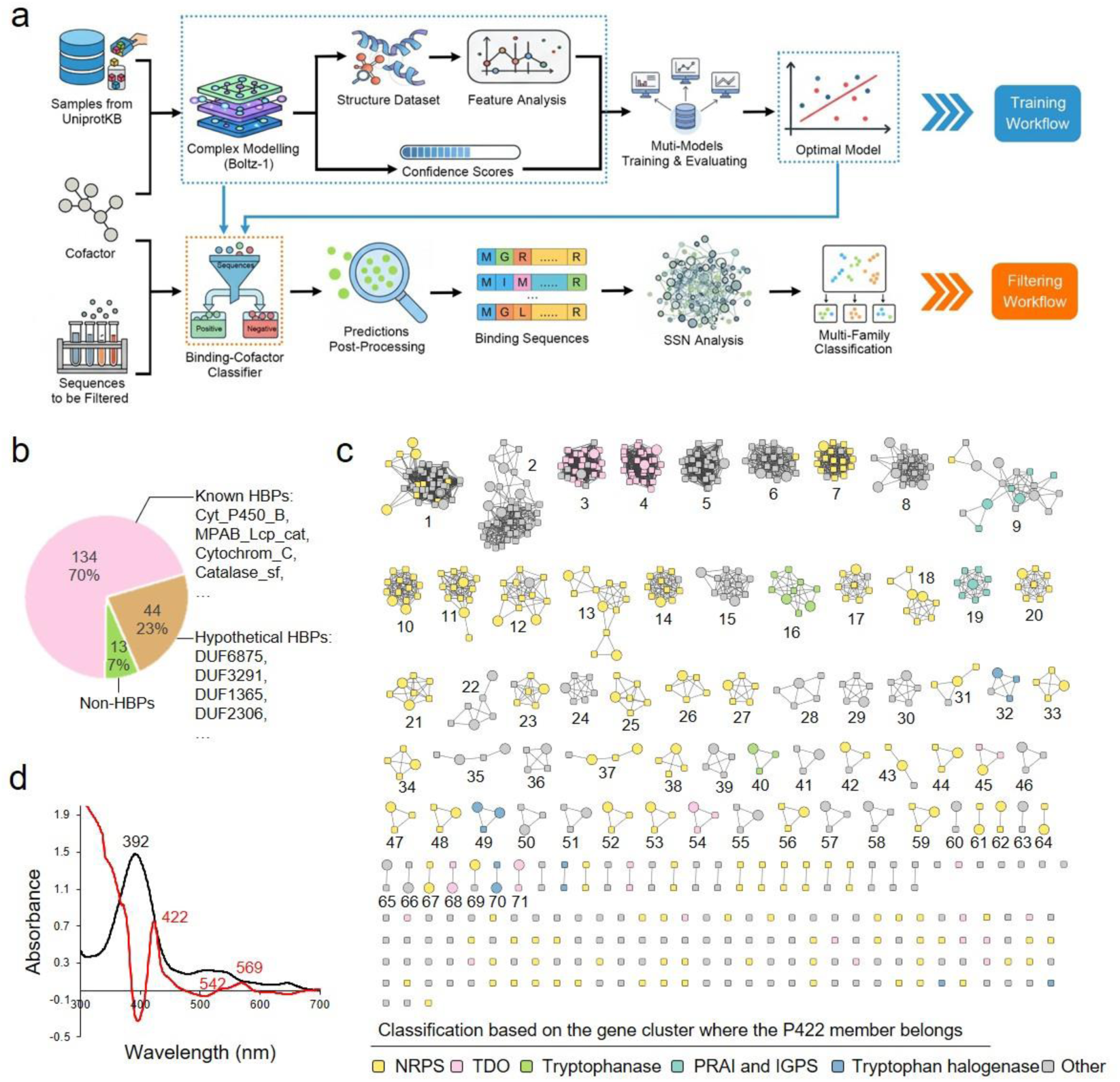
Development of CISSspector and HBP analysis. **a**, CISSspector workflow. Schematic diagram of the HBP screening pipeline. **b**, Pie chart analysis of CISSspector hits. **c**, Sequence similarity network (SSN) analysis of cytochrome P422 family. A protein SSN was constructed for the cytochrome P422 family (Pfam number: PF21780) using a sequence length of 180-300 amino acids, and a total of 707 sequences were retrieved. To achieve comprehensive functional diversity while minimizing redundancy, we performed clustering with a sequence identity threshold of 65%. From each cluster, 1–5 representative sequences were selected for functional validation, resulting in 103 chosen sequences (large circular nodes). This selection accounted for 14.6% of the total cytochrome P422 members registered in UniProtKB. Different colors indicate the classification of Trp-related genes or NRPS gene clusters where cytochrome P422 member genes are located. **d**, UV-visible absorption spectrum for Mc170 in oxidized form (black line) and CO-bound reduced difference form by sodium hydrosulfite (dithionite) (red line).

Three machine learning models were further trained and evaluated, including logistic regression (LR)^13^, naive Bayes (NB)^14^, and random forest (RF)^15^. Grid search (GR)^16^ was used to find the optimal threshold combination for direct screening as a control (see Training Details in Methods and Supplementary Table 1). All four algorithms were tested independently after training (Extended Data Fig. 1b-e). In three positive-sample benchmark scores, LR achieved the highest precision (0.89) and F1 score (0.90), while LR and RF shared the highest recall (0.91). During the testing on negative samples, RF achieved the highest accuracy (0.96), with LR coming in second place (0.94). Therefore, LR was chosen as the model for practical classification due to its better performance. In contrast, the classification performance of GR for positive and negative samples was significantly weaker than that of the three algorithms, demonstrating the necessity of employing machine learning algorithms (Extended Data Fig. 1).

For validation, the screening of a total of 7,333 protein sequences from *Streptomyces coelicolor* A3(2) by CISSspector led to the discovery of 53 candidates, 44 of which were manually annotated as HBPs including the 18 known P450 enzymes, which provides practical support for the reliability of CISSspector (Supplementary Table 2).

### 2 Global genome mining and genomic neighborhood analysis of HBPs

Screening of 100 deep-sea-derived actinomycete proteomes using CISSpector identified 191 putative heme-binding protein (HBP) sequences (Supplementary Figs. 1 and 2 and Supplementary Table 3). To classify the candidate proteins, all 191 sequences were analyzed by BLASTP with the UniProt database. Based on BLASTP-derived family and domain annotations, the 191 proteins were categorized into 134 known HBPs, 13 non-HBPs, and 44 unannotated proteins designated as hypothetical HBPs. The known HBPs grouped into 20 families, comprising cytochrome P450s, DUF2236 (fungal oxygenase MpaB’ and rubber oxygenase^17,18^), TDOs, and others (Supplementary Table 3). We further analyzed the hypothetical HBPs by constructing a sequence similarity network (SSN) at 25% identity using EFI-EST^19,20^. Strikingly, clustering revealed 23 distinct families, the majority of which are large, widely distributed, and functionally uncharacterized; these include DUF1365, DUF2306, DUF3291, and DUF6875 (Fig. 2b and Supplementary Fig. 2). These results demonstrate the capacity of CISSpector to systematically identify both novel and known HBP families with high precision.

Little is known about the role of the DUF6875 family (renamed in this study as cytochrome P422), even for members that are associated with the biosynthetic gene clusters for natural products including the antitumorigenic actinomycin^21,22^, the antibiotic mycemycin^23,24^, and the anti-bacterial agent thienodolin^25,26^ (Extended Data Fig. 2). Surprisingly, the CLEAN algorithm annotated cytochrome P422 members as ferredoxin oxidoreductase (EC 1.3.7.3) or thymidylate synthase (EC 2.1.1.148) (Supplementary Fig. 3 and Supplementary Table 4), which highlights its limitation in predicting novel enzymatic functions.

To gain deep functional insights into cytochrome P422, we constructed SSN using EFI-EST^19,20^ with the Pfam family PF21780 (Fig. 2c). Subsequently, an *E* value of 1×10^−40^, sequence length constraints (180–300 aa), and 65% identity clustering were used for edge generation, which retrieved 707 sequences derived from diverse prokaryotic phyla (*Actinomycetota*, *Cyanobacteriota*, *Pseudomonadota*, *Myxococcota*, *Bacteroidota*, *Bacillota*) and eukaryotes (*Rotifera*) (Supplementary Tables 5 and 6). Genomic neighborhood (GN) analysis further revealed frequent co-localization of cytochrome P422 with Trp metabolism enzymes including TDO, kynureninase, tryptophanase, phosphoribosylanthranilate isomerase (PRAI), and indole-3-glycerol phosphate synthase (IGPS) (Fig. 2c; Supplementary Fig. 4).

### 3 Functional characterizations of the cytochrome P422 family

To investigate the functions of the cytochrome P422 family, we randomly selected 103 candidates from different SSN clusters and synthesized them with codon optimization, to be expressed in *Escherichia coli* BL21(DE3); we successfully obtained 84 soluble proteins, originating from diverse phyla including *Actinomycetota* (52), *Pseudomonadota* (18), *Cyanobacteriota* (10), *Myxococcota* (2), *Bacillota* (1), and *Bacteroidota* (1) (Supplementary Tables 5 and 6).

The purified proteins exhibited a distinct brownish-red coloration and displayed Soret absorption maxima ranging between 390 nm and 416 nm, consistent with heme incorporation^27^ (Supplementary Data 1). Further characterization using CO-binding assays with dithionite-reduced protein revealed a characteristic Soret band shift to 422 nm (Fig. 2d), a diagnostic feature of HBPs^28^. The combination of ICP-MS and quantitation assays revealed that family member ThnD contains iron and binds iron in an equimolar ratio. This iron was subsequently identified as a non-covalently bound *b*-type heme cofactor by liquid chromatography-high resolution mass spectrometry (LC-HRMS) analysis (Supplementary Fig. 5). Based on these analysis and conserved spectral signature, we thus propose the nomenclature of cytochrome P422 for the DUF6875 family.

Ten potential substrates, including L-Trp and its derivatives, were systematically evaluated by *in vitro* functional reconstruction in the presence of the redox partners of ferredoxin Fdx1499 and ferredoxin reductase FdR0978 from *Synechococcus elongatus* PCC 7942 along with sodium ascorbate and NADPH^29^ (Extended Data Fig. 3a). Enzymes known to catalyze hydroxylation of the indole ring of free Trp are rare^30^. High-performance liquid chromatography (HPLC) and LC-HRMS analyses revealed three distinct types of tryptophan oxidation products. Among them, 41 enzymes (e.g., Mc170 from *Mucilaginibacter corticis*) catalyzed the formation of compound **1**, which exhibited two key mass features: a mass 2 amu lower than that of L-Trp (*m*/*z* 203.0813, [M-H_2_O+H]^+^), indicative of dehydration, and a mass 16 amu higher (*m*/*z* 221.0813, [M+H]^+^), consistent with hydroxylation (Fig. 3a, b). Additionally, negative-ion LC-HRMS confirmed the [M-H]^−^ ion peak at *m/z* 219.0771, further supporting the hydroxylation reaction (Supplementary Fig. 6).

**Fig. 3.**
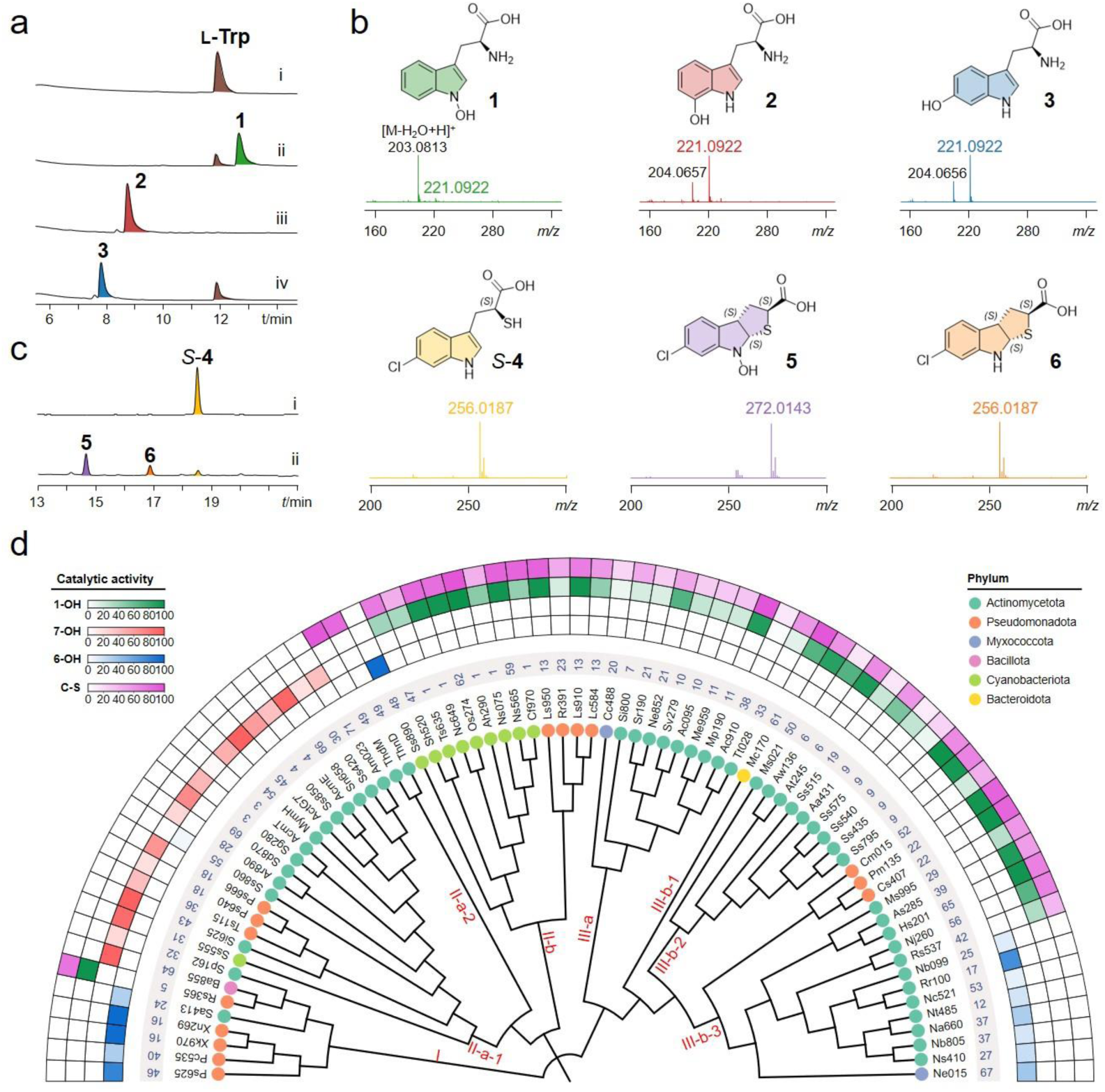
Functional characterization and catalytic diversity of cytochrome P422s. **a**, HPLC profiles of the cytochrome P422 members. (i) _L_-Trp standard; (ii to iv) reaction mixtures containing different cytochrome P422 members (ii, Ba855; iii, Ss890; iv, Mc170), Trp, *Sel*Fdx1499, *Sel*FdR0978, NADPH, and sodium ascorbate. **b**, Positive-ion HR-MS profiles of products **1** to **6**. **c**, HPLC profile of ThnD *in vitro*. (i) The standard for *S*-**4**; (ii) ThnD was incubated with *S*-**4**, *Sel*Fdx1499, *Sel*FdR0978, NADPH, and DTT. **d**, Functional diversification analysis across the cytochrome P422 phylogeny. The Neighbor-joining phylogenetic tree was constructed with the conserved substrate-binding motifs. Colors of the phylogenetic tree terminal nodes correspond to the phylum of each cytochrome P422 member. The blue number specifies the corresponding cluster in the SSN. Colors of heatmap represent the different reaction types: green, N1-hydroxylation (1-OH); red, C7-hydroxylation (7-OH); blue, C6-hydroxylation (6-OH); and purple, C–S bond-formation (C–S). Heatmap shows the *in vitro* catalytic efficiency of selected cytochrome P422 enzymes toward _L_-Trp or *S*-**4**.

For structural characterization, compound **1** was isolated from a large-scale enzymatic reaction of Mc170. ^1^H NMR analysis revealed the disappearance of the indole N–H proton signal relative to L-Trp, indicating hydroxylation at the indole nitrogen (Supplementary Figs. 7–8). To unambiguously assign the site of modification, ^15^N-labeled compound **1** was prepared from ^15^N-labeled L-Trp. Its ^15^N NMR spectrum displayed a characteristic downfield shift (*δ* = 131.98 to 176.43), consistent with N–OH formation and confirming hydroxylation specifically at the N1 position, thereby establishing the structure as 1-hydroxytryptophan (1-OH-Trp, **1**) (Fig. 3a–c, Extended Data Fig. 4a, Supplementary Figs. 8–14 and Supplementary Table 7). To our knowledge, this work reports the first enzymes known to catalyze N1-hydroxylation of L-Trp. Furthermore, 28 of these L-Trp N-hydroxylases also accepted L-abrine, yielding the corresponding N1-hydroxylated product (Extended Data Fig. 3b–d).

In this study, 18 cytochrome P422 enzymes (e.g., Ba855 from *Bacillus atrophaeus*) mediated the formation of new compound **2** with an *m*/*z* value of 221.0922 ([M+H]^+^), consistent with hydroxytryptophan (Fig. 3a-c). Isolation and structural characterization of the hydroxytryptophan from a culture of *E. coli* expressing Ba855 elucidated its structure as 7-hydroxytryptophan (7-OH-Trp, **2**) by HR-LCMS and NMR analysis (Fig. 3a-c, Supplementary Figs. 15-20 and Supplementary Table 8). To the best of our knowledge, enzymes that form 7-OH-Trp were previously unknown until this finding of a cytochrome P422 with 7-hydroxylase activity. This enzyme also presents an alternative pathway to produce 7-hydroxy-*N*-formylkynurenine (7-OH-NFK) through incorporation by TDO, which bypasses the classic kynurenine monooxygenase (KMO) route to enter the kynurenine pathway^2^ (Extended Data Fig. 5a).

It should be mentioned that, although a genetic inactivation mutant of *acnF* (homolog of *acmE*) was not able to produce actinomycin D in a previous study, the timing of the 7-OH installation on the Trp subunit of actinomycin D remains to be determined^22^. Our *in vitro* investigation of AcmE showed that this enzyme catalyzes the regioselective 7-hydroxylation on the free Trp substrate (Extended Data Fig. 2a). Our work also elucidates the uncharacterized 7-hydroxylation step in tasikamide biosynthesis through the identification of Sd870, a Tsk8 homolog (86% identity) located in a highly conserved biosynthetic gene cluster (Extended Data Fig. 2b)^31^. Furthermore, we identified MymH as a Trp-7-hydroxylase, suggesting its involvement in the macemycin biosynthetic pathway through the generation of the precursor 3-hydroxyanthranilic acid (Extended Data Fig. 2c). Moreover, 7-hydroxylase activity toward L-abrine and *N*-acetyl- L-Trp was also observed with these 18 enzymes (Extended Data Fig. 3b-d).

Another group that included 15 enzymes (e.g., Ss890 from *Scytonema* sp. HK-05) gave rise to an alternative product whose mass value (*m/z* 221.0922, [M+H]^+^) is also consistent with that of hydroxytryptophan. The product from the Ss890 reaction mixture was determined to be 6-hydroxy- L-Trp (**3**) by LC-HRMS through comparison with commercial standards (Fig. 3a-c and Extended Data Fig. 5b). A recent study demonstrated that the two HBPs SaT6H and AxT6H also act as Trp 6-hydroxylases of the histidine-ligated type^30^. Our results further expand the enzyme diversity of Trp 6-hydroxylases to thiolate-ligated HBPs.

The functional diversity of the cytochrome P422 family further emerged from investigation of ThnD and ThdM; these two enzymes were first reported to be encoded within the biosynthetic gene clusters *thn* and *thd*, respectively, which are colocalized with the Trp halogenase responsible for producing the sulfur-containing thienodolin^26,32^ (Extended Data Fig. 2d). To investigate the function of ThnD/ThdM, we reconstituted the unknown sulfur incorporation step involved in the formation of the key intermediate *S*-**4** using an *in vitro* “one-pot” incubation with sulfur carrier protein ThnL, deubiquitinase-like sulfurtransferase ThnF, MoeZ-like activating enzyme ThnM, short-chain dehydrogenase ThnE, and L-6-chlorotryptophan^33^ (Extended Data Fig. 6, Supplementary Figs. 21-26 and Supplementary Table 9). As expected, the subsequent incubation of *S*-**4** with cytochrome P422 ThnD/ThdM and redox partners predominantly yielded 6-Cl-4H-HTHA (**5**) (Fig. 3d-f), thereby establishing the enzymatic basis for C-S bond formation in thienodolin biosynthesis. Structural characterization by HRMS and NMR confirmed the thieno[2,3-*b*]indole scaffold (TNI) of **5** and identified the minor *N*-dehydroxylated product 6-Cl-4H-THA (**6**) (Fig. 3d, Supplementary Figs. 27-34 and Supplementary Table 10).

Sulfur-containing heterocycles represent privileged structural motifs in natural products, where C–S bonds play indispensable roles in the bioactivity of numerous pharmaceuticals and biologically active molecules^34^. The development of mild and general methods for C–S bond formation has therefore attracted significant attention, with enzymatic strategies offering particularly promising routes under biocompatible conditions^35^. Known heme-dependent enzymes catalyzing intramolecular C–S bond formation are only found with cytochrome P450s^34,36^. Beyond these established systems, we report that the cytochrome P422 enzyme family represents a newly discovered and distinct class of HBPs capable of catalyzing intramolecular C–S bond formation, expanding the biocatalytic toolbox for sulfur heterocycle assembly and underscoring the diversity of enzymatic strategies for C–S bond construction in natural product pathways.

We further determined the steady-state kinetic constants of representative cytochrome P422 enzymes across different reaction types using L-Trp as the substrate. Mc170 (N1-hydroxylation) exhibited a *K*_m_ of 56.5 μM with a *k*_cat_*/K*_m_ of 1.5 × 10^3^ M^−1^ s^−1^; Ba855 (C7-hydroxylation) showed a *K*_m_ of 90.0 μM and a *k*_cat_*/K*_m_ of 1.3 × 10^4^ M^−1^ s^−1^; and Ss890 (C6-hydroxylation) had a *K*_m_ of 69.9 μM and a *k*_cat_*/K*_m_ of 1.5 × 10^3^ M^−1^ s^−1^ (Extended Data Fig. 7). Separately, ThnD, which catalyzes C–S bond formation, displayed a *K*_m_ of 11.1 μM for substrate *S*-**4**, with a catalytic efficiency *k*_cat_*/K*_m_ of 8.0 × 10^3^ M^−1^ s^−1^ (Extended Data Fig. 7). Collectively, these comprehensive analyses establish the cytochrome P422 family as a class of versatile biocatalysts capable of mediating diverse and unprecedented transformations of Trp, including intramolecular C–S bond formation, and regiospecific N1, C6, and C7 hydroxylations.

### 4 Crystal structure of Mc170 and catalytic mechanisms of cytochrome P422

In order to help elucidating the catalytic mechanism of the cytochrome P422 family, we tried to determine the crystal structures of several representative members, including Mc170, Ba855, Ss890, and ThnD. The crystal structure of Mc170, representative of an N1-hydroxylation enzyme in complex with Trp, was solved at 2.5 Å resolution and revealed key architectural and cofactor-binding features relevant to catalysis (Supplementary Table 11). The asymmetric unit contains one molecule that folds into a compact α/β structure comprising seven α-helices (α1-α7) and four β-strands (β1-β4), with a disordered loop spanning residues 153-165 (Fig. 4a and Extended Data Fig. 8). The overall topology adopts a sandwich-like arrangement where peripheral α-helices encapsulate the central four-stranded β-sheet (Fig. 4a). The disordered loop is proposed to function as a dynamic lid regulating substrate access to the hydrophobic active site (Extended Data Fig. 8).

**Fig. 4.**
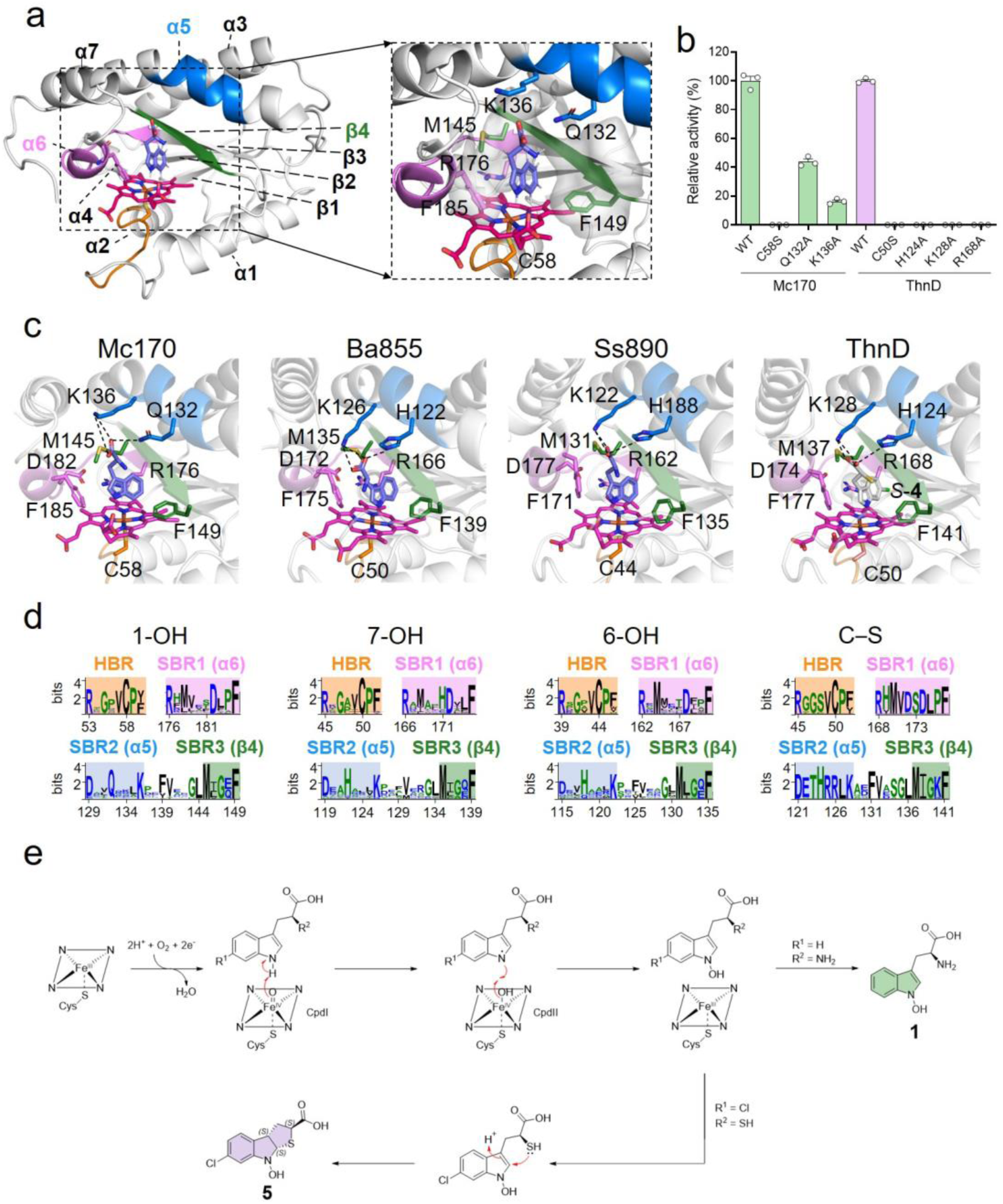
Structural insights into the function and catalytic mechanisms of cytochrome P422 enzymes. **a,** Overall structure of the Mc170-Trp complex. **b,** Relative activity of Mc170 mutants. **c**, Substrate binding modes of different cytochrome P422 members. **d**, Conserved heme-binding motif (HBR) and substrate-binding regions (SBR) of different cytochrome P422 members. **e,** Proposed catalytic mechanisms of N1-hydroxylation by Mc170 and C–S formation by ThnD.

The substrate-binding cavity is delineated by helices α3-α7, β-strands β3-β4, and their flanking loop regions. The structural core involved in heme coordination is formed by helices α1, α2, α4, and α6 together with β-strands β3 and β4. Within the active site, the heme group is nestled between the central β-sheet and two underlying helices, displaying well-defined electron density (Extended Data Fig. 9). Cys-58 serves as the axial fifth ligand to the heme iron. The two heme propionate groups adopt distinct conformations: propionate-6 extends above the heme plane, while propionate-7 lies below the plane and interacts with Arg-53 (Extended Data Fig. 9).

Electron density clearly observed above the heme iron was unambiguously modeled as a L-Trp molecule. The substrate binds in an orthogonal orientation relative to the heme plane, with its carboxylate and indole groups serving as anchoring points at opposite ends (Fig. 4a). The indole ring engages in hydrophobic interactions with Met-145 and Phe-149 on the β4 strand. Helix α5 contributes Gln-132 and Lys-136, which form a hydrogen bond and a salt bridge, respectively, with the carboxylate group of Trp. Furthermore, Arg-176, located on β-strand β3, participates in a stabilizing π-cation interaction with the indole moiety (Fig. 4a). Collectively, Arg-176 (β3), Met-145 (β4), and Phe-149 (β4) form a structural “clamp” that precisely positions the indole ring of the substrate. The measured distance of 4.6 Å between the heme iron center and the N1 atom of Trp is consistent with potential N1-hydroxylation. Pairwise structural alignments with known heme-enzymes in Trp metabolism (e.g., TDOs, P450s, HDAOs) reveal Mc170 has a markedly different fold and active site (RMSD >5 Å), further supporting its classification as a structurally unique HBP (Supplementary Figs. 35 and 36).

Complex structures of Ss890/L-Trp, Ba855/L-Trp, and ThnD/*S*-**4** were predicted using AlphaFold3^37^ (Fig. 4c and Supplementary Fig. 37). Sequence and structural alignments identified a conserved cysteine within the RxGxVCP (heme-binding region, HBR) motif that coordinates the heme cofactor (Fig. 4c, d and Supplementary Fig. 38). The substrate-binding pocket comprises three conserved motifs: Rx(M/L)x_3_Dx_2_F (substrate-binding region 1, SBR1), formed by the loop between β2 and helix α6 and helix α6 itself; Dx_2_(Q/H)x_3_K (SBR2) located in helix α5; and MxGxF (SBR3) formed by β-strand β4 (Fig. 4a, c and d). Within SBR1, the Arg residue participates in a π-cation interaction with the indole ring of Trp, and its protonated guanidinium group forms bidentate hydrogen bonds with the deprotonated carboxylate group of the Asp residue, constituting a stabilizing salt bridge. Within SBR2, the Lys and Gln/His residues anchor the Trp carboxyl group through hydrogen bonding and a salt bridge interaction, respectively, while the Met and Phe residues of SBR3 engage the substrate via hydrophobic interactions.

These analyses indicate a conserved binding mode for Trp/Trp-derivatives in the cytochrome P422 family: the Lys and Gln/His residues anchor the substrate’s carboxyl group, positioning the indole ring near the heme center. Meanwhile, the Arg residue of SBR1, Met and Phe residues of SBR2 form a “clamp” that fixes the indole ring nearly orthogonal to the heme plane. This binding arrangement positions the N1/C6/C7 sites of the indole ring near the heme catalytic center to facilitate their hydroxylation. Further analysis highlighted regio-specificity conferred by non-conserved residues within SBR1. These residue variations serve as a molecular gate, inducing a subtle conformational change in SBR1 in which the Phe side chain swings to a near-orthogonal orientation relative to the indole ring; this relocation repositions the binding mode of substrate Trp relative to the heme plane, thereby controlling the regioselectivity of cytochrome P422 hydroxylation (Extended Data Fig. 10).

To functionally validate these structural observations, we generated the site-directed mutants of Mc170 (C58S, Q132A, K136A, and R176A) and ThnD (C50S, H124A, K128A, and R168A). Both the Mc170^C58S^ and ThnD^C50S^ variants completely lost catalytic activity, and UV–visible spectroscopy confirmed severely disrupted heme binding (Fig. 4b; Supplementary Fig. 39), indicating that the conserved Cys in the HBR is essential for prosthetic group coordination and structural integrity. Attempted expression of the Mc170^R176A^ variant resulted in the formation of inclusion bodies, while the ThnD^R168A^ mutant was successfully purified and exhibited a complete loss of C-S bond formation activity, along with severely impaired heme binding (Supplementary Fig. 40). These results suggest that the Arg residue from SBR1 is also involved in heme binding. The Mc170^Q132A^ and Mc170^K136A^ variants retained partial activity, producing compound **1** at approximately 43% and 16% of wild-type yield, respectively). In contrast, both ThnD^H124A^ and ThnD^K128A^ variants completely lost C-S bond formation activity (Fig. 4b). These results demonstrate that Lys and His/Gln residues within SBR2 play a critical role in anchoring the carboxyl group of the substrate, with their functional importance varying between enzyme subtypes.

Based on integrated structural and mutagenesis data, we propose a catalytic mechanism for Mc170. The reaction initiates when Mc170 accepts two electrons from NAD(P)H via Fdx/FdR to activate molecular oxygen, forming Compound I (CpdI), an oxoiron(IV) species (Fe^IV^=O, i.e. FeO^3+^) coupled with a porphyrin π-cation radical^5,38^. This high-valent intermediate mediates regioselective N1-hydroxylation of the Trp indole ring and converts to the ferric (Feᴵᴵᴵ) state, yielding 1-OH-Trp. A similar mechanistic framework applies to cytochrome P422 enzymes that catalyze C6 or C7-hydroxylation of tryptophan, in which the CpdI facilitates hydroxylation at the C6 or C7 position of the indole ring. The regioselectivity difference between N1-, C6-, and C7-hydroxylation is likely determined by fine-tuning of the substrate orientation within the active site, controlled by key residues lining the enzyme’s binding pocket.

Interestingly, all cytochrome P422 members capable of catalyzing N1-hydroxylation were also found to mediate C–S bond formation when incubated with substrate *S*-**4**. This suggests that after the N1-hydroxylated product is generated, a thio-Michael addition occurs, leading to the formation of the TNI scaffold when a free thiol group is present instead of an amino group (Fig. 4e). In contrast, none of the enzymes (e.g. ThnD, ThdM) catalyzing C–S bond formation were able to hydroxylate tryptophan, highlighting their strict substrate specificity for sulfur-containing compounds.

Phylogenetic reconstruction based on conserved protein regions revealed strong functional segregation within the evolutionary topology of the cytochrome P422 family, demonstrating a clear correlation between phylogenetic clades and reaction types. The family comprises three major clades with distinct catalytic specialization and phylogenetic distribution (Fig. 3g). Clade I consists primarily of *Pseudomonadota* representatives that mainly catalyze Trp C6-hydroxylation; their consistent genomic association with tryptophanase gene clusters suggests a role in synthesizing aromatic compound precursors^39^ (Supplementary Fig. 4). Members with N1 and C7-hydroxylation activity are also revealed within Clade I.

Clade II exhibits functional diversification across three subclades: II-a-1 (predominantly *Actinomycetota*) mediates Trp C7-hydroxylation. Genes for subclade II-a-1 members are frequently encoded within NRPS or TDO-containing gene clusters, potentially supplying 7-OH-Trp as NRPS extender units; notably, 7-OH-Trp can be converted to 7-OH-NFK via TDO to bypass KMO (Extended Data Fig. 5a). Genes for II-a-2 (mostly *Actinomycetota*) reside exclusively in the thienodolin biosynthetic gene cluster, uniquely catalyzing sequential *N*-hydroxylation and C–S bond formation. Subclade II-b members (mainly *Cyanobacteriota* and *Pseudomonadota*) catalyze N1-hydroxylation and are predominantly encoded within NRPS gene clusters, suggesting the widespread existence of undiscovered N1-hydroxylation Trp-containing non-ribosomal peptides (Supplementary Fig. 4).

Clade III members, primarily composed of *Actinomycetota* representatives, are also encoded within NRPS biosynthetic gene clusters, consistent with roles in natural product biosynthesis (Supplementary Fig. 4). Clade III is divided into four functional subgroups: members of III-a, III-b-1, and III-b-2 mediate Trp N1-hydroxylation, while III-b-3 is composed of members that catalyze Trp C6-hydroxylation and others dedicated to N1-hydroxylation. Interestingly, no Trp C7-hydroxylation enzyme was found within this clade, potentially due to functional divergence or limited sampling.

## Discussion

Trp metabolism is involved in multiple physiological processes and biosynthesis of bioactive compounds, yet a significant fraction of HBPs involved in Trp modification remain uncharacterized. To address this gap, we developed CISSspector, an artificial intelligence-driven workflow that integrates deep learning-based structural prediction with genome mining and genomic neighborhood analysis for systematic identification of novel HBPs. Application of this tool to underexplored deep-sea actinomycetes led to the discovery of a group of cysteine-ligated HBPs now designated as the cytochrome P422 family (formerly annotated as the DUF6875 family).

Cytochrome P422 enzymes exhibit exceptional catalytic versatility from a conserved structural scaffold, enabling diverse chemical transformations through precise substrate positioning within a shared active site. Structural elucidation of the Trp N1-hydroxylation enzyme Mc170 at 2.5 Å resolution revealed a unique protein fold unprecedented among known HBPs. Biochemical analyses reveal that reaction specificity is governed by fine-tuning of the conserved substrate-binding pocket, where variations in key residues direct the tryptophan substrate toward distinct outcomes.

The substrate-binding pocket composed of SBR1 and SBR2 motifs anchors the carboxylate and indole groups of tryptophan across all cytochrome P422 family members. Functional specialization arises from subtle yet consequential modifications: in SBR2, a Gln-to-His substitution at the carboxylate-anchoring site distinguishes Trp N1-hydroxylation enzymes from those catalyzing C–S bond formation or hydroxylation of *S*-**4**, potentially reorienting the substrate to expose different reactive centers (N1, C6, or C7) to the heme iron. Additionally, a conserved phenylalanine in SBR1 acts as a conformational switch; its side-chain rotamer reorients the indole ring to govern hydroxylation regioselectivity.

The site-selective C–H hydroxylation of the ubiquitous Trp scaffold remains a synthetic challenge, typically requiring directing groups to achieve regiocontrol by chemical synthesis^40^. Here, we present a biocatalytic solution through the discovery of cytochrome P422 enzymes that directly mediate Trp hydroxylation at the N1, C6, and C7 positions. This unprecedented enzymatic capability addresses the long-standing absence of dedicated N1- and C7-hydroxylases. Together with known Trp-modifying enzymes, the P422 family now enables nearly complete regio-hydroxylation of the indole ring, providing powerful new biocatalytic tools for synthetic biology and metabolic engineering.

Notably, significant catalytic overlap among cytochrome P422 enzymes underscores a unified mechanistic foundation. The capacity of Trp N1-hydroxylation enzymes to also facilitate C–S bond formation points to a shared initiation step involving N1-hydroxylation. The ultimate reaction trajectory is then determined by substrate availability and active site microenvironments. In this paradigm, a common high-valent iron-oxo intermediate (CpdI) is selectively channeled into divergent pathways through exquisite control of substrate positioning—establishing the cytochrome P422 family as a novel example of divergent evolution within a conserved structural framework.

Our findings establish the cytochrome P422 family as a structurally and functionally unique class of HBPs critical to specialized metabolism. Beyond expanding the catalytic repertoire of Trp metabolism, this work demonstrates the power of CISSspector as a scalable framework for discovering cryptic enzymatic functions in underexplored proteomes. The strategy is readily adaptable to other cofactor-dependent enzymes, opening new avenues for natural product discovery, protein engineering, and synthetic biology applications.

## Methods

### Inference optimization

We chose Boltz-1 to predict complex structures because Boltz-1 has achieved predictive accuracy comparable to AlphaFold3^37^ while offering computational efficiency more suitable for large-scale prediction. In addition, to accelerate the prediction process, multiple sequence alignments (MSAs) were generated locally by invoking MMseqs2^41^ via ColabFold^42^, which improved computational efficiency by more than fourfold compared to the use of remote MSA servers.

### The preparation of a database for model training and validation

The dataset preparation process is shown in Extended Data Fig. 1a. A total of 571,521 protein sequences were retrieved from the UniProtKB Reviewed database (version 2025.1.25) and divided into two subsets according to the presence of the keyword “heme binding”. For the keyword-containing subset (13,007 sequences), single-chain entries within a reasonable length range (80–800 residues) were filtered and redundancy was removed, yielding 11,083 sequences. Random sampling revealed a significant proportion of non-heme-binding proteins due to annotation errors; therefore, stratified sampling and manual curation were performed.

Based on Boltz-1 confidence scores, 200 sequences were randomly sampled from the critical range of high uncertainty (both complex_iplddt and iPTM between 0.5–0.8) for manual curation, along with another 200 sequences sampled from non-critical range, ensuring an approximate 1:1 ratio of positive and negative cases. These formed dataset A2 for training and validation. To prevent data leakage, the entire data set is clustered with a similarity of 0.5. The results exclude all categories belonging to dataset A2, and then randomly sample 100 from the remaining categories to form dataset A1. Similarly, the keyword-absent subset (558,244 sequences) was deduplicated, length-filtered and clustered with a similarity of 0.5, followed by a random sampling of 1000 sequences (nearly 1%), generating dataset B specifically for testing model performance on negative cases.

### Training features selection

Training features were constructed from both confidence scores and structural features. Confidence scores from Boltz-1 provide critical indicators of structural reliability and have been also employed in Boltz-Design^11^ for sequence generation through backpropagation. Given that the study targets intermolecular binding, complex_iplddt and iPTM were selected, as they evaluate interface-specific structural confidence and global structural accuracy, respectively.

Structural features included: (i) structural rationality (absence of unreasonable atomic distances), (ii) coordination rationality (presence of coordinating residues), (iii) the number of residues surrounding the heme (reflecting burial depth), and (iv) the number of coordinating histidines, the critical residue type that were found to coordinate with HEME in 83% of HEME-binding proteins^43^. The first two features were applied for direct filtering. The latter two were quantified using a structure-analysis script across all predicted structures, which were combined with complex_iplddt and iPTM to form the final feature set for model training.

### Training details

Normalization of sample features for LR^13^ and NB^14^ was performed to eliminate the effect of dimensional differences, and RF^15^ that is not sensitive to the numerical range still used the original features as input. To avoid overfitting RF, the value ranges of its key parameters were limited and grid search was used to find the optimal parameter combination. For direct threshold screening, the complex_iplddt, iPTM and surrounding residue number with a large range of values were used as the screening thresholds to find the optimal threshold combination using grid search. The settings of key parameters of all models were shown in Supplementary Table 1.

### Post-processing of predictions

Incorporating prior knowledge of key features into the post-processing of predictions can compensate for limitations in feature coverage and generalization, while also introducing biological constraints that purely data-driven models may overlook^44^. First, coordination and structural validity were assessed using outputs from the structure-analysis script, retaining only those samples with at least one plausible coordinating residue (H, C, M, Y, K)^45^ and no sterically unreasonable atomic distances (≤2.0 Å). Furthermore, feature importance analysis from the LR model revealed that complex_iplddt and surrounding residue number exerted dominant influence (with coefficient 3.78 and 2.20, respectively), identifying them as the most critical features for classification. So complex_iplddt was set to a threshold of no less than 0.7, following the official recommendation for pLDDT to ensure structural reliability^46^. For surrounding residue number, the threshold was determined as 27, based on the mean (30.27) and standard deviation (4.10) of all positive samples from the total curated set of 500 sequences, thereby ensuring sufficient heme embedding depth. Applying these four post-processing filters enabled the extraction of high-quality candidates from the prediction pool that may effectively reduce false positives and enhance precision.

The 44 hypothetical HBPs were subjected to sequence similarity network (SSN) analysis using the online EFI-Enzyme Similarity Tool (EFI-EST)^19,20^, with clustering performed at a 25% sequence identity threshold. All results are referenced in Supplementary Table 3 and Supplementary Fig. 2.

### The preparation of a database of 100 *Actinomycetes* strains for HBPs screening

Draft genomes of 100 *Actinobacteria* strains were sequenced, and their coding sequences (CDSs) were predicted using GeneMarkS-2^47^, resulting in a total of 531,004 predicted protein sequences. All sequences were functionally annotated using eggNOG-mapper^48^. From the annotation results, 100,211 sequences that were either unannotated or annotated as domains of unknown function (DUF) in the Pfam database were selected. These sequences were further filtered by length (80–800 amino acid residues) and then clustered at 90% sequence identity using MMseqs2^41^, yielding a non-redundant dataset of 31,193 sequences. This final dataset was subsequently used for HBPs screening with CISSpector.

### Sequence-only models evaluating

To compare classification performance, the model CLEAN^7^, which only relies on amino acid sequences to predict EC numbers, was used to predict the EC numbers of four representative sequences from the cytochrome P422 family. The catalytic functions corresponding to the predicted EC numbers showed no correlation with the actual catalytic functions and do not require heme involvement. Additionally, we did not succeed in deploying the DeepECtransformer^8^, so its classification performance will not be evaluated (Supplementary Table 4).

### General analysis procedures

HPLC analyses were performed using a YMC-Triart C18 (250 × 4.6 mm, 5 μm, YMC, Japan) column with a biphasic solvent system of acetonitrile (ACN) and water (containing 0.1% formic acid, FA) on a Thermo Scientific Vanquish HPLC system. All compounds were isolated using a YMC-Pack Pro C18 column (250 × 10 mm × 5 μm, YMC, Japan). HPLC-HRMS analyses were carried out on a Bruker impact HD High Resolution Q-TOF mass spectrometry. HPLC-ESI-MS spectra for proteins were obtained using a Thermo High Resolution LTQ-Orbitrap mass spectrometer with ETD. NMR spectra were acquired on a Bruker Avance III 600 MHz spectrometer (Switzerland). The UV-visible spectra were recorded on a X-7 UV-vis spectrophotometer (Shanghai Metash Instruments Co., Ltd.). CD spectra were collected on a JASCO J-815 CD spectrometer. The concentration of purified protein was determined by measuring its absorption at 280 nm using an Ultramicro-UV-vis spectrophotometer ND-100C (Hangzhou Miu Instruments Co., Ltd.). “ÄKTA pure” or “ÄKTA prime plus” fast protein liquid chromatography machines were used to perform protein purification.

### Protein expression and purification

A single colony of recombinant *E. coli* BL21(DE3) harboring the target protein expression vector was inoculated into 5 mL of LB medium supplemented with 50 mg/L kanamycin. The culture was incubated at 37 ℃ with shaking at 220 rpm for 16 h. This seed culture was then transferred into 500 mL of fresh LB medium containing 50 mg/L kanamycin and grown under the same conditions until the OD_600_ reached 0.6–0.8 (approximately 3–5 h). Protein expression was induced by adding isopropyl-β-D-thiogalactopyranoside (IPTG) to a final concentration of 0.2 mM. To enhance heme biosynthesis, 0.2 mM 5-aminolevulinic acid (5-ALA) was also added. The induced culture was incubated at 18 ℃ for 18–20 h with shaking. Cells were harvested by centrifugation at 6,000 × *g* for 10 min at 4 °C, and the pellets were stored at –80℃. For purification, frozen cell pellets were thawed and resuspended in 50 mL of lysis buffer (50 mM NaH_2_PO_4_, 300 mM NaCl, 10 mM imidazole, 10% glycerol, pH 7.4). Cell disruption was performed by sonication using a Model 500 Sonic Dismembrator. The lysate was clarified by centrifugation at 10,000 × *g* for 60 min at 4 ℃. The supernatant was incubated with 2 mL of Ni-NTA resin (Qiagen, Venlo, Netherlands) for 30 min with gentle mixing. The resin was then washed extensively with wash buffer (50 mM NaH_2_PO_4_, 300 mM NaCl, 20 mM imidazole, 10% glycerol, pH 7.4) until no protein was detected in the flow-through by Coomassie Brilliant Blue G-250 staining. The His_6_-tagged target protein was eluted with elution buffer (50 mM NaH_2_PO_4_, 300 mM NaCl, 250 mM imidazole, 10% glycerol, pH 7.4). The eluate was concentrated using an Amicon Ultra centrifugal filter (appropriate molecular weight cutoff; Merck KGaA, Germany) at 5,000 × *g*. The concentrated sample was buffer-exchanged into desalting buffer (50 mM NaH_2_PO_4_, 100 mM NaCl, 10% glycerol, pH 7.4) using a PD-10 column (GE Healthcare, Buckinghamshire, UK) and concentrated again. Protein concentration To improve the solubility of challenging proteins (Pp145, Bi410, and Sh310), an N-terminal SUMO fusion tag was utilized, which enabled their successful purification.

For crystallization of Mc170, the eluted sample was applied onto size-exclusion chromatography using the ÄKTA pure25 system (GE) mount with Superdex 200 pg 16/600 column and equilibrated by buffer C (20 mM Tris pH 8.0 and 150 mM NaCl) for further purification. Purified proteins were collected and concentrated to 4 mg/mL Amicon Ultra-15 10 K device (Millipore) and stored at -80 ℃.

### Crystallization, data collection and structure determination

Purified proteins were mixed with the L-Trp (100 mM stock in H_2_O) to a 1:5 molar stoichiometry of protein: L-Trp. Crystals were obtained by utilizing the sitting drop vapor diffusion method in a 1:1 ratio with the crystallization condition. The co-complex crystals of Mc170 with L-Trp were grown in the precipitant solution of 15% (w/v) PEG 20,000, 100 mM HEPES/ Sodium hydroxide pH 7.0. Crystals were supplemented with cryoprotectants containing the reservoir contents plus 20% ethylene glycol and flash-frozen in liquid nitrogen. Diffraction data of Mc170 crystals was collected at 100 K on beamline BL18U at the Shanghai Synchrotron Radiation Facility (SSRF) and processed using the HKL3000 program^49^.

### Structure determination and refinement

The complex structure of Mc170 with L-Trp was solved by molecular replacement with the program Phaser^50^, using the AF3 model as the search model. Further manual model building was facilitated by using Coot^51^, combined with the structure refinement using Phenix^52^. Data collection and structure refinement statistics are summarized in Supplementary Table 1. The Ramachandran statistics, as calculated by Molprobity^53^, are 88.36%/5.59% (favored/outliers) for structures of Mc170. All structure figures were prepared in PyMOL (Schrödinger) and ChimeraX^54^.

### UV–Visible spectroscopic characterization of cytochrome P422 enzymes

For preparation of the dithionite-reduced ferrous-CO complex of cytochrome P422 enzymes, carbon monoxide gas was slowly bubbled into a solution of purified ferric enzyme (∼1–20 μM) in 50 mM NaH_2_PO_4_, 300 mM NaCl, 10% glycerol, pH 7.4, immediately followed by sufficient reduction of the protein with sodium dithionite (1–3 mg)^55^. The optical absorption spectra of the ferric and ferrous–CO forms were recorded before and after the addition of sodium dithionite, respectively.

### Inductively coupled plasma mass spectrometry (ICP-MS) assays of ThnD

For inductively coupled plasma mass spectrometry (ICP-MS) analysis, a sample of purified ThnD (typically containing 2.7 mg of protein) was digested with 3 mL of 70% nitric acid (HNO_3_) using a microwave digestion system (Multiwave PRO). The resulting digestate was then analyzed with an Agilent Technologies 7900 ICP-MS, and the data were processed using Agilent MassHunter software.

### Identification of auxiliary proteins in the sulfur transfer system

In our recent identified canonical DUB-like sulfurtransferase systems, sulfur incorporation depends on the coordinated action of a sulfur carrier protein (SCP) and its cognate MoeZ-like activating enzyme^33^. Using Cxm4 and CxmM as probes, we identified their respective homologs in the genome of *Streptomyces* sp. FXJ1.172: ThnL (GenBank: WP_067053588.1), which shares 40% sequence identity with Cxm4, and ThnM (GenBank: WP_067038662.1), which shares 90% sequence identity with CxmM.

### In vitro enzymatic assays and products detection

Unless otherwise specified, all enzymatic assays were performed in a total volume of 100 μL using 50 mM desalting buffer (pH 7.4) at 30 ℃. Heat-inactivated (boiled) enzyme preparations were used as negative controls.

Cytochrome P422 activity screening: Reactions contained 20 μM cytochrome P422 enzyme, 40 μM SelFdx1499, 20 μM SelFdR0978 (redox partners from Synechococcus elongatus PCC 7942), 10 mM sodium ascorbate, 5 mM NADPH, and 0.5 mM substrate. After 4 h at 30 ℃, reactions were terminated for analysis.

Sulfur-transfer assay: Standard reactions contained 1 mM L-6-Cl-Trp, 10 μM ThnJ (GenBank: AMR44310.1), 10 μM ThnF (GenBank: AMR44306.1), 10 μM ThnL, 10 μM ThnM (GenBank: WP 067038662.1), 2 mM ATP, 2 mM Na2S2O3, 5 mM MgCl2, and 10 mM DTT in 100 μL. Incubation proceeded for 4 h at 30 ℃.

ThnE-containing assay: The system included all components of the sulfur-transfer assay plus 10 μM ThnE (GenBank: AMR44305.1) and 10 mM NADPH.

Assays with ThnD or ThdM: Separate reactions were assembled, each containing either 10 μM ThnD or 10 μM ThdM, together with 40 μM SelFdx1499, 20 μM SelFdR0978, 5 mM NADPH, 10 mM DTT, and 0.5 mM substrate (S-4). Incubation was carried out at 30 ℃ for 2 h.

Reaction termination and analysis: All reactions were quenched with 200 μL methanol. Precipitated proteins were removed by centrifugation (12,000 × g, 10 min), and supernatants were analyzed by HPLC or HPLC-HRMS using the following gradient programs (flow rate: 1 mL/min; mobile phase: acetonitrile (ACN)/H2O with 0.1% formic acid):

Compounds 1–3: 5% ACN (0–1 min), 5%→40% ACN (2–25 min), 100% ACN (26–30 min), 5% ACN (31–35 min); detection at 280 nm. Compounds 4–6: 20% ACN (0–1 min), 20%→80% ACN (2–25 min), 100% ACN (26–30 min), 20% ACN (31–35 min); detection at 254 nm.

### Enzymatic synthesis of compound 1, 4, and 5

Compound **1**: A large-scale enzymatic reaction (50 mL total volume) was carried out in desalting buffer containing 1 mM L-Trp, 20 μM Mc170, 40 μM *Sel*FdR0978, 80 μM *Sel*Fdx1499, 10 mM sodium ascorbate, and 5 mM NADPH. After incubation at 30 ℃ for 4 h, the reaction mixture was freeze-dried directly. The residue was dissolved in 20% ACN and centrifuged. The supernatant was purified by semi-preparative HPLC using 15% ACN in water (0.1% FA) at a flow rate of 2.5 mL/min, yielding ∼5 mg of compound **1**.

Compound *S*-**4**: A 100 mL enzymatic reaction was performed in desalting buffer containing 1 mM L-6-Cl-Trp, 10 μM ThnF, 10 μM ThnL, 10 μM ThnM, 2 mM ATP, 2 mM Na_2_S_2_O_3_, 5 mM MgCl_2_, 10 mM dithiothreitol, 10 μM ThnE, and 10 mM NADPH. After 4 h at 30 ℃, the mixture was extracted twice with ethyl acetate (2 × 200 mL). The combined organic phases were concentrated under reduced pressure, redissolved in 5 mL methanol, and centrifuged. The supernatant was purified by semi-preparative HPLC with 50% ACN in water (0.1% FA) at 3 mL/min, affording approximately 10 mg of compound *S*-**4**.

Compound *R*-**4**: A 100 mL enzymatic reaction was set up in desalting buffer containing 1 mM L-6-Cl-Trp, 10 μM ThnF, 10 μM ThnL, 10 μM ThnM, 2 mM ATP, 2 mM Na_2_S_2_O_3_, 5 mM MgCl_2_, 10 mM dithiothreitol, 10 μM Cxm6^33^ (GenBank: AVL27081.1), and 10 mM NADPH. Following incubation at 30 ℃ for 4 h, the mixture was extracted twice with ethyl acetate (2 × 200 mL). The organic phases were combined, concentrated, dissolved in 5 mL methanol, and centrifuged. The supernatant was subjected to semi-preparative HPLC (50% ACN in water with 0.1% FA, 3 mL/min), yielding ∼10 mg of compound *R*-**4**.

Compound **5**: A 50 mL enzymatic reaction containing 0.5 mM *S*-**4**, 20 μM ThnD, 40 μM *Sel*FdR0978, 80 μM *Sel*Fdx1499, 10 mM sodium ascorbate, and 5 mM NADPH in desalting buffer was incubated at 30 ℃ for 2 h. The mixture was freeze-dried, dissolved in 20% ACN, and centrifuged. The supernatant was purified by semi-preparative HPLC using 30% ACN in water (0.1% FA) at 3 mL/min, affording approximately 2 mg of compound **5**.

### Fermentation and purification of compound 2

A single colony of recombinant *E. coli* BL21(DE3) harboring the *ba855* expression vector was inoculated into 5 mL of LB medium supplemented with 50 mg/L kanamycin and cultured at 37 ℃ with shaking at 220 rpm for 16 h. This seed culture was then transferred into 500 mL of fresh LB medium (with 50 mg/L kanamycin) and grown under the same conditions until the OD_600_ reached 0.6–0.8. Protein expression was induced by adding 0.2 mM IPTG and 0.2 mM 5-aminolevulinic acid (5-ALA), after which the culture was incubated at 18 ℃ with shaking at 150 rpm for 15 h. Subsequently, 1 mM L-Trp was added to initiate biotransformation, and the culture was further incubated at 30 ℃ for 24 h.

After fermentation, the cells were extracted three times with an equal volume of ethyl acetate/methanol/glacial acetic acid (80:15:5, *v*/*v*/*v*). The combined organic extracts were dissolved in methanol, centrifuged, and the supernatant was purified by semi-preparative HPLC. Compound **2** was isolated using a gradient of 10% ACN in water (0.1% FA) at a flow rate of 2.5 mL/min.

### The steady-state kinetic analysis of Mc170, Ba855, Ss890, and ThnD

Steady-state kinetic parameters were determined for Mc170, Ba855, Ss890, and ThnD. All reactions were performed in triplicate in 1.5 mL centrifuge tubes containing 200 μL of reaction buffer (50 mM NaH_2_PO_4_, 10% glycerol, pH 7.4) and pre-incubated at 30 ℃ for 5 min. Each reaction mixture contained the respective enzyme, 25 μM *Sel*FdR0978, 50 μM *Sel*Fdx1499, 10 mM sodium ascorbate, and varying concentrations of substrate, as specified below. Reactions were initiated by adding 5 mM NADPH, quenched at 0, 30, and 60 s with 50 μL methanol, and centrifuged at 14,000 × *g* for 10 min. Supernatants were subjected to HPLC analysis to monitor substrate consumption. Kinetic parameters were analyzed using GraphPad Prism 8.3.0. Mc170: Enzyme concentration: 2–10 μM; substrate: L-Trp (50–250 μM). Ba855: Enzyme concentration: 0.25–1.25 μM; substrate: L-Trp (50–250 μM). Ss890: Enzyme concentration: 2–10 μM; substrate: L-Trp (50–250 μM). ThnD: Enzyme concentration: 1–4 μM; substrate: *S*-**4** (10–150 μM); additional component: 5 mM TCEP.

### Phylogenetic Analysis

A multiple sequence alignment of the retrieved homologs was performed using ClustalW^56^ to assess sequence conservation. Catalytically important and substrate-binding residues were identified from the alignment and used for phylogenetic reconstruction (Supplementary Fig. 38). Phylogenetic trees were constructed from the validated enzyme sequences using the Neighbor-joining method in MEGA 7.0^57^ with 1,000 bootstrap replicates, and visualized with tvBOT^58^.

## Acknowledgements

This work was supported by the National Natural Science Foundation of China (U2106227 and 82022066), and the Independent Deployment Project of the Institute of Oceanology, Chinese Academy of Sciences (OCASZZCG009). We also thank the staff of the Institute of Oceanology, Chinese Academy of Sciences and the Core Facilities for Life and Environmental Sciences, State Key Laboratory of Microbial Technology of Shandong University for the HRESI LCMS and NMR analyses.

## Author contributions

W.Z. conceived and designed the study, developed the experimental strategy, analyzed and interpreted the data, and wrote the manuscript. W.M. designed and performed most of the experiments. Q.W. contributed to development and validation of CISSspector. Q.Y. performed the collection of X-ray diffraction data and structural analysis. Z.T. contributed to phylogeny analysis. Q.L., R.W., and R.Z. contributed to purification of protein. P.F. contributed to separation and purification of compounds. W.M., X.H., M.S., J.Z., Y.W., Y.Z., L.J., and W.Z. interpreted the experimental data. W.M., Q.W., Q.Y., F.P.G., and W.Z. revised the manuscript.

---

**Extended Data Fig. 1.**
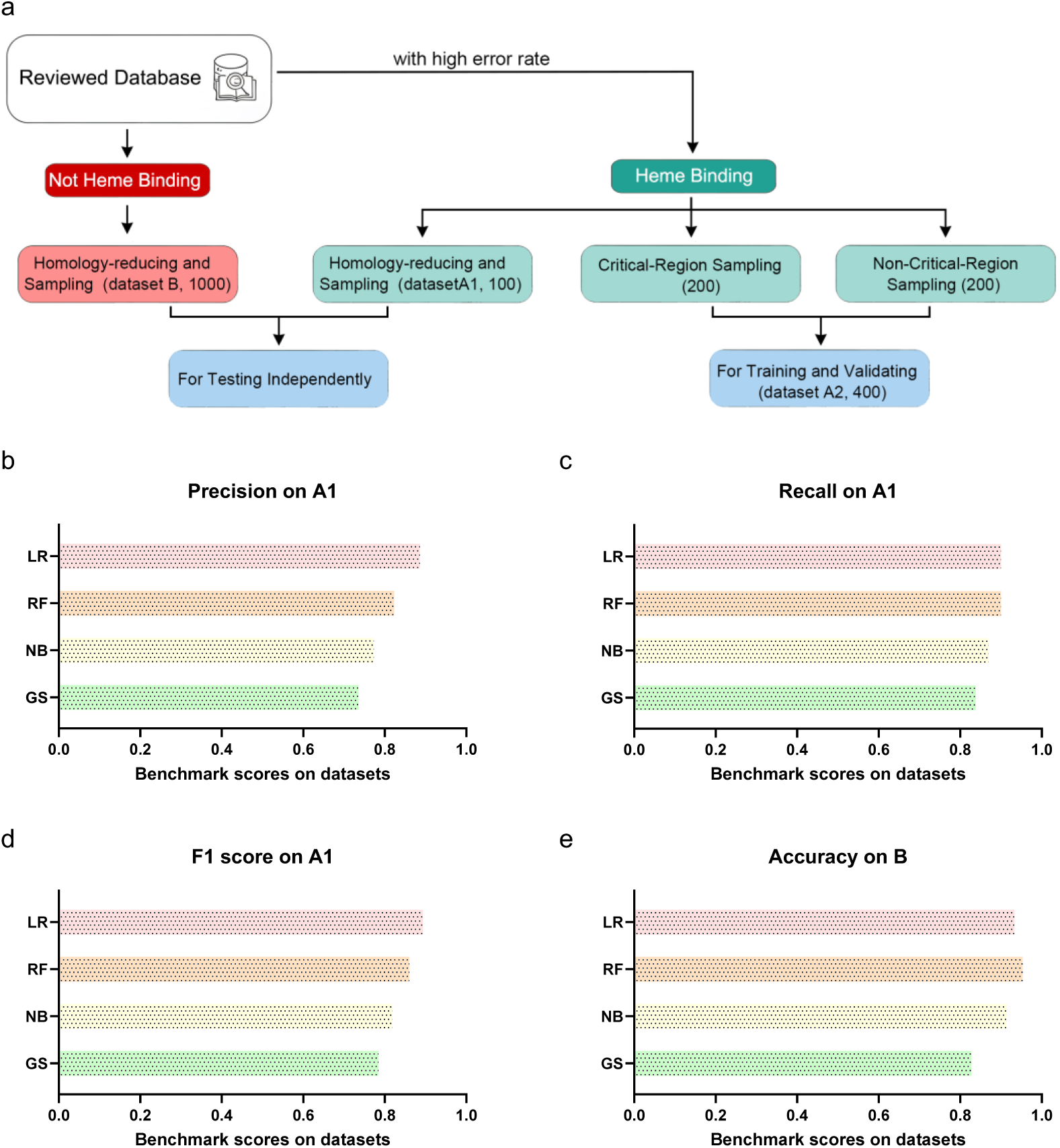
Benchmarking of classification models. **a**, Schematic illustrating the sampling strategy for constructing datasets A1 (training), A2 (validation), and B (independent test). The Precisions (**b**), recalls (**c**), F1 scores on dataset A1 (**d**), and the negative-sample accuracies on dataset B (**e**) of 4 classification algorithms.

**Extended Data Fig. 2.**
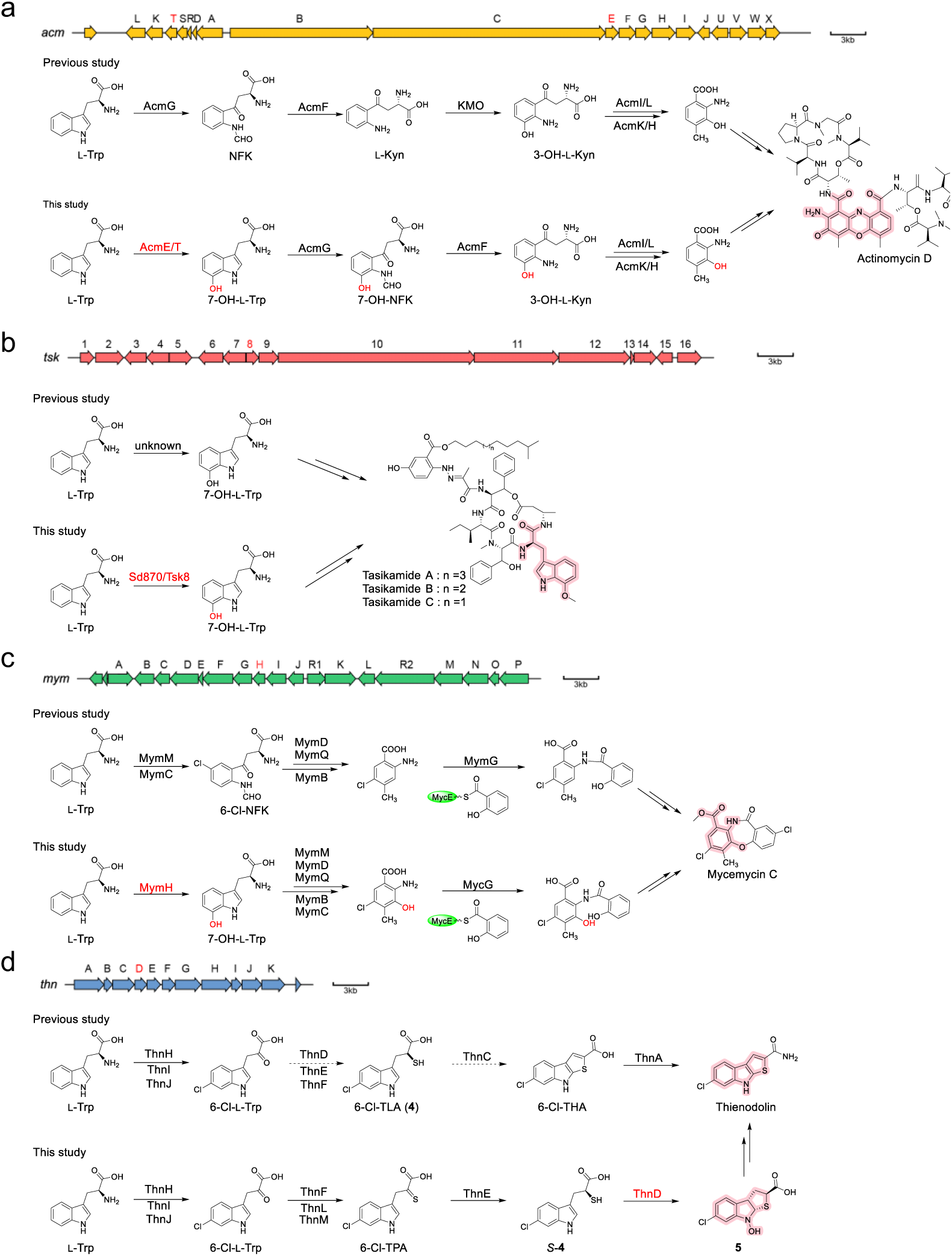
Natural products biosynthesized by cytochrome P422 family-containing gene clusters. Biosynthetic gene cluster and a partial pathway for actinomycin D (**a**), tasikamide (**b**), mycemycin C (**c**) and thienodolin (**d**).

**Extended Data Fig. 3.**
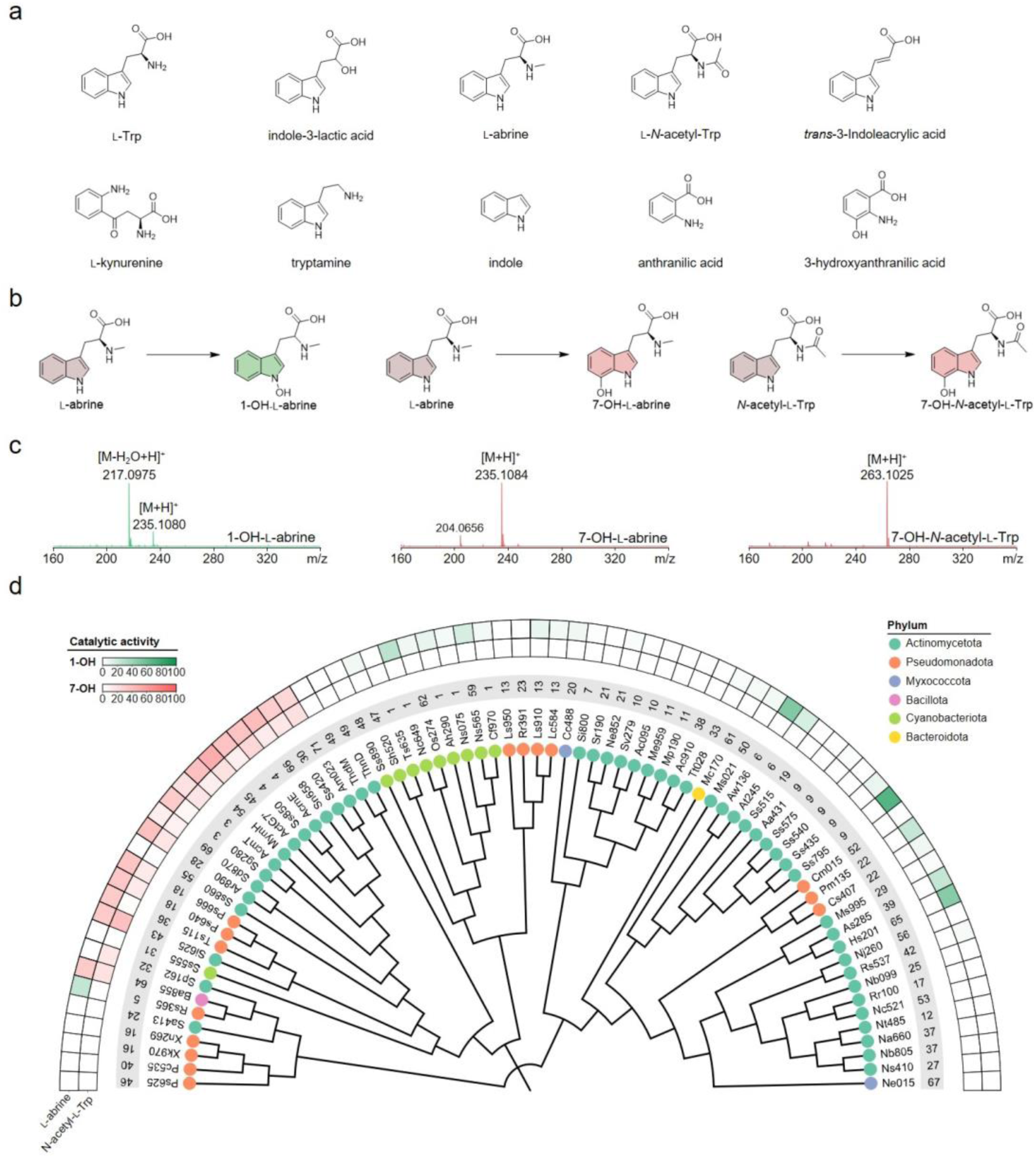
Catalytic diversity and functional evolution of cytochrome P422 enzymes. **a,** Screening substrates: tryptophan-related metabolites and structural analogs. **b**, Representative reaction types catalyzed by cytochrome P422 family members. **c**, Positive-ion HR-MS profiles of products derived from _L_-abrine and N-acetyl-_L_-Trp, including _L_-dehydroabrine, 7-OH-_L_-abrine, and 7-OH-N-acetyl-_L_-Trp. **d**, Substrate scope profiling and phylogenetic-functional correlation analysis of a cytochrome P422 enzyme capable of both N1-hydroxylation and C7-hydroxylation reactions. The Neighbor-joining phylogenetic tree was constructed with the conserved substrate-binding motifs. Colors of the phylogenetic tree terminal nodes correspond to the phylum of each cytochrome P422 member. The gray-background number specifies the corresponding cluster in the SSN. Colors of heatmap represent the different reaction types: green, N1-hydroxylation (1-OH); red, C7-hydroxylation (7-OH). The heatmap displays *in vitro* catalytic efficiencies of selected cytochrome P422 enzymes toward N-acetyl-_L_-Trp and _L_-abrine.

**Extended Data Fig. 4.**
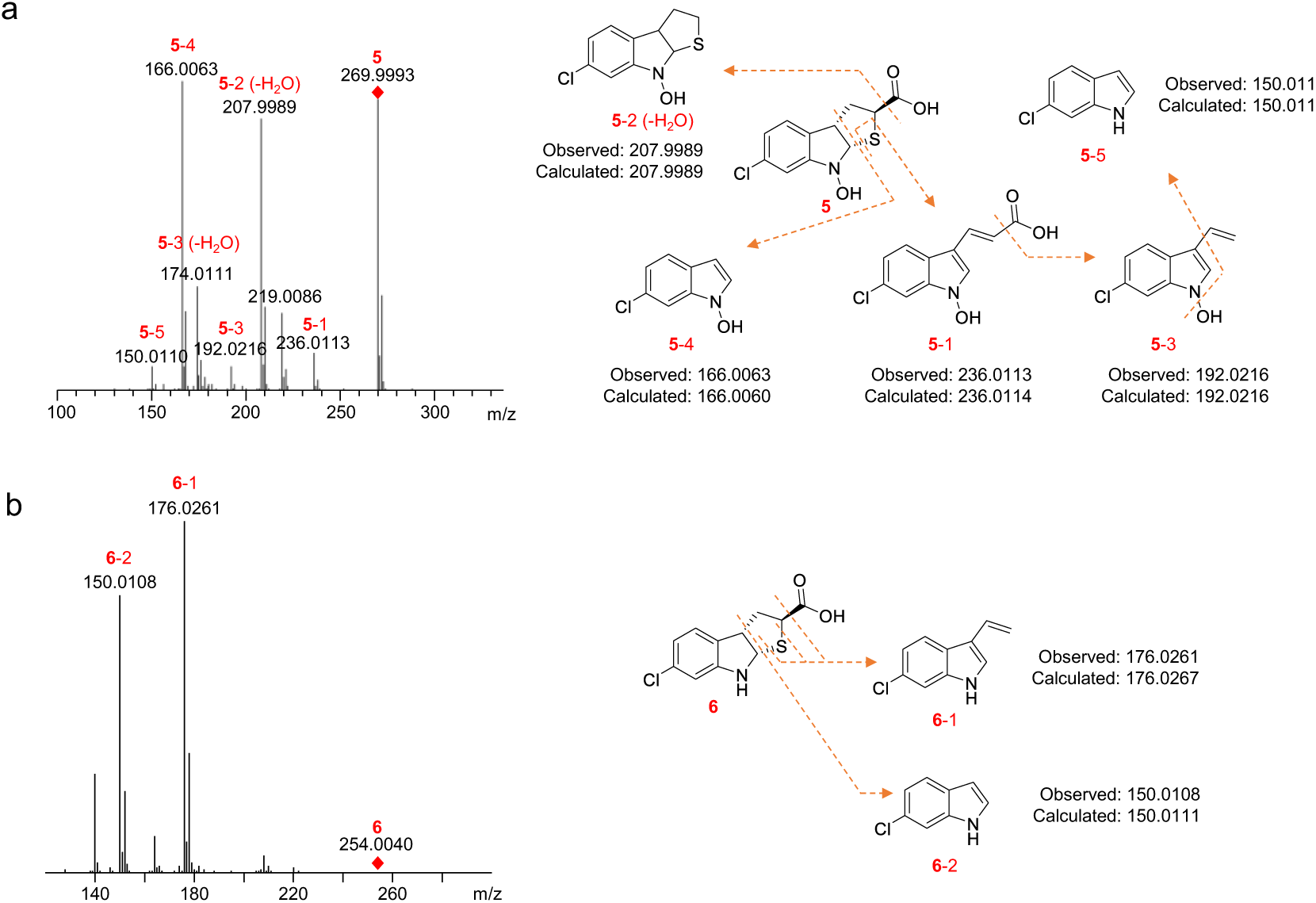
HR-ESI-MS/MS analysis in negative ion mode of compound 5 (a) and compound 6 (b). **a**, The peaks at *m/z* 192.0216 (**5**-3) and 160.0060 (**5**-4) correspond to fragment ions derived from compound **5** through elimination of –SCOOH and –CH_2_CHSCOOH, respectively. **b**, The peaks at *m/z* 176.0261 (**6**-1) and 150.0111 (**6**-2) correspond to fragment ions originating from compound **6** via loss of –SCOOH and – CH_2_CHSCOOH, respectively. These results indicate that compound **6** lacks the *N*-hydroxy group present in compound **5**.

**Extended Data Fig. 5.**
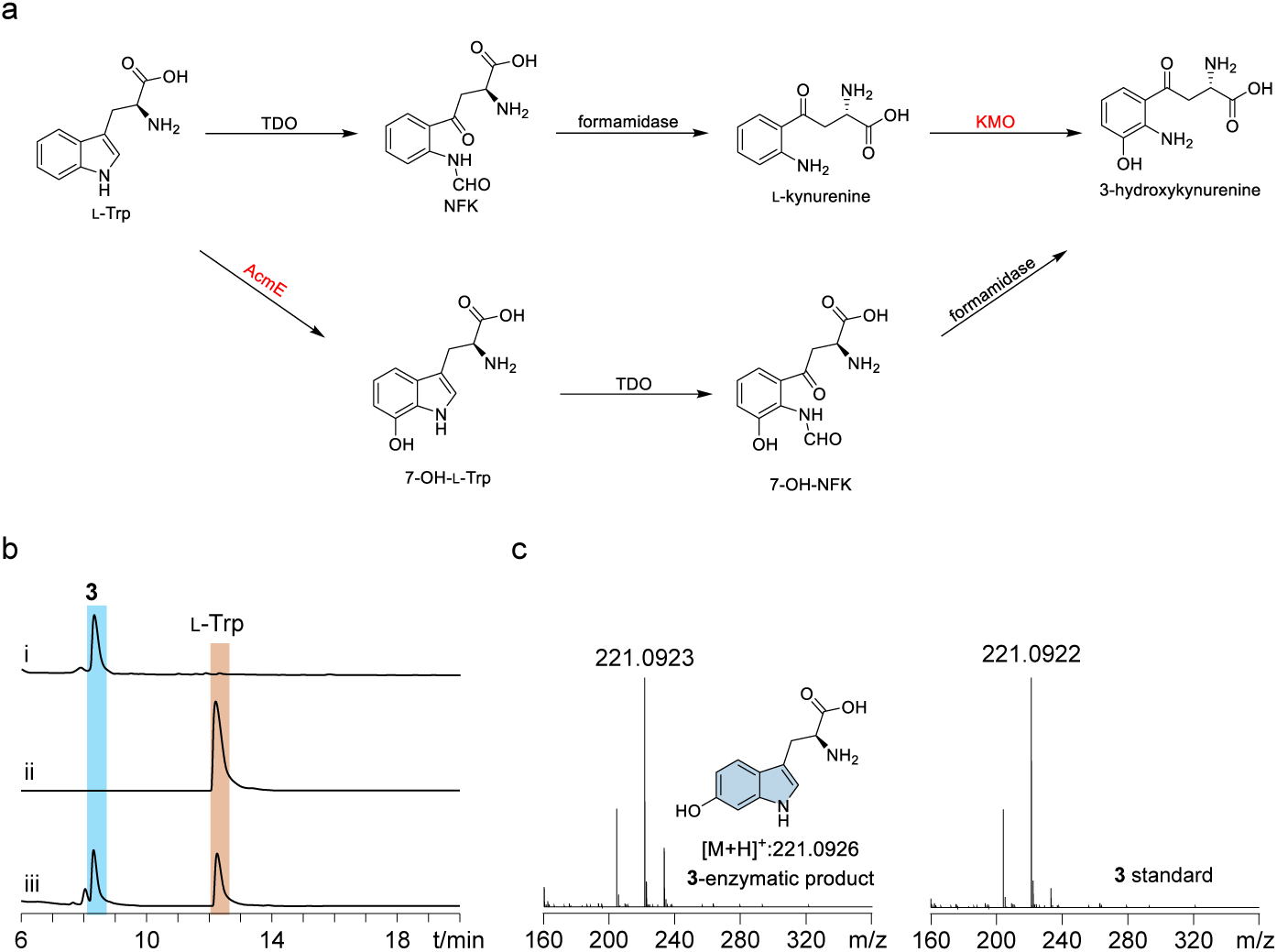
Functional characterization of hydroxylase members in the cytochrome P422 enzyme family. **a**, Biosynthetic pathway of 3-hydroxykynurenine via two distinct routes. Classical route: _L_-Trp is hydroxylated by TDO to form *N*-formylkynurenine (NFK), which is deformylated by kynurenine formamidase to kynurenine, then converted by kynurenine monooxygenase (KMO) into 3-hydroxykynurenine. Alternative route: Cytochrome P422 enzymes (e.g., AcmE) catalyze C7-hydroxylation of _L_-Trp to form 7-OH-Trp, which is oxidized by TDO to 7-OH-NFK. Subsequent deformylation by kynurenine formamidase directly affords 3-hydroxykynurenine, enabling direct entry into kynurenine metabolism. **b**, Analysis of the reaction catalyzed by Ss890. (i) Standard of **3**; (ii–iii) Reaction mixtures containing _L_-Trp (0.5 mM), *Sel*Fdx1499 (100 μM), *Sel*FdR0978 (50 μM), NADPH (10 mM), and sodium ascorbate (10 mM), incubated with either (ii) heat-inactivated Ss890 (10 μM) or (iii) active Ss890 (10 μM). **c**, HRMS profiles of **3** compared with its commercial standard.

**Extended Data Fig. 6.**
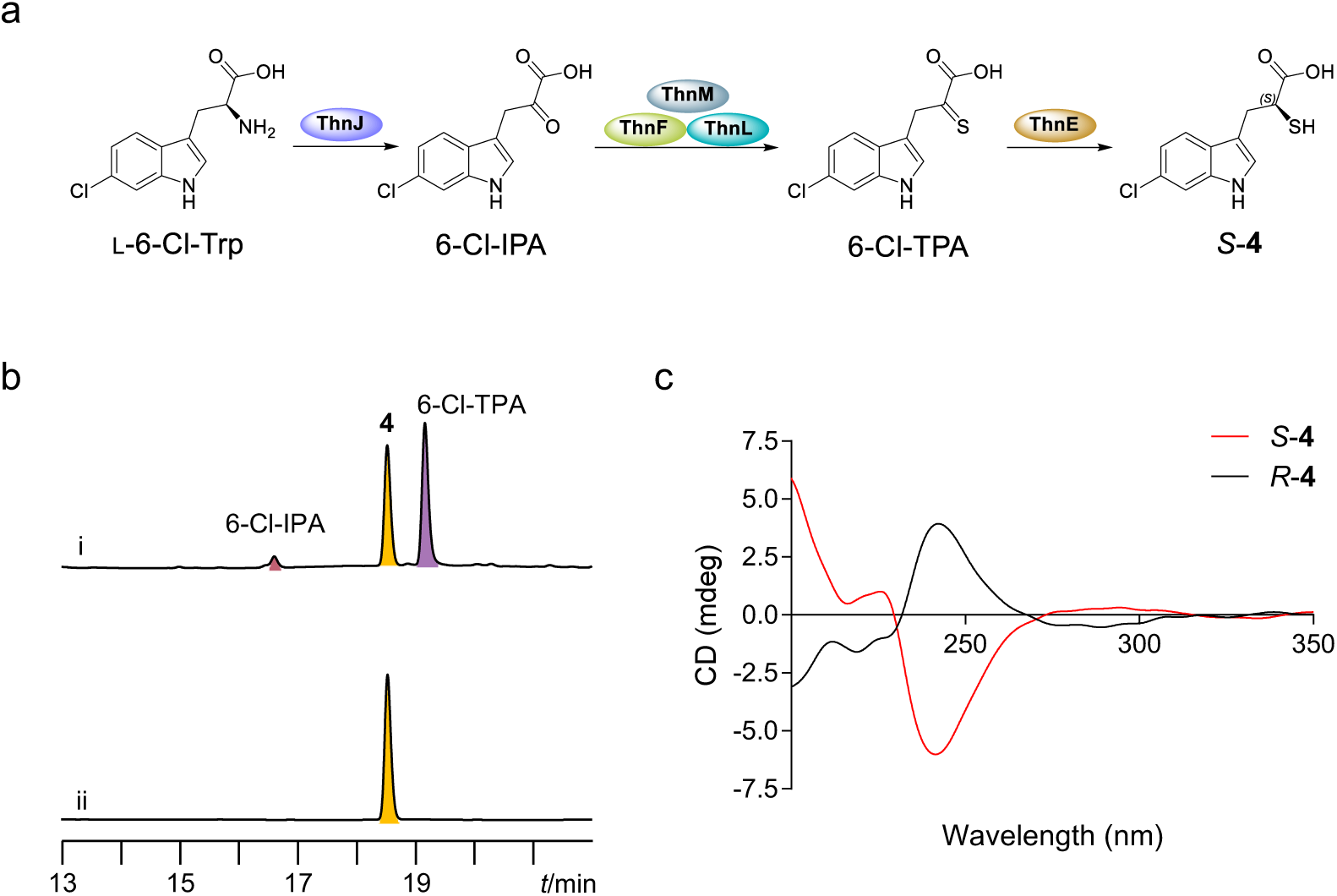
Construction of the sulfur incorporation pathway in the biosynthesis of Thienodolin. **a**, Biosynthetic pathway to compound *S*-**4**. **b**, HPLC analysis of the one-pot enzymatic reaction. (i) Incubation of _L_-6-Cl-Trp with ThnJ, ThnF, ThnL, and ThnM in the presence of ATP, MgCl_2_, Na_2_S_2_O_3_, and dithiothreitol. (ii) Subsequent addition of ThnE and NADPH to the reaction mixture of (i). **c**, Configuration analysis by circular dichroism (CD) spectroscopy. The *S*-**4** diastereomer was obtained via ThnE-mediated reduction, while *R*-**4** was produced by known 2-dehydropantoate reductase Cxm6 from chuangxinmycin biosynthesis.

**Extended Data Fig. 7.**
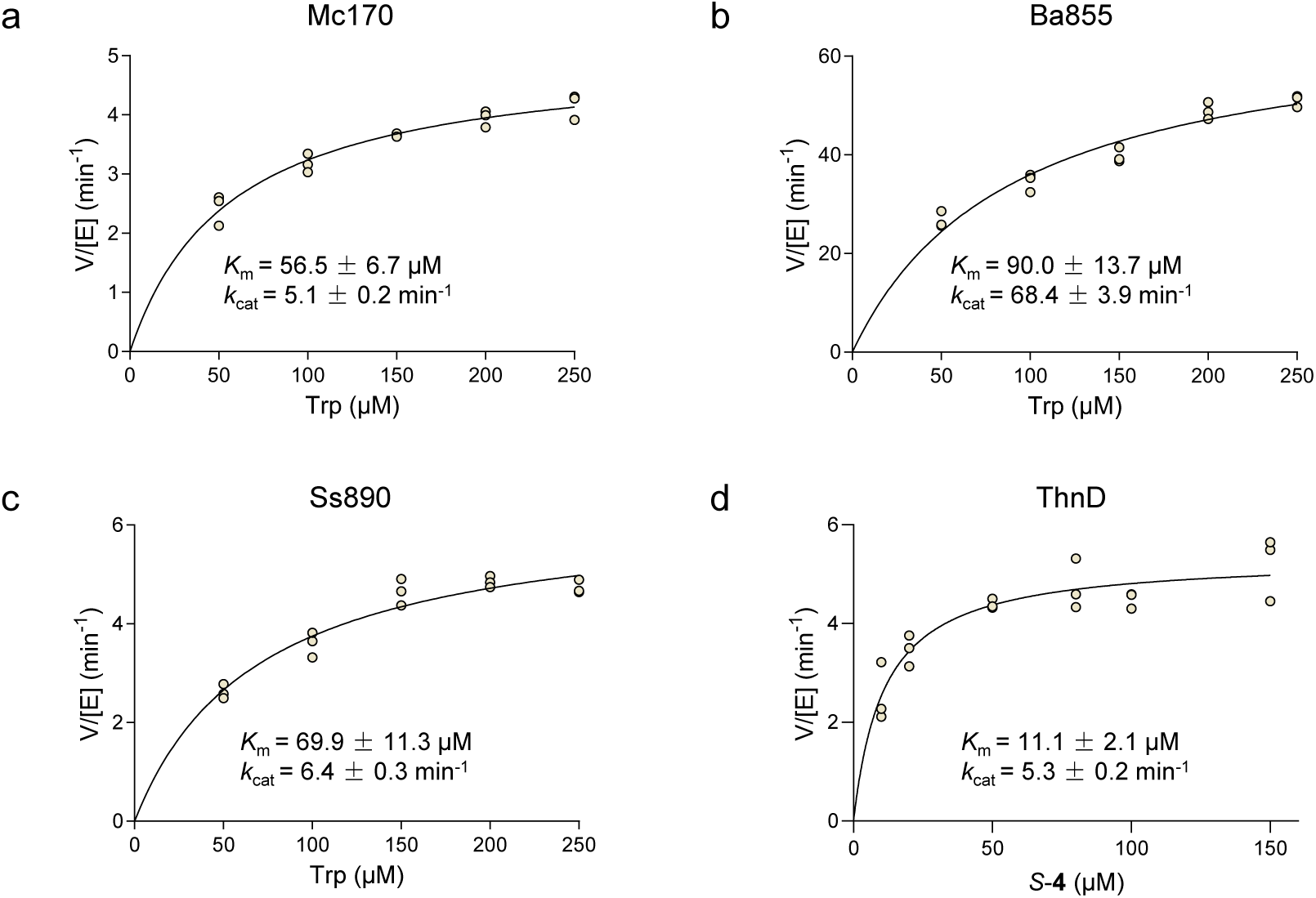
Kinetic analyses of selected cytochrome P422 enzymes. Initial rate kinetics were measured in the presence of *Sel*Fdx1499 (100 μM), *Sel*FdR0978 (50 μM), NADPH (10 mM), and sodium ascorbate (10 mM) for: **a**, Mc170 with L-Trp; **b**, Ba855 with L-Trp; **c**, Ss890 with L-Trp; **d**, ThnD with *S*-**4**.

**Extended Data Fig. 8.**
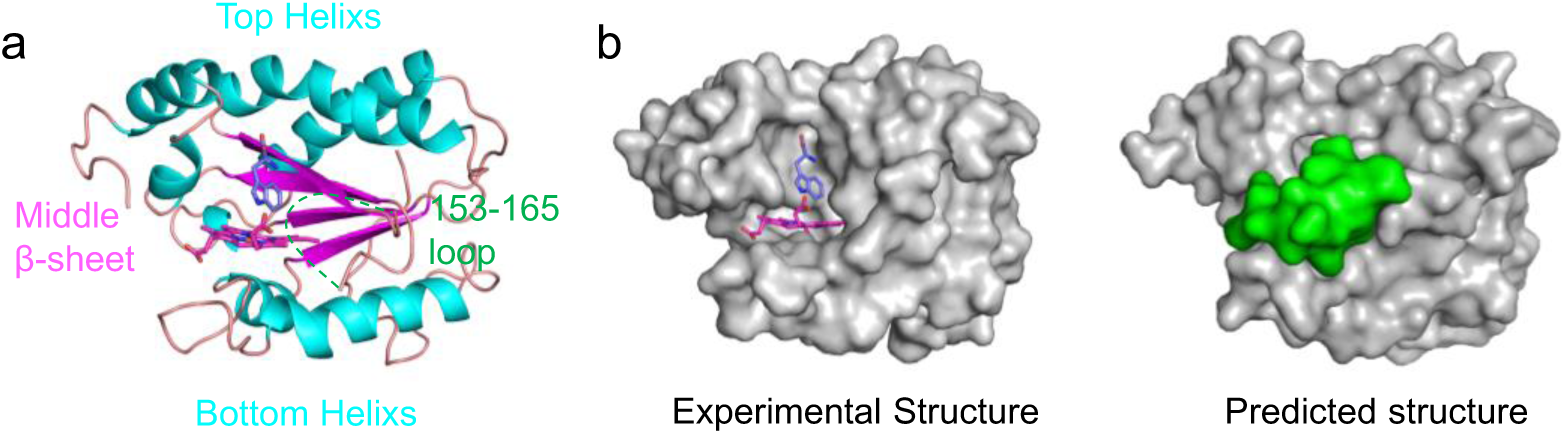
Structural architecture of Mc170. **a**, Overall sandwich-like fold of Mc170. **b**, Hydrophobic surface representation of the experimental crystal structure (left) and the AlphaFold3-predicted model (right). The loop comprising residues 153–165 is colored green to highlight its hydrophobic character.

**Extended Data Fig. 9.**
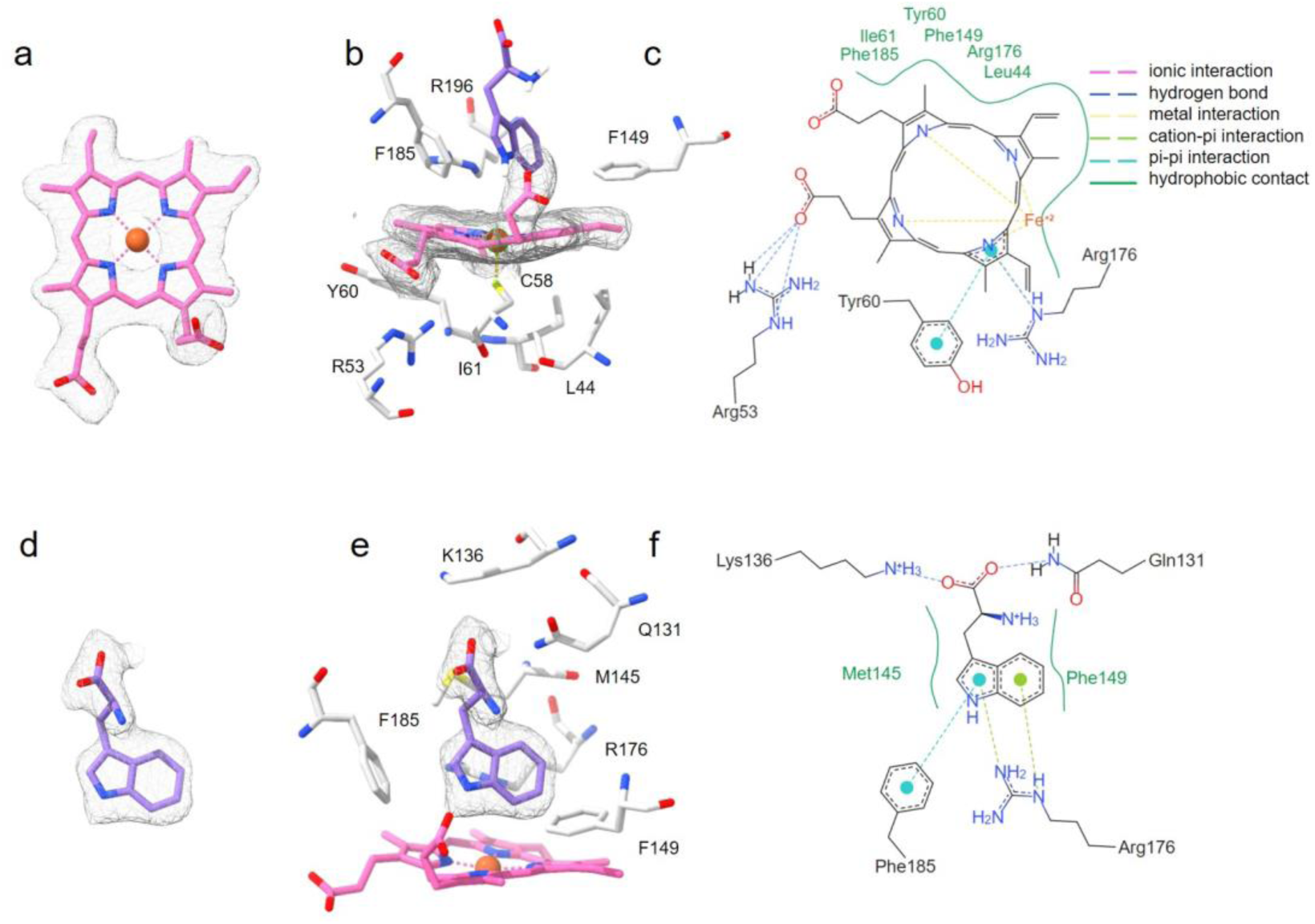
Electron density and binding interactions in the Mc170 active site. **a**, Omit 2*F*₀–*F*₀ electron density map (contoured at 1.0σ, gray) for the heme group (pink carbons). **b**, Detailed view of the heme-binding site in the Mc170 crystal structure. **c**, Two-dimensional interaction diagram of residues coordinating the heme cofactor. **d**, Omit 2*F*₀–*F*₀ electron density map (contoured at 1.0σ, gray) for the tryptophan substrate (purple carbons). **e**, Detailed view of the tryptophan-binding site. **f**, Two-dimensional interaction diagram of residues involved in tryptophan recognition.

**Extended Data Fig. 10.**
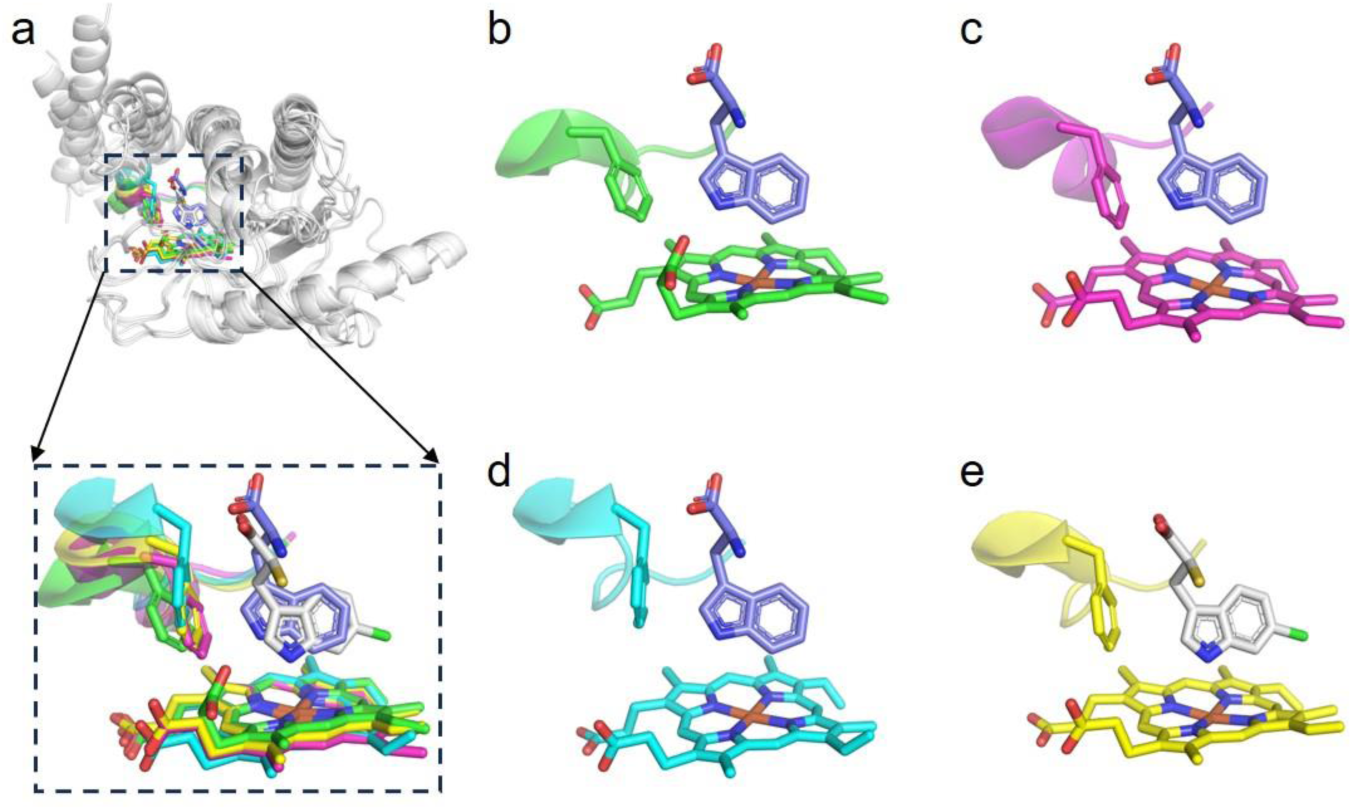
Structural comparison and ligand docking across cytochrome P422 enzymes. **a**, Structure-based alignment of Mc170 (green), Ss890 (cyan), Ba855 (red), and ThnD (yellow). **b**–**d**, Predicted binding pose of tryptophan (purple) in the active sites of Mc170 (**b**), Ss890 (**c**), and Ba855 (**d**), based on alignment with the Mc170–Trp crystal structure. **e**, Predicted conformation of substrate *S*-**4** (gray) in ThnD, as modeled by AlphaFold3.

## References

1 Barik, S. The uniqueness of tryptophan in biology: properties, metabolism, interactions and localization in proteins. Int. J. Mol. Sci. 21, 22 8776 (2020).

2 Xue, C. et al. Tryptophan metabolism in health and disease. Cell Metab. 35, 1304–1326 (2023).

3 Alkhalaf, L. M. & Ryan, K. S. Biosynthetic manipulation of tryptophan in bacteria: pathways and mechanisms. Chem. Biol. 22, 317–328 (2015).

4 Meng, B. et al. Structural and functional analyses of human tryptophan 2,3-dioxygenase. Proteins: Struct., Funct., Bioinf. 82, 3210–3216 (2014).

5 Li, Z. et al. Engineering cytochrome P450 enzyme systems for biomedical and biotechnological applications. J. Biol. Chem. 295, 833–849 (2020).

6 Shin, I., Wang, Y. & Liu, A. A new regime of heme-dependent aromatic oxygenase superfamily. Proc. Natl. Acad. Sci. U. S. A. 118, e2106561118 (2021).

7 Yu, T. et al. Enzyme function prediction using contrastive learning. Science 379, 1358–1363 (2023).

8 Kim, G. B. et al. Functional annotation of enzyme-encoding genes using deep learning with transformer layers. Nat. Commun. 14, 7370 (2023).

9 Coelho, P. S. et al. A serine-substituted P450 catalyzes highly efficient carbene transfer to olefins in vivo. Nat. Chem. Biol. 9, 485–487 (2013).

10 Wen, J. & Shi, Z. From C4 to C7: innovative strategies for site-selective functionalization of indole C–H bonds. Acc. Chem. Res. 54, 1723–1736 (2021).

11 Wohlwend, J., et al. Boltz-1 democratizing biomolecular interaction modeling. bioRxiv, 2024.11.19.624167 (2025).

12 Boutet, E., Lieberherr, D., Tognolli, M., Schneider, M. & Bairoch, A. UniProtKB/Swiss-Prot. Methods Mol Biol. 406, 89–112 (2007).

13 Nelder, J. A. & Wedderburn, R. W. M. Generalized linear models. J. R. Stat. Soc. Ser. A Stat. Soc. 135, 370–384 (2018).

14 Peretz, O., Koren, M. & Koren, O. Naive Bayes classifier – An ensemble procedure for recall and precision enrichment. Eng. Appl. Artif. Intell. 136, 108972 (2024).

15 Breiman, L. Random Forests. Mach. Learn 45, 5–32 (2001).

16 Bergstra, J. & Bengio, Y. Random search for hyper-parameter optimization. J. Mach. Learn. Res. 13, 281–305 (2012).

17 Hiessl, S. et al. Latex clearing protein—an oxygenase cleaving poly(*cis*-1,4-isoprene) rubber at the cis double bonds. Appl. Environ. Microbiol. 80, 5231–5240 (2014).

18 Zhang, W. et al. Compartmentalized biosynthesis of mycophenolic acid. Proc. Natl. Acad. Sci. U. S. A. 116, 13305–13310 (2019).

19 Zallot, R., Oberg, N. & Gerlt, J. A. The EFI web resource for genomic enzymology tools: leveraging protein, genome, and metagenome databases to discover novel enzymes and metabolic pathways. Biochemistry 58, 4169–4182 (2019).

20 Oberg, N., Zallot, R. & Gerlt, J. A. EFI-EST, EFI-GNT, and EFI-CGFP: enzyme function initiative (EFI) web resource for genomic enzymology tools. J. Mol. Biol. 435, 168018 (2023).

21 Keller, U., Lang, M., Crnovcic, I., Pfennig, F. & Schauwecker, F. The actinomycin biosynthetic gene cluster of *Streptomyces chrysomallus*: a genetic hall of mirrors for synthesis of a molecule with mirror symmetry. J. Bacteriol. 192, 2583–2595 (2010).

22 Liu, M. et al. Identification of the actinomycin D biosynthetic pathway from marine-derived *Streptomyces costaricanus* SCSIO ZS0073. Mar. Drugs 17, 240 (2019).

23 Zhang, C. et al. Genome mining for mycemycin: discovery and elucidation of related methylation and chlorination biosynthetic chemistries. Org. Lett. 20, 7633–7636 (2018).

24 Song, F., Liu, N., Liu, M., Chen, Y. & Huang, Y. Identification and characterization of mycemycin biosynthetic gene clusters in *Streptomyces olivaceus* FXJ8.012 and *Streptomyces* sp. FXJ1.235. Mar. Drugs 16, 3 98 (2018).

25 Wang, Y. et al. Identifying the minimal enzymes for unusual carbon-sulfur bond formation in thienodolin biosynthesis. ChemBioChem 17, 799–803 (2016).

26 Milbredt, D., Patallo, E. P. & van Pee, K. H. A tryptophan 6-halogenase and an amidotransferase are involved in thienodolin biosynthesis. ChemBioChem 15, 1011–1020 (2014).

27 Poulos, T. L. Heme enzyme structure and function. Chem. Rev. 114, 3919–3962 (2014).

28 Guengerich, F. P., Martin, M. V., Sohl, C. D. & Cheng, Q. Measurement of cytochrome P450 and NADPH–cytochrome P450 reductase. Nat. Protoc. 4, 1245–1251 (2009).

29 Zhang, W. et al. Mechanistic insights into interactions between bacterial class I P450 enzymes and redox partners. ACS Catal. 8, 9992–10003 (2018).

30 Shi, X. et al. Hydroxytryptophan biosynthesis by a family of heme-dependent enzymes in bacteria. Nat. Chem. Biol. 19, 1415–1422 (2023).

31 Ma, G. L. et al. Biosynthesis of tasikamides via pathway coupling and diazonium-mediated hydrazone formation. J. Am. Chem. Soc. 144, 1622–1633 (2022).

32 Milbredt, D., Patallo, E. P. & van Pee, K. H. Characterization of the aminotransferase ThdN from thienodolin biosynthesis in *Streptomyces albogriseolus*. ChemBioChem 17, 1859–1864 (2016).

33 Zhang, X., et al. Biosynthesis of chuangxinmycin featuring a deubiquitinase-like sulfurtransferase. Angew. Chem., Int. Ed. Engl. 60, 24418–24423 (2021).

34 Dunbar, K. L., Scharf, D. H., Litomska, A. & Hertweck, C. Enzymatic carbon–sulfur bond formation in natural product biosynthesis. Chem. Rev. 117, 5521–5577 (2017).

35 Cheng, R. et al. Single-step replacement of an unreactive C–H bond by a C–S bond using polysulfide as the direct sulfur source in the anaerobic ergothioneine biosynthesis. ACS Catal. 10, 8981–8994 (2020).

36 Ushimaru, R. & Abe, I. C–N and C–S bond formation by cytochrome P450 enzymes. Trends Chem. 5, 526–536 (2023).

37 Abramson, J. et al. Accurate structure prediction of biomolecular interactions with AlphaFold 3. Nature 630, 493–500 (2024).

38 Guengerich, F. P. & Yoshimoto, F. K. Formation and cleavage of C–C bonds by enzymatic oxidation–reduction reactions. Chem. Rev. 118, 6573–6655 (2018).

39 Ma, N., He, T., Johnston, L. J. & Ma, X. Host–microbiome interactions: the aryl hydrocarbon receptor as a critical node in tryptophan metabolites to brain signaling. Gut Microbes 11, 1203–1219 (2020).

40 Lv, J. et al. Metal-free directed *sp*^2^-C–H borylation. Nature 575, 336–340 (2019).

## References

41 Steinegger, M. & Söding, J. MMseqs2 enables sensitive protein sequence searching for the analysis of massive data sets. Nat. Biotechnol. 35, 1026–1028 (2017).

42 Mirdita, M. et al. ColabFold: making protein folding accessible to all. Nat. Methods 19, 679–682 (2022).

43 Fufezan, C., Zhang, J. & Gunner, M. R. Ligand preference and orientation in *b*- and *c*-type heme-binding proteins. Proteins 73, 690–704 (2008).

44 Fassio, A. V. et al. Prioritizing virtual screening with interpretable interaction fingerprints. J. Chem. Inf. Model. 62, 4300–4318 (2022).

45 Li, T., Bonkovsky, H. L. & Guo, J. T. Structural analysis of heme proteins: implications for design and prediction. BMC Struct. Biol. 11, 13 (2011).

46 Jumper, J. et al. Highly accurate protein structure prediction with AlphaFold. Nature 596, 583–589 (2021).

47 Lomsadze, A., Gemayel, K., Tang, S. & Borodovsky, M. Modeling leaderless transcription and atypical genes results in more accurate gene prediction in prokaryotes. Genome Res. 28, 1079–1089 (2018).

48 Cantalapiedra, C. P., Hernández-Plaza, A., Letunic, I., Bork, P. & Huerta-Cepas, J. eggNOG-mapper v2: functional annotation, orthology assignments, and domain prediction at the metagenomic scale. Mol. Biol. Evol. 38, 5825–5829 (2021).

49 Minor, W., Cymborowski, M., Otwinowski, Z. & Chruszcz, M. HKL-3000: the integration of data reduction and structure solution--from diffraction images to an initial model in minutes. *Acta Crystallogr.*, Sect. D:Biol. Crystallogr. 62, 859–866 (2006).

50 McCoy, A. J. et al. Phaser crystallographic software. J. Appl. Crystallogr. 40, 658–674 (2007).

51 Emsley, P. & Cowtan, K. Coot: model-building tools for molecular graphics. Acta Crystallogr., Sect. D:Biol. Crystallogr. 60, 2126–2132 (2004).

52 Adams, P. D. et al. PHENIX: a comprehensive Python-based system for macromolecular structure solution. Acta Crystallogr., Sect. D:Biol. Crystallogr. 66, 213–221 (2010).

53 Davis, I. W. et al. MolProbity: all-atom contacts and structure validation for proteins and nucleic acids. Nucleic Acids Res. 35, W375–383 (2007).

54 Pettersen, E. F. et al. UCSF ChimeraX: Structure visualization for researchers, educators, and developers. Protein Sci. 30, 70–82 (2021).

55 Xu, H. et al. In vitro oxidative decarboxylation of free fatty acids to terminal alkenes by two new P450 peroxygenases. Biotechnol. Biofuels 10, 208 (2017).

56 Larkin, M. A. et al. Clustal W and Clustal X version 2.0. Bioinformatics 23, 2947–2948 (2007).

57 Kumar, S., Nei, M., Dudley, J. & Tamura, K. MEGA: a biologist-centric software for evolutionary analysis of DNA and protein sequences. Briefings Bioinf. 9, 299–306 (2008).

58 Xie, J. et al. Tree visualization by one table (tvBOT): a web application for visualizing, modifying and annotating phylogenetic trees. Nucleic Acids Res. 51, W587–W592 (2023).

