## Supplementary materials for "Tryptophan Chemistry Driven by a Widespread Cytochrome P422 Enzyme Family"

Growth medium components were obtained from MDBio Biotech Co., Ltd. (Qingdao, China). Antibiotics and sodium ascorbate were sourced from Solarbio (Beijing, China). L-6-chlorotryptophan (L-6-Cl-Trp) and 6-hydroxytryptophan (6-OH-Trp) were procured from Shanghai Haohong Scientific Co., Ltd. (Shanghai, China). L-Tryptophan (L-Trp), ATP, DTT, and sodium thiosulfate ($Na_2S_2O_3$) were supplied by Shanghai Macklin Biochemical Technology Co., Ltd. (Shanghai, China). NADPH was purchased from BONTAC (Shenzhen, China). Enzymes and molecular weight markers for cloning were obtained from Fisher Scientific (PA, USA). Plasmid mini-prep and DNA gel extraction kits were acquired from Omega Bio-tek, Inc. (GA, USA). The protein expression vectors pET28a and pET28a-SUMO were procured from TsingKe Bio Technology (Beijing, China). Oligonucleotide primers and DNA sequencing services were provided by Sangon Biotech (Shanghai, China). Ni-NTA Sefinose Resin (Settled Resin) for protein purification was obtained from Sangon Biotech (Shanghai, China). All synthetic genes were ordered from BGI Genomics Co., Ltd. (Shenzhen, China). Additional chemicals and reagents were purchased from Sinopharm Chemical Reagent Co., Ltd. (Shanghai, China).

**Data availability statement**

The atomic coordinates and structure factors have been deposited in the Protein Data Bank with accession codes 9X39.

**Supplementary Table 1:** Settings of key parameters of all models.

| Models | Key Parameters |
|---|---|
| Logistic Regression | penalty: L2, solver: lbfgs |
| Naive Bayes | priors: none |
| Random Forest | max_depth: 5, n_estimators: 100<br>min_samples_leaf: 1<br>min_samples_split: 2 |
| Threshold Screening | rang of complex_iplddt: 0.1-0.8<br>rang of iPTM: 0.1-0.8<br>rang of surrounding residue number: 8-30 |

**Supplementary Table 2:** The screening result of from *Streptomyces coelicolor* A3(2) by CISSspector.

| HBPs | | |
|---|---|---|
| ID | Ligand | Annotation |
| CAB36612.1 | TYR337 | catalase |
| CAC17503.1 | CYS360 | cytochrome P450 |
| CAB88824.1 | CYS351 | cytochrome P450 |
| CAB61278.1 | CYS362 | cytochrome P450 |
| CAB61183.1 | TYR418 | catalase |
| CAB53427.1 | HIS142 | putative two-component sensor |
| CAA22217.1 | HIS189, MET339 | cytochrome oxidase subunit I |

| | | |
|---|---|---|
| CAB88975.1 | CYS356 | cytochrome P450 |
| CAB42033.1 | HIS229 | oxidoreductase |
| CAB76337.1 | CYS349 | cytochrome P450 |
| CAD55402.1 | CYS368 | cytochrome P450 |
| CAC09540.1 | CYS366 | cytochrome P450 |
| CAD55330.1 | HIS194 | regulatory protein |
| CAB56738.1 | MET111, MET343 | putative membrane protein |
| CAB94657.1 | HIS100, HIS413 | cytochrome C oxidase polypeptide I |
| CAC10308.1 | CYS342 | cytochrome P450 |
| CAB66201.1 | CYS440 | cytochrome P450 |
| CAB39882.1 | HIS403, HIS90 | cytochrome c oxidase subunit I |
| CAB88947.1 | HIS102 | MPAB_Lcp_cat |
| CAB89766.1 | HIS88 | flavohemoprotein |
| CAB61277.1 | CYS354 | cytochrome P450 |
| CAB39713.1 | HIS192 | MPAB_Lcp_cat |
| CAC16511.1 | TYR338 | catalase |
| CAA22220.1 | HIS127 | putative two component sensor |
| CAB61722.1 | HIS326 | putative membrane protein |
| CAC01593.1 | HIS225 | conserved hypothetical protein |
| CAB76336.1 | CYS355 | cytochrome P450 |
| CAC01489.1 | CYS353 | cytochrome P450 |
| CAD30936.1 | HIS188, HIS66 | nitrate reductase gamma chain NarI3 |
| CAB59479.1 | CYS368 | cytochrome P450 |
| CAB94608.1 | CYS410 | cytochrome P450 |
| CAB58320.1 | TYR340 | catalase |
| CAC08382.1 | HIS218, HIS365 | putative cytochrome biogenesis related protein |
| CAB39875.1 | HIS118, HIS219 | cytochrome B subunit |
| CAC14339.1 | CYS361 | cytochrome P450 |
| CAA20629.1 | HIS184, HIS63 | nitrate reductase gamma chain NarI |
| CAB39877.1 | HIS159, MET196 | cytochrome C heme-binding subunit |
| CAB42023.1 | CYS363 | cytochrome P450 |
| CAB88825.1 | CYS354 | cytochrome P450 |
| CAB76347.1 | HIS88 | flavohemoprotein |
| CAB57412.1 | HIS272 | catalase |
| CAC37531.1 | CYS351 | cytochrome P450 |
| CAC04226.1 | HIS116, HIS217 | ubiquinol-cytochrome C reductase cytochrome B subunit |
| CAB52917.1 | HIS125 | flavohemoprotein |
| Non-HBPs | | |
| ID | Ligand | Annotation |
| CAB50755.1 | HIS250, HIS311 | COX15/CtaA Heme A synthase, prokaryotes |
| CAD55179.1 | HIS110 | Oxygen sensor histidine kinase NreB |
| CAB59442.1 | HIS200, LYS278 | CopD |
| CAC32324.1 | HIS292, LYS363 | CopD |
| CAC01309.1 | HIS268 | peptidase |

| | | |
|---|---|---|
| CAD55467.1 | HIS310, LYS386 | CopD |
| CAB76993.1 | HIS105, HIS177 | Oxidored_molyb |
| CAB76065.1 | LYS141 | aminodeoxychorismate lyase |
| CAB92889.1 | CYS246 | porphobilinogen deaminase |

**Supplementary Table 3:** Statistical analysis of heme binding protein types screened from 100 strains of *Actinomycetes*.

| HBPs | | |
|---|---|---|
| **Pfam/InterPro ID** | **Pfam/InterPro number** | **Quantity** |
| MPAB_Lcp_cat (DUF2236) | PF09995 | 48 |
| Cyt_P450_B | IPR002397 | 44 |
| DUF2231 | PF09990 | 7 |
| DUF6875 | PF21780 | 6 |
| Catalase_sf | IPR020835 | 7 |
| DUF3291 | PF11695 | 6 |
| DUF1365 | PF07103 | 4 |
| HODM_asu-like | PF11927 | 3 |
| Tll0287-like | PF11845 | 3 |
| DUF3483 | PF11982 | 2 |
| RoxA-like_Cyt-c | PF21419 | 2 |
| Globin-like_sf | IPR009050 | 2 |
| Multihaem_cyt_sf | IPR036280 | 2 |
| DUF1516 | PF07457 | 2 |
| DUF2306 | PF10067 | 3 |
| Cyt_c-like_dom_sf | IPR036909 | 3 |
| Cytochrom_C | PF00034 | 2 |
| Trp/Indoleamine_2_3_dOase-like | IPR037217 | 2 |
| SO_2930-like_C | IPR022269 | 1 |
| CCP_MauG | PF03150 | 1 |
| PrnB | PF08933 | 1 |
| DUF4405 | PF14358 | 1 |
| Dyp_peroxidase | IPR006314 | 2 |
| NarG-like_sf | IPR036197 | 1 |
| DUF1415 | PF07209 | 1 |
| DUF998 | PF06197 | 1 |
| ActD | PF11821 | 1 |
| SQR/QFR_C/D | IPR034804 | 1 |
| Monooxy_af470-like | PF13826 | 1 |
| Ni_hydr_CYTB | PF01292 | 1 |
| DUF3492 | PF11997 | 1 |
| DUF4079 | PF13301 | 1 |
| DUF1440 | PF07274 | 1 |
| Non-HBPs | | |

| Pfam/InterPro ID | Pfam/InterPro ID | Pfam/InterPro ID |
|---|---|---|
| TPR-like_helical_dom_sf | IPR011990 | 1 |
| Pkinase | PF00069 | 1 |
| AB_hydrolase_fold | IPR029058 | 1 |
| HAMP | PF00672 | 1 |
| DUF2269 | PF10027 | 4 |
| DUF6879 | PF21806 | 1 |
| Por_Secre_tail | PF18962 | 2 |
| PCuAC | PF04314 | 1 |
| Alkaline_phosphatase_core_sf | IPR017850 | 1 |
| No known domain ("-") | - | 14 |

**Supplementary Table 4:** The model CLEAN prediction results.

| Enzymes | Predicted EC Number | Corresponding Types | Depends on Heme | Actual natural substrates and reactions |
|---|---|---|---|---|
| Ss890 | 1.3.7.3 | ferredoxin oxidoreductase | No | Trp, 6-hydroxylation |
| Ba855 | 1.3.7.5 | ferredoxin oxidoreductase | No | Trp, 7-hydroxylation |
| Mc170 | 1.3.7.5 | ferredoxin oxidoreductase | No | Trp, 1-hydroxylation |
| ThnD | 2.1.1.148 | thymidylate synthase | No | Compound **4**, C-S bond formation |

**Supplementary Table 5:** Features of Solubilized Proteins in this study.

| Protein name | Uniprot ID | Phylum | Sequence length | Reaction type | Organism |
|---|---|---|---|---|---|
| Sa575 | A0AAP6BMA6 | Actinomycetota | 225 | unknown | *Streptomyces acidiscabies.* |
| Ss537 | A0A286I1H9 | Actinomycetota | 217 | unknown | *Streptomyces* sp. 2323.1. |
| Ss860 | A0A7X1IS07 | Actinomycetota | 232 | 6-OH | *Streptomyces* sp. TYQ1024. |
| Am023 | C6WFN5 | Actinomycetota | 215 | 7-OH | *Actinosynnema mirum* |
| Ss420 | A0A6C0Q126 | Actinomycetota | 220 | 7-OH | *Streptomyces* sp. S4.7. |
| Sn658 | A0A853BV61 | Actinomycetota | 203 | 7-OH | *Streptomonospora nanhaiensis.* |
| AcmT | D6R234 | Actinomycetota | 211 | 7-OH | *Streptomyces anulatus* |
| Sg280 | A0A4U5W5J8 | Actinomycetota | 211 | 7-OH | *Streptomyces galbus* |
| ActG7' | K8FE45 | Actinomycetota | 219 | 7-OH | *Streptomyces iakyrus.* |
| Ss850 | A0A3N6HL00 | Actinomycetota | 210 | 7-OH | *Streptomyces* sp. ADI98-10 |
| AcmE | D6R241 | Actinomycetota | 235 | 7-OH | *Streptomyces anulatus* |
| MymH | A0A2R3ZQ27 | Actinomycetota | 220 | 7-OH | *Streptomyces olivaceus.* |

| | | | | | |
|---|---|---|---|---|---|
| Sd870 | K4RBM6 | Actinomycetota | 230 | 7-OH | *Streptomyces davaonensis* |
| Ar890 | A0A154MRM9 | Actinomycetota | 216 | 7-OH | *Amycolatopsis regifaucium.* |
| Sp162 | A0A840Q4G5 | Actinomycetota | 209 | 7-OH | *Saccharopolyspora phatthalungensis.* |
| Si625 | A0A0G3APM5 | Actinomycetota | 237 | 7-OH | *Streptomyces incarnatus.* |
| Ts115 | A0A2N7WBB0 | Pseudomonadota | 247 | 7-OH | *Trinickia soli.* |
| Ps640 | A0A4R5M467 | Pseudomonadota | 245 | 7-OH | *Paraburkholderia silviterrae* |
| Ps666 | A0A4R6F500 | Pseudomonadota | 245 | 7-OH | *Paraburkholderia* sp. BL10I2N1 |
| Ne015 | A0A1I2HNG7 | Myxococcota | 237 | 6-OH | *Nannocystis exedens.* |
| Cm015 | A0A1Z1FA26 | Pseudomonadota | 223 | 1-OH | *Croceicoccus marinus.* |
| Rs365 | A0A3S2X2H1 | Pseudomonadota | 231 | 1-OH | *Rhizobium* sp. RMa-01 |
| Cs407 | A0A158HAL1 | Pseudomonadota | 201 | 1-OH | *Caballeronia sordidicola.* |
| Pm135 | A0A149PGL8 | Pseudomonadota | 205 | 1-OH | *Paraburkholderia monticola.* |
| Sh310 | A0A7W2D8B4 | Actinomycetota | 225 | unknown | *Streptomyces himalayensis* subsp. *aureolus* |
| Bi410 | A0A508T3H2 | Pseudomonadota | 254 | unknown | *Bradyrhizobium ivorense.* |
| Ba855 | A0A837XV78 | Bacillota | 217 | 7-OH | *Bacillus atrophaeus.* |
| Pp145 | A0A7D5ZYA0 | Pseudomonadota | 238 | unknown | *Pseudomonas putida* |
| Sa413 | A0A1Z2KYE7 | Actinomycetota | 237 | 6-OH | *Streptomyces albireticuli.* |
| Rr391 | A0A1I4PGA9 | Pseudomonadota | 256 | 1-OH | *Rugamonas rubra.* |
| Ls950 | A0A7Y4Y5E2 | Pseudomonadota | 254 | 1-OH | *Lysobacter* sp |
| Ls910 | A0A0Q8D691 | Pseudomonadota | 285 | 1-OH | *Lysobacter* sp. Root604. |
| Lc584 | A0A108UBG6 | Pseudomonadota | 271 | 1-OH | *Lysobacter capsici* AZ78. |
| ThdM | A0A142I752 | Actinomycetota | 223 | C-S | *Streptomyces albogriseolus* |
| ThnD | A0A2B8AL05 | Actinomycetota | 202 | C-S | *Streptomyces* sp. FXJ1.172. |
| Ns565 | A0A8J6WP09 | Cyanobacteriota | 252 | 1-OH | *Nodosilinea* sp. FACHB-141. |
| Os274 | A0A978SBE7 | Cyanobacteriota | 242 | 1-OH | *Leptolyngbyaceae cyanobacterium* M33_DOE_097. |
| Cf970 | A0A3S0ZQG6 | Cyanobacteriota | 242 | 1-OH | *Chlorogloeopsis fritschii* PCC 6912. |
| Ns075 | A0A6P1KUG2 | Cyanobacteriota | 225 | 1-OH | *Nostoc* sp. ATCC 53789 |
| Ss890 | A0AA91H005 | Cyanobacteriota | 220 | 6-OH | *Scytonema* sp. HK-05. |
| Ss555 | A0A856MJE7 | Cyanobacteriota | 236 | 7-OH | *Brasilonema sennae* CENA114. |
| Sh520 | A0A139WVV4 | Cyanobacteriota | 229 | 1-OH | *Scytonema hofmannii* PCC 7110. |
| Ah290 | A0AAP5M4T2 | Cyanobacteriota | 256 | 1-OH | *Aetokthonos hydrillicola* Thurmond2011. |

| | | | | | |
|---|---|---|---|---|---|
| Ts635 | A0A6G9SJS3 | Cyanobacteriota | 224 | 1-OH | *Tolypothrix* sp. PCC 7910. |
| Nc649 | A0A2H6LNR8 | Cyanobacteriota | 230 | 1-OH | *Nostoc cycadae* WK-1. |
| Xn269 | D3VI28 | Pseudomonadota | 234 | 6-OH | *Xenorhabdus nematophila* |
| Xk970 | A0A1I7GRM0 | Pseudomonadota | 234 | 6-OH | *Xenorhabdus koppenhoeferi.* |
| Pc535 | A0A089WT68 | Pseudomonadota | 230 | 6-OH | *Pseudomonas cremoricolorata* |
| Ps625 | A0A0Q8BRF7 | Pseudomonadota | 244 | 6-OH | *Pseudomonas* sp. Root569. |
| Es115 | A0AAP8VLS8 | Pseudomonadota | 219 | unknown | *Ensifer* sp. NM-2. |
| Ms995 | A0A6I3ZX66 | Actinomycetota | 212 | 1-OH | *Mycolicibacterium* sp. CBMA 226. |
| As285 | A0A848KAF2 | Actinomycetota | 220 | 1-OH | *Antrihabitans stalactiti.* |
| Hs201 | F6EF82 | Actinomycetota | 218 | 1-OH | *Hoyosella subflava* |
| Rs537 | A0A0D8HYK3 | Actinomycetota | 217 | 1-OH | *Rhodococcus* sp. (strain AD45) |
| Nj260 | A0A917VT16 | Actinomycetota | 220 | 1-OH | *Nocardia jinanensis.* |
| Nb099 | A0A543F3X1 | Actinomycetota | 251 | 6-OH | *Nocardia bhagyanarayanae* |
| Nc521 | A0A231H7J4 | Actinomycetota | 231 | 6-OH | *Nocardia cerradoensis* |
| Rr100 | A0A562E549 | Actinomycetota | 234 | 6-OH | *Rhodococcus rhodochrous* J45. |
| Nm520 | A0A7K0D322 | Actinomycetota | 254 | unknown | *Nocardia macrotermitis.* |
| Mb565 | A0A850DMP1 | Actinomycetota | 224 | unknown | *Mycobacteriaceae bacterium.* |
| Nt485 | A0A7K1UWC0 | Actinomycetota | 232 | 6-OH | *Nocardia terrae* |
| Na660 | A0A6G9YPW7 | Actinomycetota | 232 | 6-OH | *Nocardia arthritidis.* |
| Ns410 | A0A0Q7NTW8 | Actinomycetota | 225 | 6-OH | *Nocardia* sp. Root136. |
| Nb805 | A0A6G9XXX9 | Actinomycetota | 235 | 6-OH | *Nocardia brasiliensis* |
| Ms021 | A0A562WKB5 | Actinomycetota | 210 | 1-OH | *Micromonospora sagamiensis.* |
| Aw136 | A0A4R2J5W9 | Actinomycetota | 223 | 1-OH | *Actinocrispum wychmicini* |
| At245 | A0A229RLY4 | Actinomycetota | 220 | 1-OH | *Amycolatopsis thailandensis* |
| Aa431 | A0A1H0M606 | Actinomycetota | 219 | 1-OH | *Actinokineospora alba* |
| Ss540 | A0A9K3T6T0 | Actinomycetota | 211 | 1-OH | *Streptomyces* sp. SID4917. |
| Ss575 | A0A1Q5MKF1 | Actinomycetota | 211 | 1-OH | *Streptomyces* sp. CB00455 |
| Ss515 | A0A384HJB5 | Actinomycetota | 211 | 1-OH | *Streptomyces* sp. AC1-42W |
| Ss795 | A0A2S9PXX9 | Actinomycetota | 230 | 1-OH | *Streptomyces solincola.* |
| Ss435 | A0A6G9FAW0 | Actinomycetota | 212 | 1-OH | *Streptomyces* sp. Tu 2975. |
| Mc170 | A0A556MU32 | Bacteroidota | 213 | 1-OH | *Mucilaginibacter corticis.* |
| Cc488 | A0A410RXA3 | Myxococcota | 224 | 1-OH | *Corallococcus coralloides* |
| Si800 | A0A918UUE0 | Actinomycetota | 190 | 1-OH | *Streptomyces inusitatus* |
| Ac095 | A0A542DBL9 | Actinomycetota | 210 | 1-OH | *Amycolatopsis cihanbeyliensis* |
| Sv279 | A0A495X981 | Actinomycetota | 224 | 1-OH | *Saccharothrix variisporea* |
| Me959 | A0A1C4Z9T6 | Actinomycetota | 244 | 1-OH | *Micromonospora echinospora* |
| Mp190 | A0A7W7WSA9 | Actinomycetota | 240 | 1-OH | *Micromonospora polyrhachis* |

| | | | | | |
|---|---|---|---|---|---|
| Sr190 | D2AY02 | Actinomycetota | 226 | 1-OH | *Streptosporangium roseum* |
| Ne852 | A0A7W8EGY6 | Actinomycetota | 217 | 1-OH | *Nonomuraea endophytica.* |
| Ac910 | A0A5M3WAL5 | Actinomycetota | 204 | 1-OH | *Acrocarpospora corrugata.* |
| Tt028 | A0A840NWY3 | Actinomycetota | 208 | 1-OH | *Thermocatellispora tengchongensis.* |

The "unknown" represents proteins whose functions have not been identified.

**Supplementary Table 6:** Features of Unobtained Proteins in this study.

| Protein name | Uniprot ID | Phylum | Sequence length | Organism |
|---|---|---|---|---|
| Ss270 | A0A7Y0B4K4 | Actinomycetota | 232 | *Streptomyces* sp. R302. |
| Ss285 | A0A1A9DK68 | Actinomycetota | 211 | *Streptomyces* sp. OspMP-M45. |
| Sp730 | A0A345HNI9 | Actinomycetota | 232 | *Streptomyces paludis.* |
| Ss560 | A0A1Q5JM97 | Actinomycetota | 232 | *Streptomyces* sp. CB02460. |
| Ss185 | A0A8T4HQ19 | Actinomycetota | 213 | *Streptomyces* sp. A73. |
| Ss025 | A0A1I6USI8 | Actinomycetota | 208 | *Streptomyces harbinensis.* |
| As975 | W7ITJ2 | Actinomycetota | 219 | *Actinokineospora spheciospongiae.* |
| Pc498 | A0AAQ1FQ59 | Pseudomonadota | 246 | *Pseudomonas chlororaphis.* |
| Ar037 | A0A814BDH8 | Rotifera | 256 | *Adineta ricciae* |
| As592 | A0A815R559 | Rotifera | 283 | *Adineta steineri.* |
| Rs895 | A0A814MR82 | Rotifera | 225 | *Rotaria sordida.* |
| Mb230 | A0A956FFW7 | Myxococcota | 208 | *Myxococcales bacterium.* |
| Cs030 | A0A2W6YVX3 | Pseudomonadota | 211 | *Chelatococcus* sp. |
| Me615 | A0A2P7S268 | Pseudomonadota | 212 | *Mesorhizobium ephedrae.* |
| Hb155 | A0A4Q3FEH4 | Pseudomonadota | 211 | *Hyphomicrobiales bacterium.* |
| Bc925 | A0A370L000 | Pseudomonadota | 211 | *Bosea caraganae.* |
| Ny655 | A0A386ZP08 | Actinomycetota | 225 | *Nocardia yunnanensis.* |
| Ag690 | A0A9W6QMN8 | Actinomycetota | 225 | *Actinokineospora globicatena.* |
| Ar885 | A0A1I5RRH4 | Actinomycetota | 236 | *Amycolatopsis rubida.* |

**Supplementary Table 7:** $^1$H (600 MHz) and $^{13}$C (150 MHz) NMR Data of **1** in DMSO-d*6*.

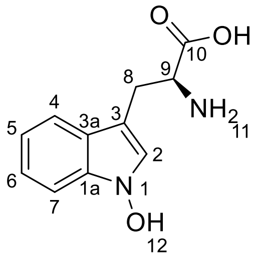

| Compound **1** | | |
|---|---|---|
| Position | $\delta_C$ | $\delta_H$, *J* (Hz) |
| 1 | | |
| 1a | 133.35 | |
| 2 | 124.87 | 7.30, *s* |
| 3 | 103.94 | |

| | | |
|---|---|---|
| 3a | 123.28 | |
| 4 | 118.60 | 7.56, *d*, (7.9), |
| 5 | 118.25 | 6.96 *t*, (7.4) |
| 6 | 121.18 | 7.11 *t*, (7.6) |
| 7 | 108.38 | 7.34 *d*, (8.1) |
| 8 | 26.58 | 3.23 *dd*, (15.1, 4.4); 3.00 *dd*, (15.1, 7.9) |
| 9 | 54.54 | 3.55, *t*, (6.1) |
| 10 | 170.82 | |

**Supplementary Table 8:** $^{1}$H (600 MHz) and $^{13}$C (150 MHz) NMR Data of **2** in DMSO-d*6*.

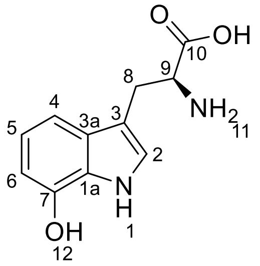

| Compound **3** | | |
|---|---|---|
| Position | *δ*C | *δ*H, *J* (Hz) |
| 1 | | 11.02, *s* |
| 1a | 126.23 | |
| 2 | 123.68 | 7.08, *s* |
| 3 | 109.27 | |
| 3a | 129.23 | |
| 4 | 105.73 | 7.02, *d*, (7.8) |
| 5 | 119.17 | 6.77, *t*, (7.6) |
| 6 | 109.30 | 6.49, *d*, (7.5) |
| 7 | 143.83 | |
| 8 | 27.23 | 3.27, *dd*, (14.9, 4.3); 2.99, *dd*, (15.0, 8.6) |
| 9 | 54.62 | 3.57, *dd*, (8.8, 4.2) |
| 10 | 171.17 | |

**Supplementary Table 9:** $^{1}$H (600 MHz) and $^{13}$C (150 MHz) NMR Data of *S*-**4** in $CD_3OD$.

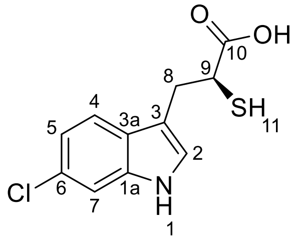

| Compound **4** | | |
|---|---|---|
| Position | *δ*C | *δ*H, *J* (Hz) |
| 1 | | |
| 1a | 138.23 | |

| | | |
|---|---|---|
| 2 | 125.37 | 7.07 (s, 1H) |
| 3 | 112.95 | |
| 3a | 127.17 | |
| 4 | 120.35 | 7.46 (d, *J* = 8.5 Hz, 1H) |
| 5 | 120.29 | 6.94, dd, (8.5, 1.9) |
| 6 | 128.29 | |
| 7 | 112.03 | 7.29, d, (1.8) |
| 8 | 32.86 | 3.07, *m*; 3.29, *m* |
| 9 | 42.77 | 3.61, dd, (8.8, 6.2) |
| 10 | 176.89 | |

**Supplementary Table 10:** $^{1}$H (600 MHz) and $^{13}$C (150 MHz) NMR Data of **5** in DMSO-d*6*.

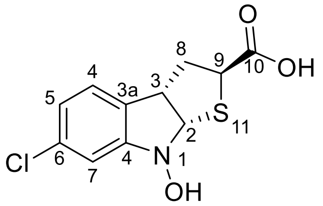

| Compound **5** | | |
|---|---|---|
| Position | $\delta_C$ | $\delta_H$, *J* (Hz) |
| 1 | | |
| 1a | 153.49 | |
| 2 | 83.55 | 5.66, *d*, (5.9) |
| 3 | 47.88 | 3.94, *td*, (6.0, 2.6) |
| 3a | 127.61 | |
| 4 | 124.83 | 7.23, *dd*, (7.9, 1.1) |
| 5 | 120.74 | 6.90, *dd*, (7.8, 2.0) |
| 6 | 132.67 | |
| 7 | 111.82 | 6.73, *s* |
| 8 | 37.23 | 2.65, *m*; 2.41, *m* |
| 9 | 48.33 | 3.65, *d*, (4.2) |
| 10 | 172.51 | |

**Supplementary Table 11:** Data collection and refinement statistics of Mc170.

| | Mc170 + L-Trp |
|---|---|
| **PDB code** | 9X39 |
| **Data collection** | |
| Space Group | P 3 2 1 |
| Cell dimensions | |
| *a, b, c* (Å) | 72.59,72.59,81.66 |
| a, b, g (°) | 90.00,90.00,120.00 |

| | |
|---|---|
| Resolution (Å) | 81.66-2.54(2.63-2.54)* |
| *R*sym or *R*merge | 6.7(21.7) |
| CC1/2 | 0.989(0.485) |
| $I/\delta I$ | 10.0(2.1) |
| Completeness (%) | 100.0(100.0) |
| Redundancy | 31.6(27.4) |
| **Refinement** | |
| Resolution (Å) | 62.86-2.544 (2.635-2.544) |
| Completeness (%) | 99.71(100.00) |
| No. reflections | 8512 (848) |
| $R_{work}/R_{free}$ | 23.8/30.7 |
| No. atoms | |
| Protein | 1455 |
| Ligand/ion | 69 |
| Water | 16 |
| B-factors | |
| Protein | 26.69 |
| Ligand/Ion | 20.63 |
| Water | 29.98 |
| R.m.s. deviations | |
| Bond lengths (Å) | 0.012 |
| Bond angles (°) | 1.54 |

One crystal was used for each structure.

*Values in parentheses are for highest-resolution shell.

**Supplementary Figures 1-40**

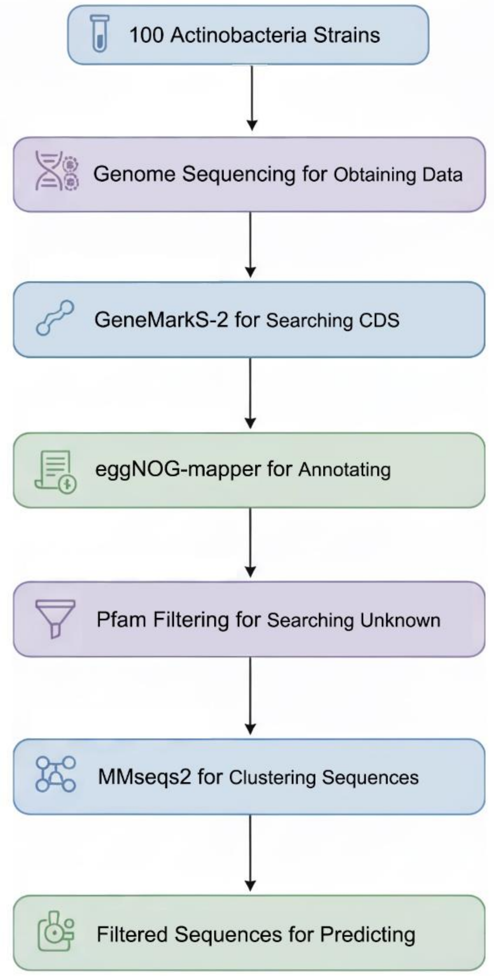

**Supplementary Fig. 1:** Dataset processing of 100 actinobacteria strains for large-scale Heme-binding protein screening.

a

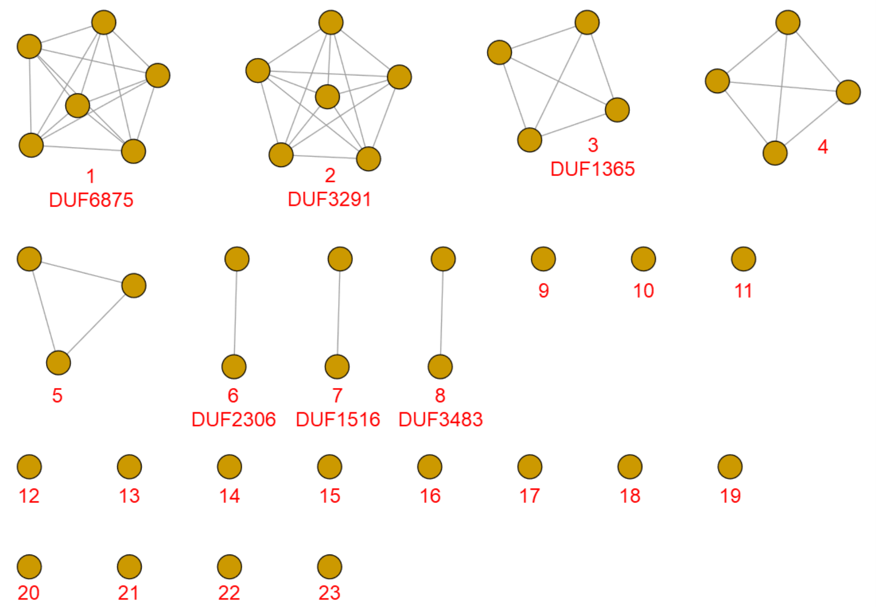

b

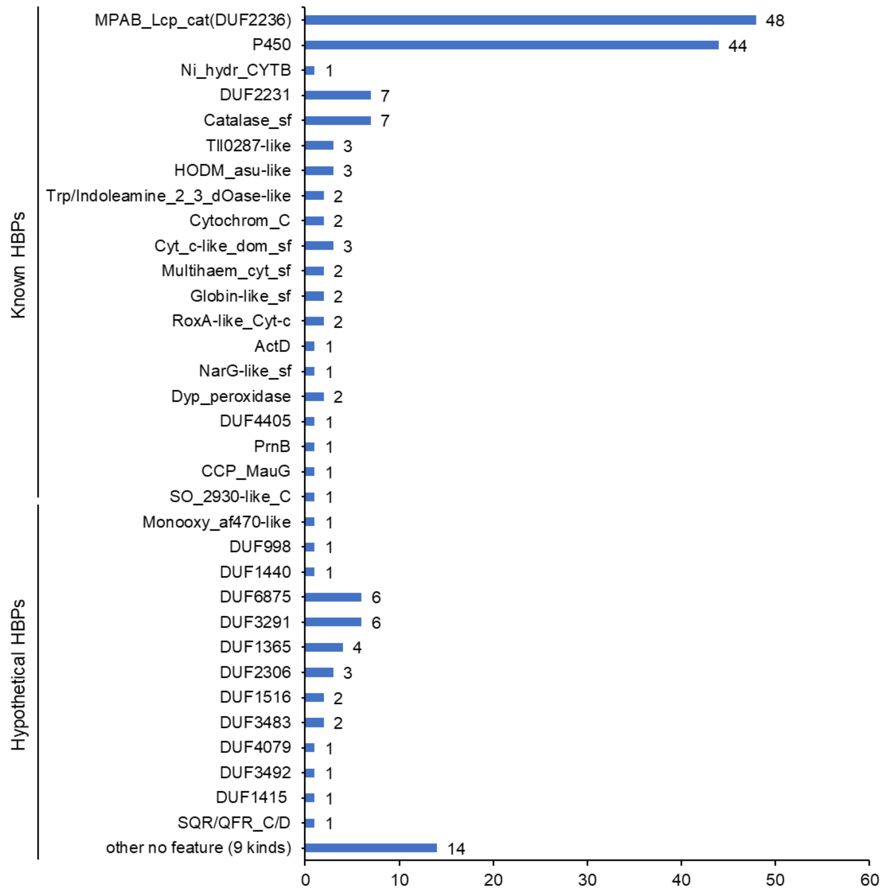

**Supplementary Fig. 2 | Identification and characterization of heme-binding proteins through CISSpector screening and sequence analysis.**

**a,** CISSpector screening identified 44 hypothetical heme-binding proteins (HBPs) from 100 deepsea actinomycete proteomes. Sequence similarity network (SSN) analysis of these sequences using the EFI-EST[1,2], constructed at a 25% identity cutoff, grouped them into 23 clusters.

**b,** Bar plot displaying the distribution of the 43 HBP families. Functional annotation based on UniProt family and domain information revealed that approximately 24.6% of clusters belong to the DUF2236 family (Pfam: MPAB_Lcp_cat), which includes biochemically characterized members such as LCPK30 from *Streptomyces* sp. K3011 and endoplasmic reticulum oxygenase MpaB′ from *Penicillium brevicompactum*. Cytochrome P450s constitute about 23% of clusters, with other known families including catalases, tryptophan 2,3-dioxygenases (TDOs), hemoglobins, cytochromes *c*, and MauG. Full annotations are available in Supplementary Table 3.

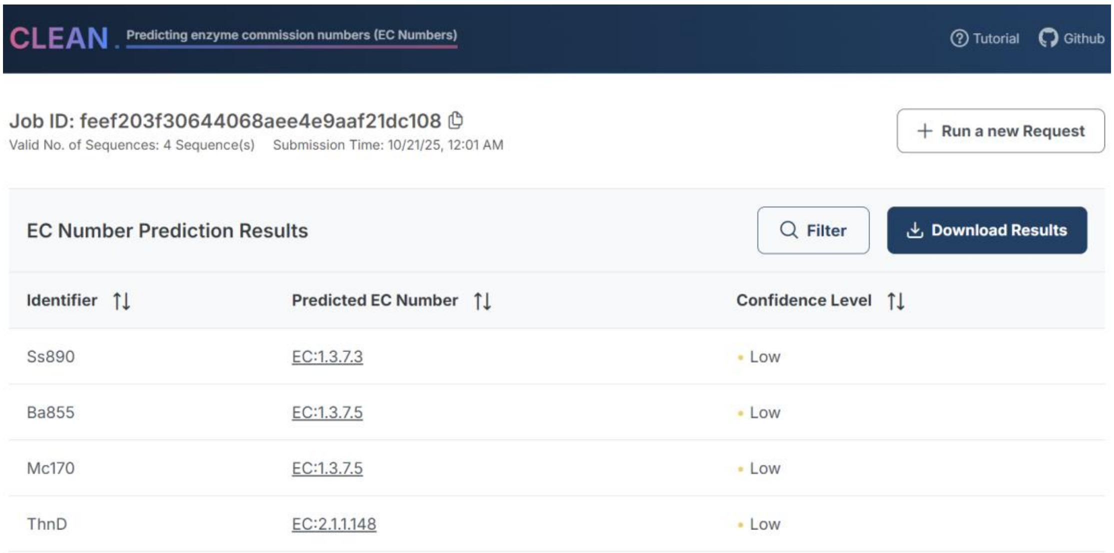

**Supplementary Fig. 3 | CLEAN-based functional prediction for cytochrome P422 family members.** Computational annotation of four representative cytochrome P422 enzymes using the CLEAN online server resulted in assignment as ferredoxin oxidoreductase (EC 1.3.7.3 and EC 1.3.7.5) or thymidylate synthase (EC 2.1.1.148), which contrasts with their experimentally verified activities.

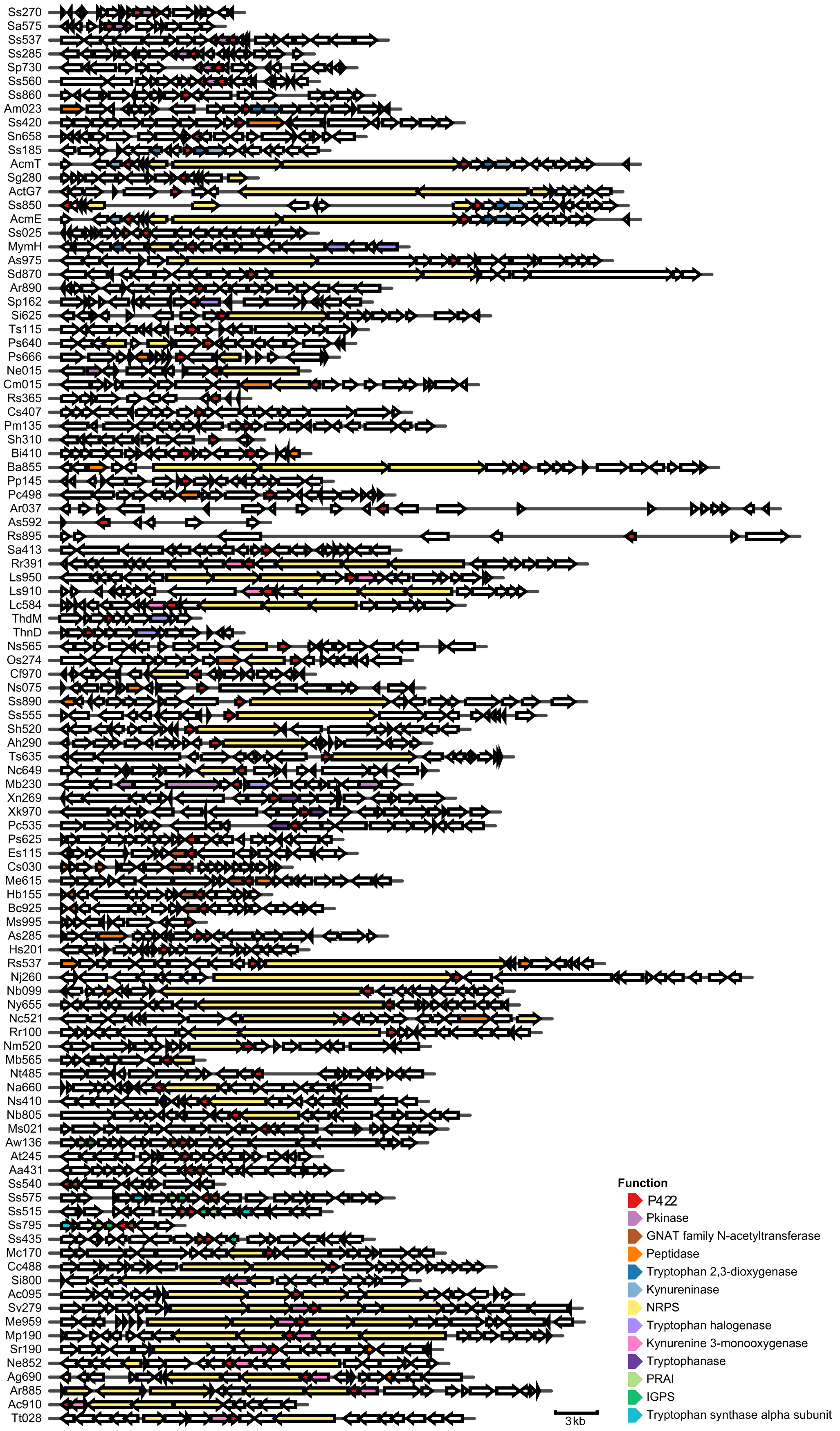

**Supplementary Fig. 4: Genomic context analysis of cytochrome P422 members.** Approximately 49% of cytochrome P422 genes reside within non-ribosomal peptide synthetase (NRPS)

biosynthetic gene clusters. Cytochrome P422 commonly clusters genomically with tryptophan metabolic enzymes, including tryptophan 2,3-dioxygenase (TDO), kynureninase, tryptophanase, phosphoribosylanthranilate isomerase (PRAI), and indole-3-glycerol phosphate synthase (IGPS). In other cases, members co-localize with protein kinases (Pkinases) and GNAT family *N*-acetyltransferases, potentially implicating their roles in specialized metabolite biosynthesis. Separately, co-localization with peptidases suggests possible peptide-modifying functions.

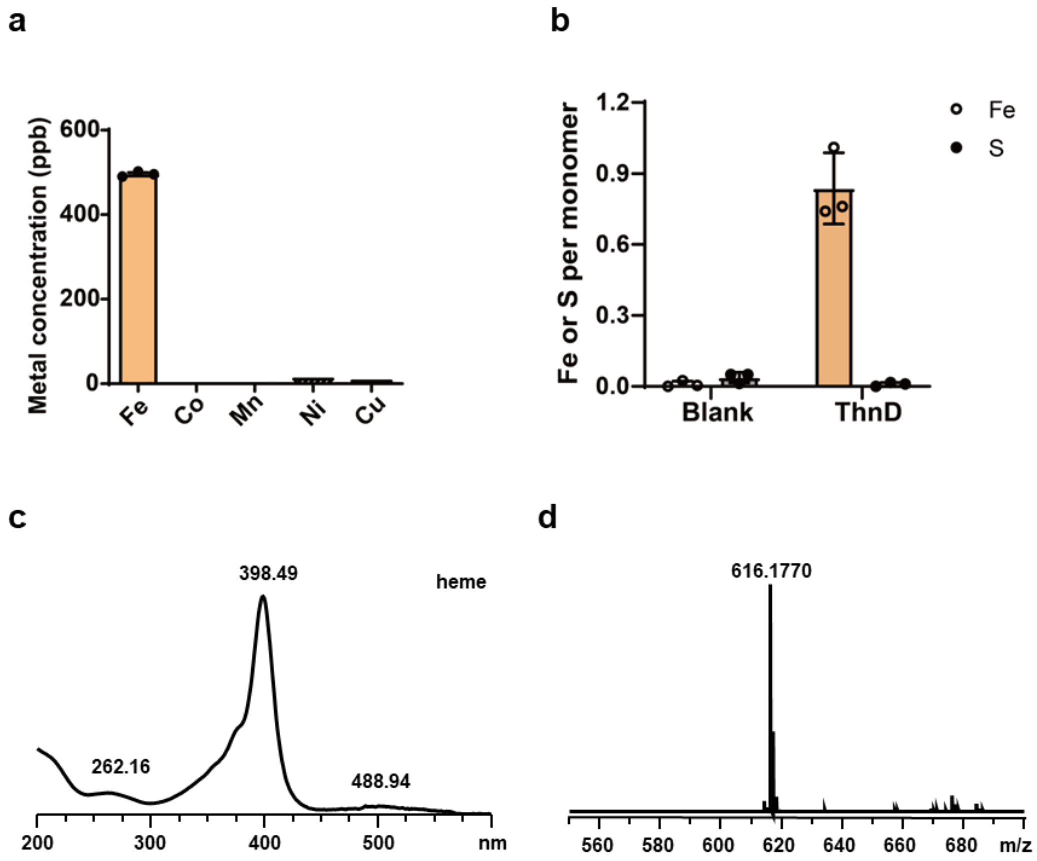

**Supplementary Fig. 5: Cofactor identification in ThnD. a**, Metal analysis of recombinant ThnD by ICP-MS. Data represent mean ± SEM (n = 3). **b**, Quantification of iron (hollow bars) and sulfur (solid bars) content in ThnD. Values are mean ± SEM (n = 3). **c**, UV-visible absorption spectrum of heme cofactor extracted from ThnD. **d**, Mass spectrometric identification of heme in ThnD. The heat-denatured protein supernatant was analyzed by UV- visible spectroscopy and LC-MS. The observed *m/z* 616.1770 matches the heme molecular mass ($[M+H]^+$) consistent with tryptophan hydroxylase cofactors[3].

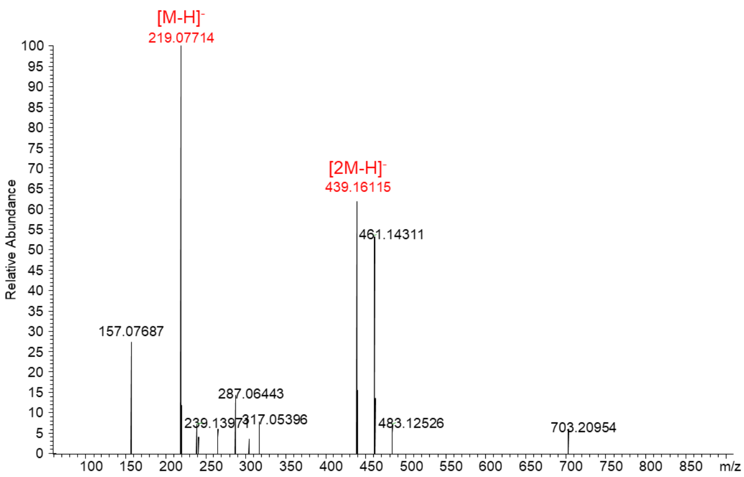

**Supplementary Fig. 6:** Negative-ion HR-MS profiles of compound **1**.

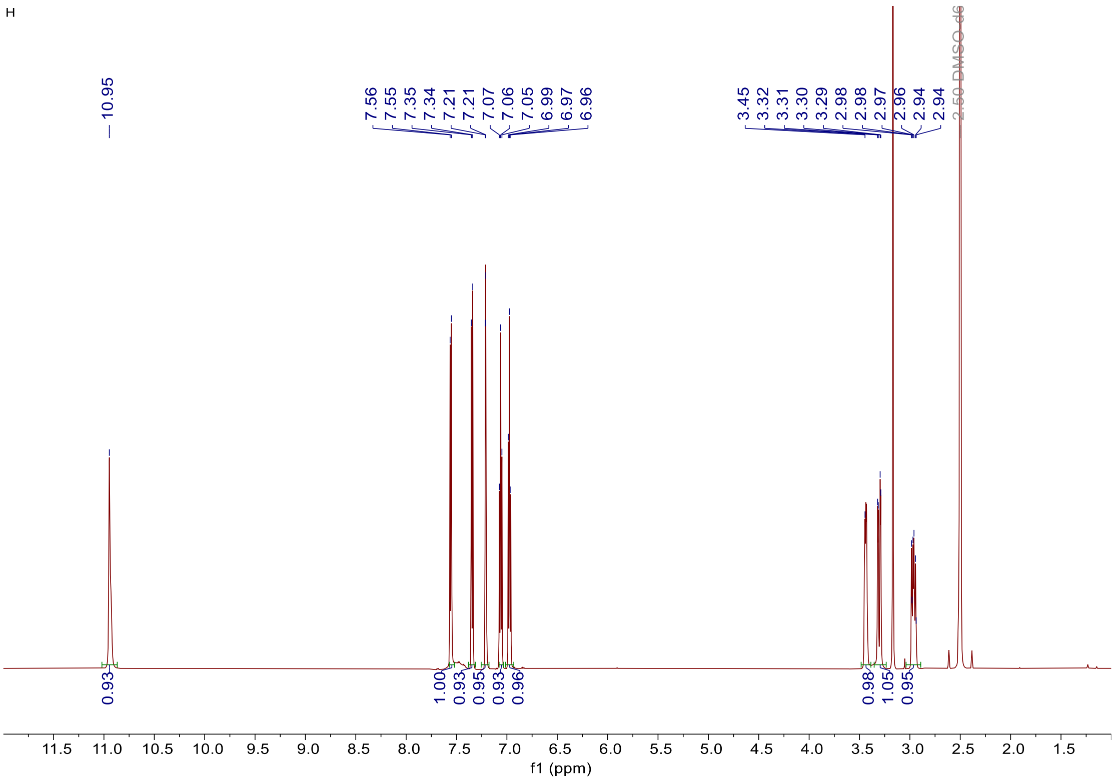

**Supplementary Fig. 7:** The $^{1}$H NMR spectrum of L-Trp in DMSO-d*6*.

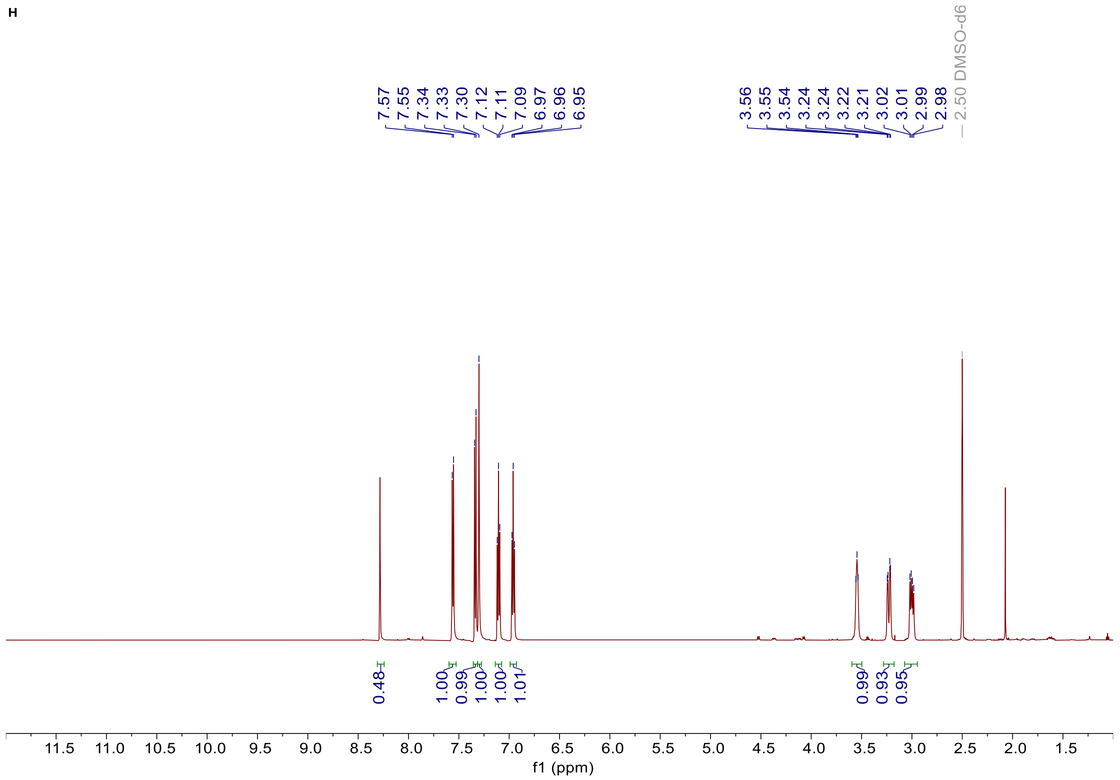

**Supplementary Fig. 8:** The $^{1}$H NMR spectrum of compound **1** in DMSO-d*6*.

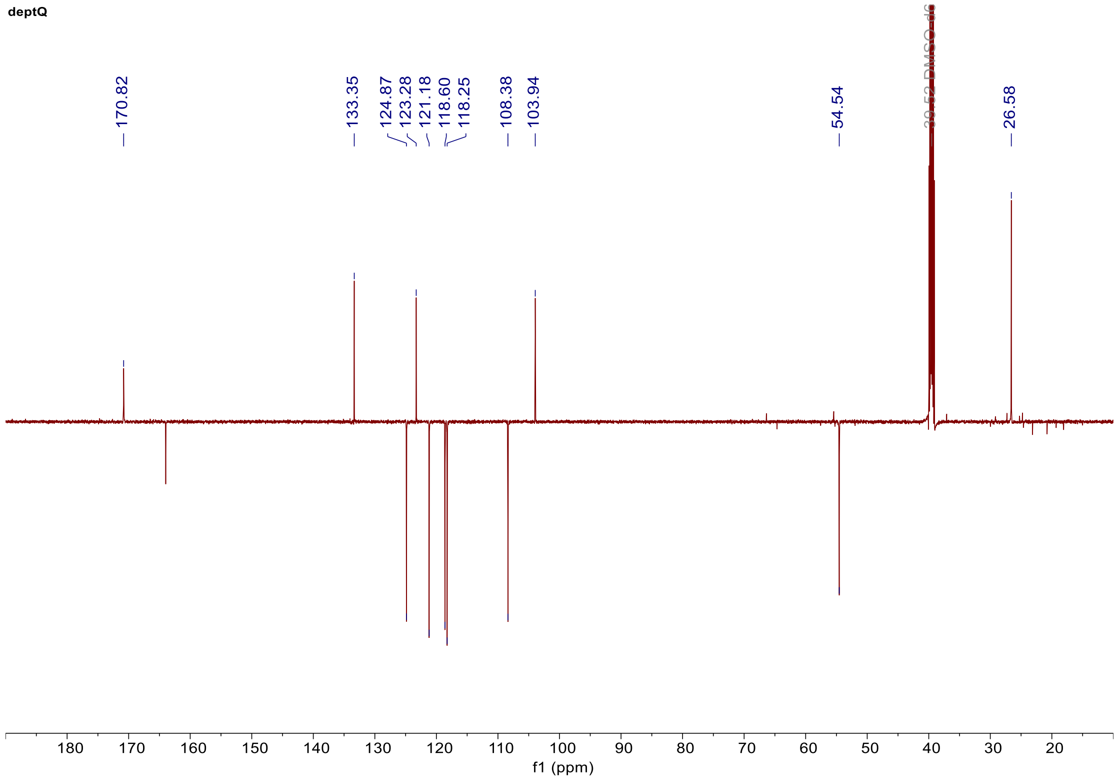

**Supplementary Fig. 9:** The $^{13}$C-NMR spectrum (DEPT-Q) of compound **1** in DMSO-d*6*.

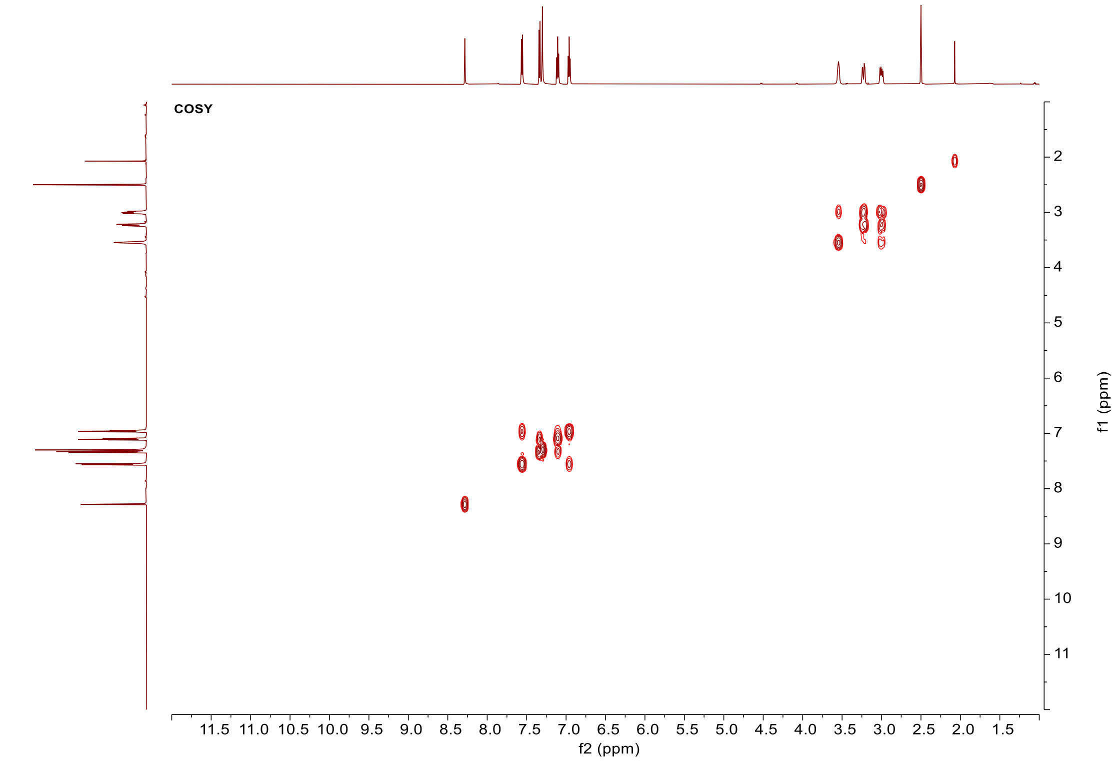

**Supplementary Fig. 10:** The $^{1}$H-$^{1}$H-COSY spectrum of compound **1** in DMSO-d*6*.

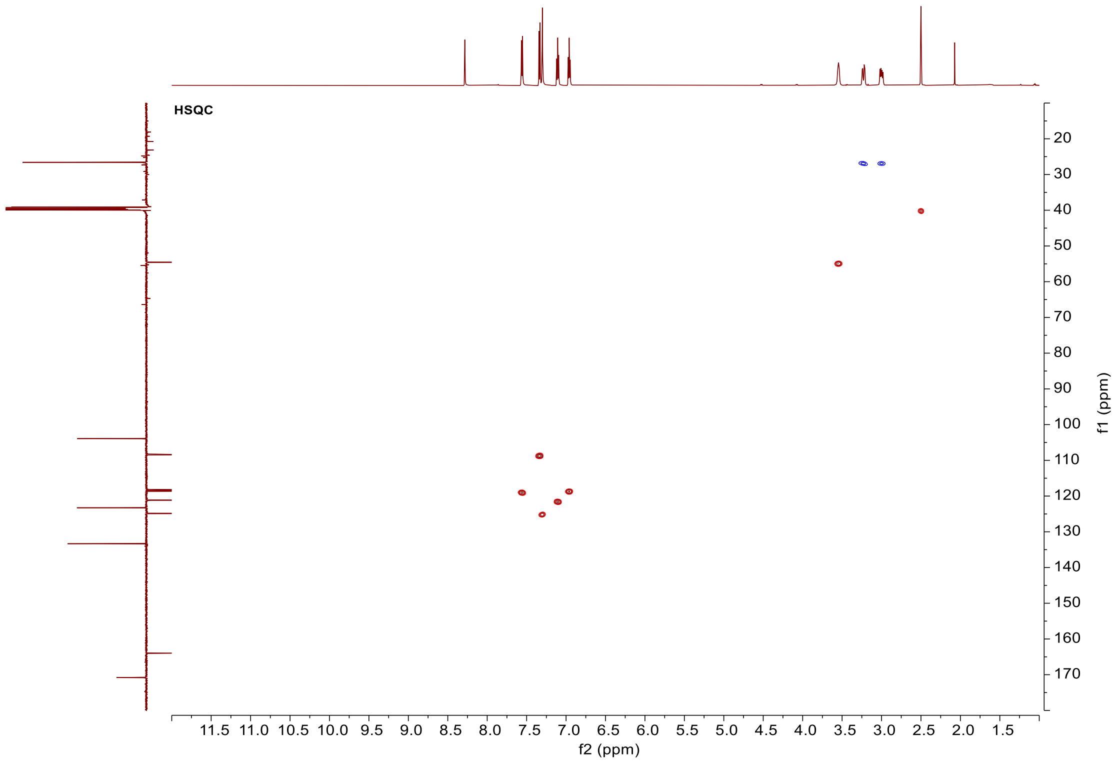

**Supplementary Fig. 11:** The $^{1}$H-$^{13}$C HSQC spectrum of compound **1** in DMSO-d*6*.

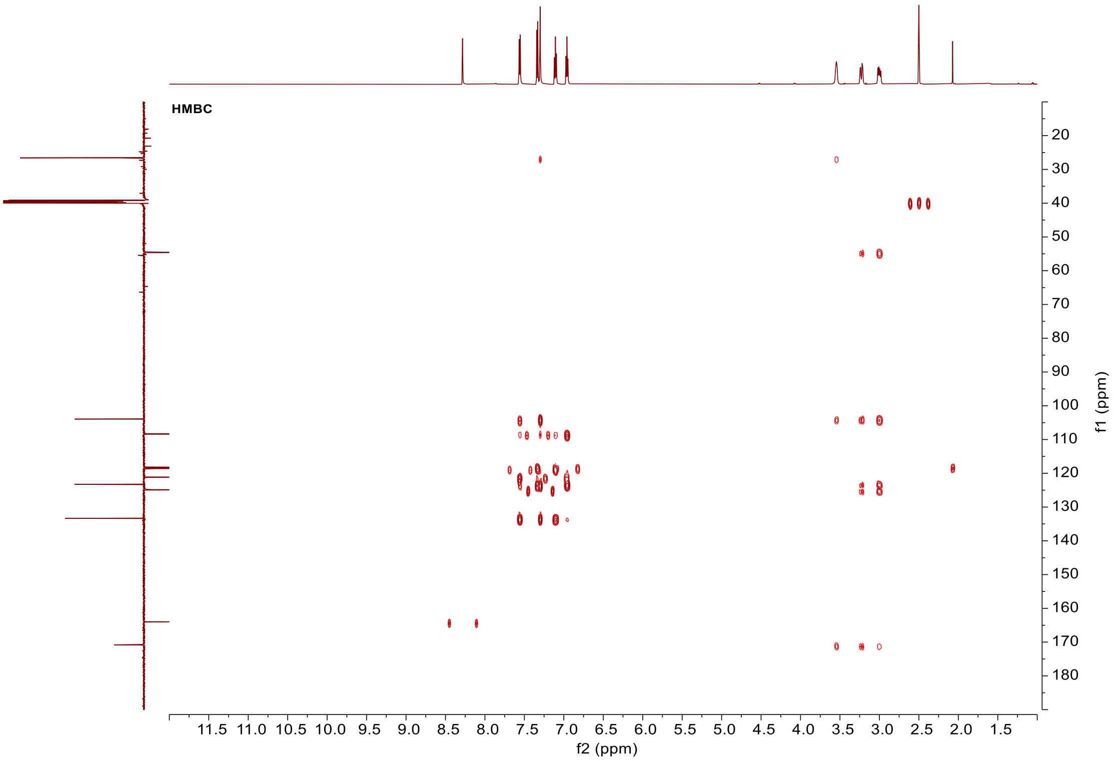

**Supplementary Fig. 12:** The $^{1}H$-$^{13}C$ HMBC spectrum of compound **1** in DMSO-d*6*.

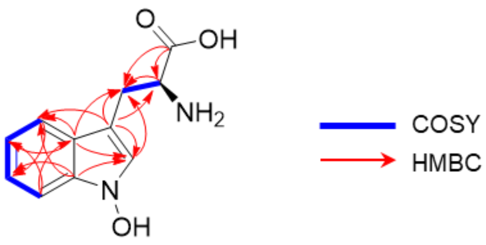

**1**

**Supplementary Fig. 13:** Key $^{1}H$-$^{1}H$ COSY and $^{1}H$-$^{13}C$ HMBC correlations of compound **1**.

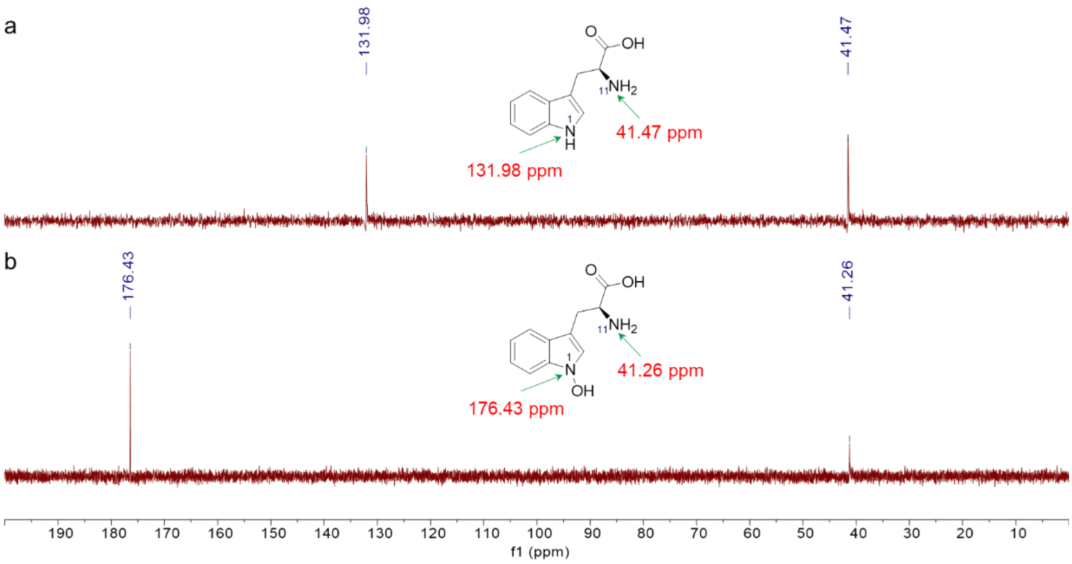

**Supplementary Fig. 14:** The $^{15}$1-OHMR spectrum of L-Trp (a) and compound **1** (b) in DMSO-d*6*. The $^{15}$1-OHMR analysis of $^{15}$N-labeled L-Trp and $^{15}$N-labeled compound **1** revealed a downfield shift of 44.45 ppm for N1 in compound **1** (relative to the resonance at 131.98 ppm in L-Trp). In contrast, the chemical shift of N11 differed by only 0.21 ppm between L-Trp ($\delta_{N11}$ = 41.47 ppm) and compound 1 ($\delta_{N11}$ = 41.26 ppm). Consequently, oxygenation is more likely to take place at N1 rather than N11, thereby generating the 1-OH-Trp.

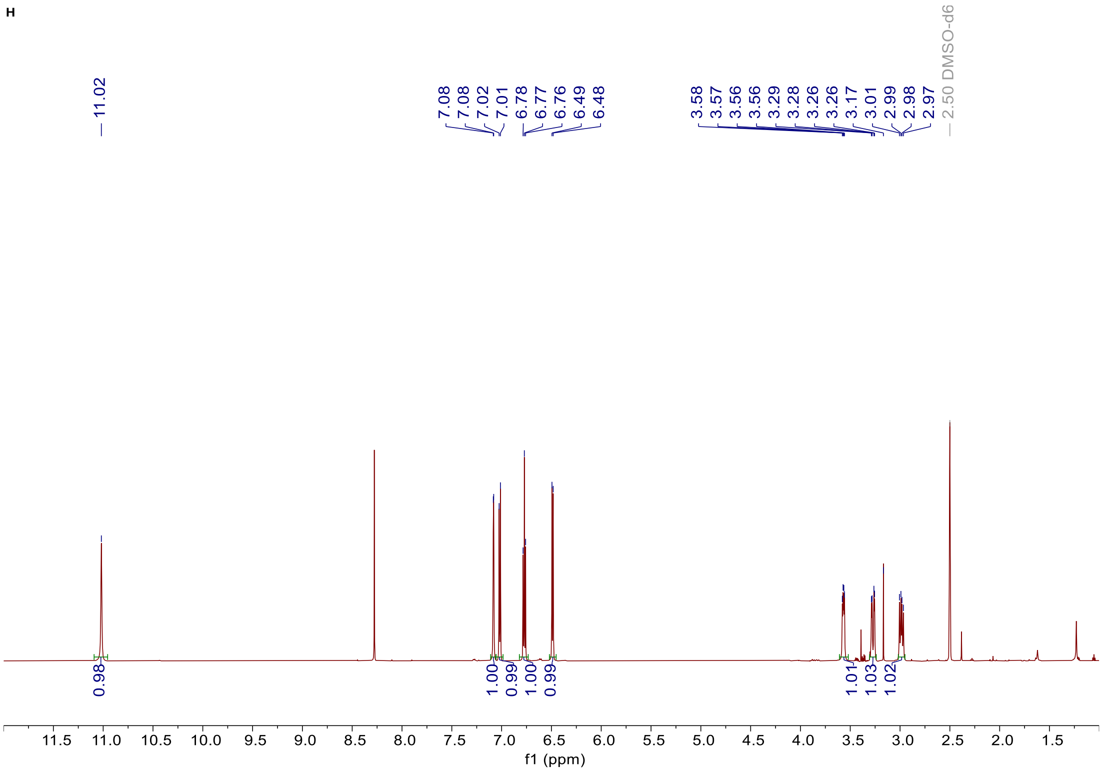

**Supplementary Fig. 15:** The $^{1}$H NMR spectrum of compound **2** in DMSO-d*6*.

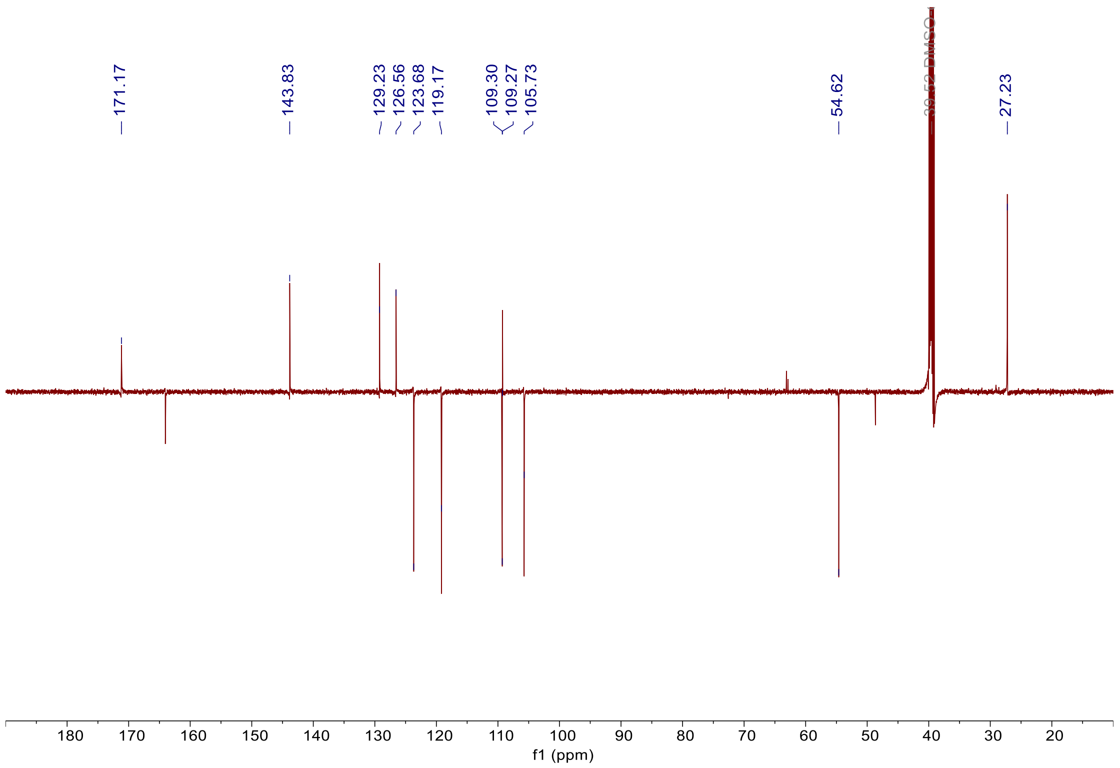

**Supplementary Fig. 16:** The $^{13}$C-NMR spectrum (DEPT-Q) of compound **2** in DMSO-d*6*.

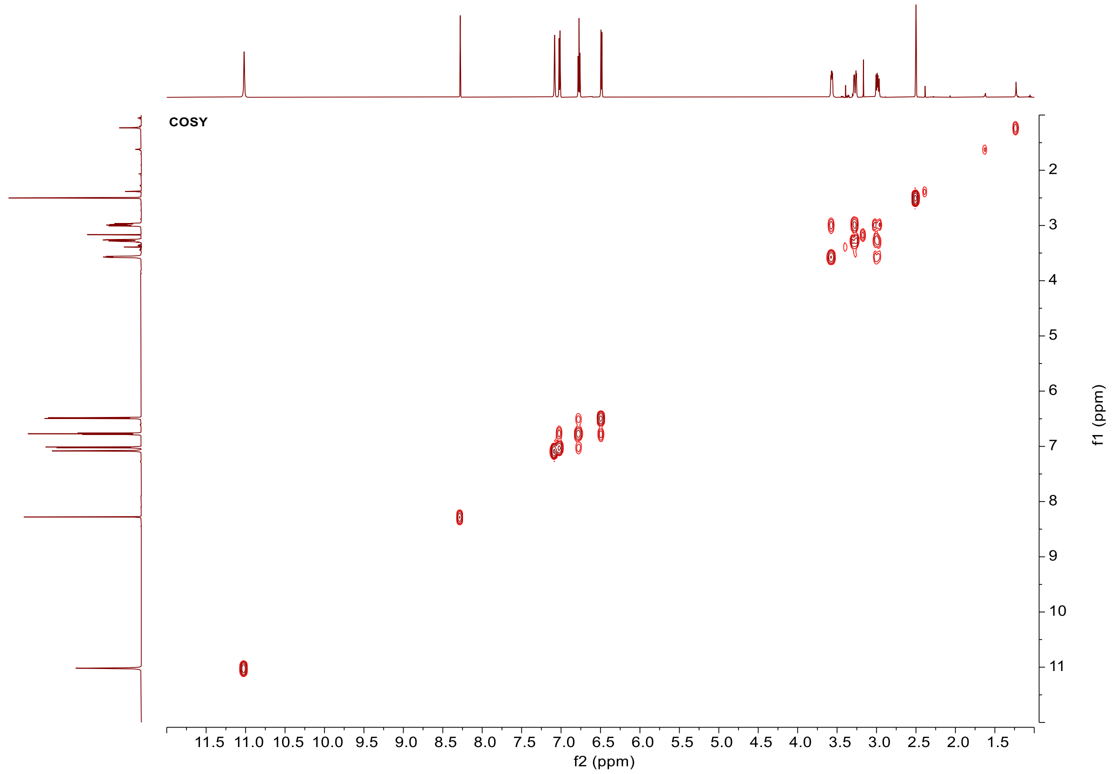

**Supplementary Fig. 17:** The $^{1}$H-$^{1}$H-COSY spectrum of compound **2** in DMSO-d*6*.

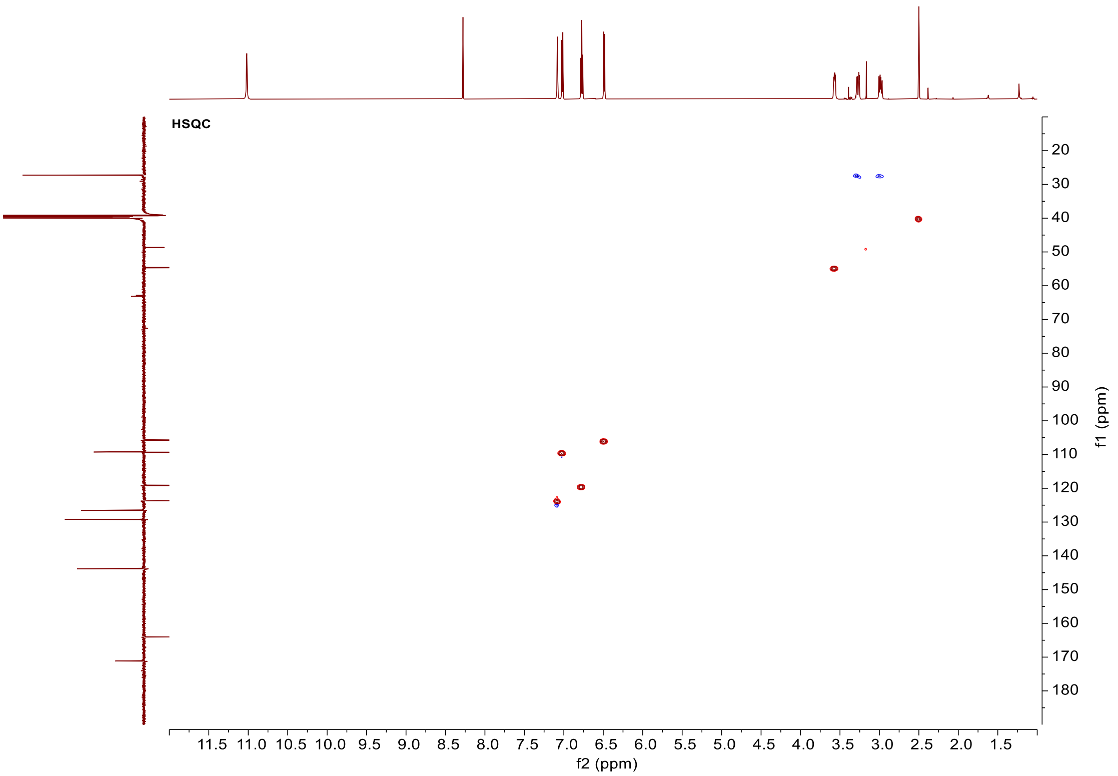

**Supplementary Fig. 18:** The $^{1}$H-$^{13}$C HSQC spectrum of compound **2** in DMSO-d*6*.

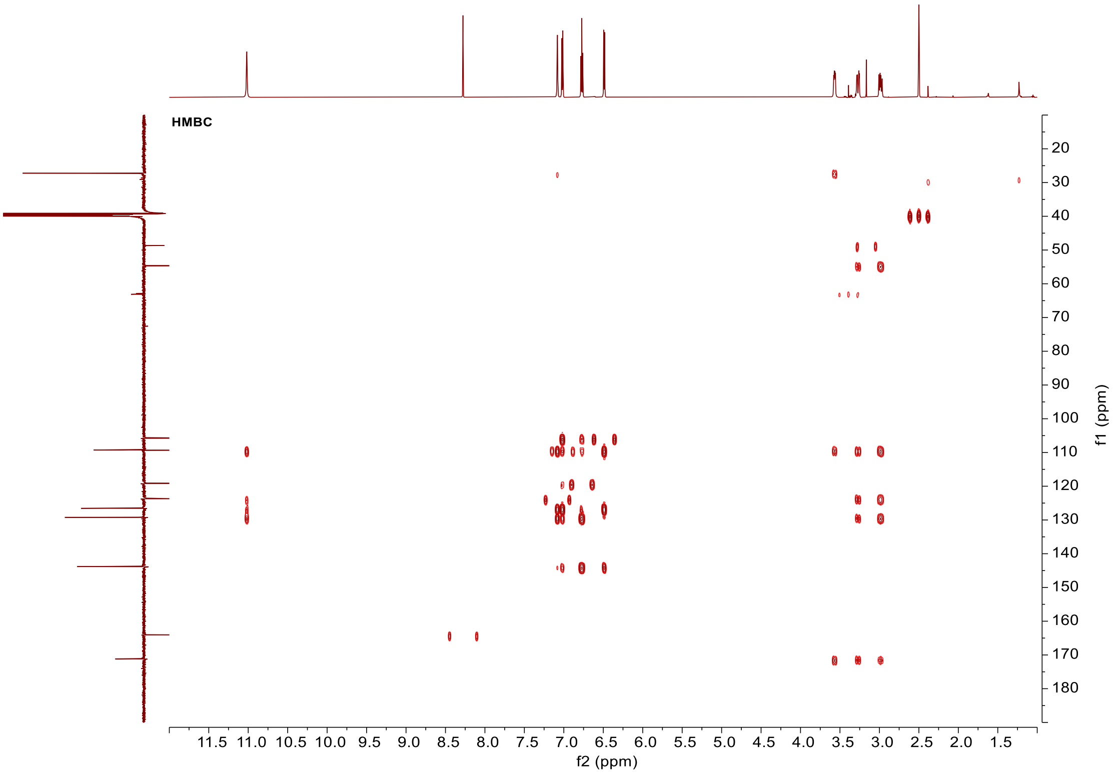

**Supplementary Fig. 19:** The $^{1}$H-$^{13}$C HMBC spectrum of compound **2** in DMSO-d*6*.

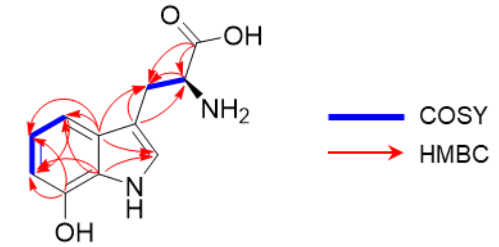

**Supplementary Fig. 20:** Key [1]H-[1]H COSY and [1]H-[13]C HMBC correlations of compound **2**.

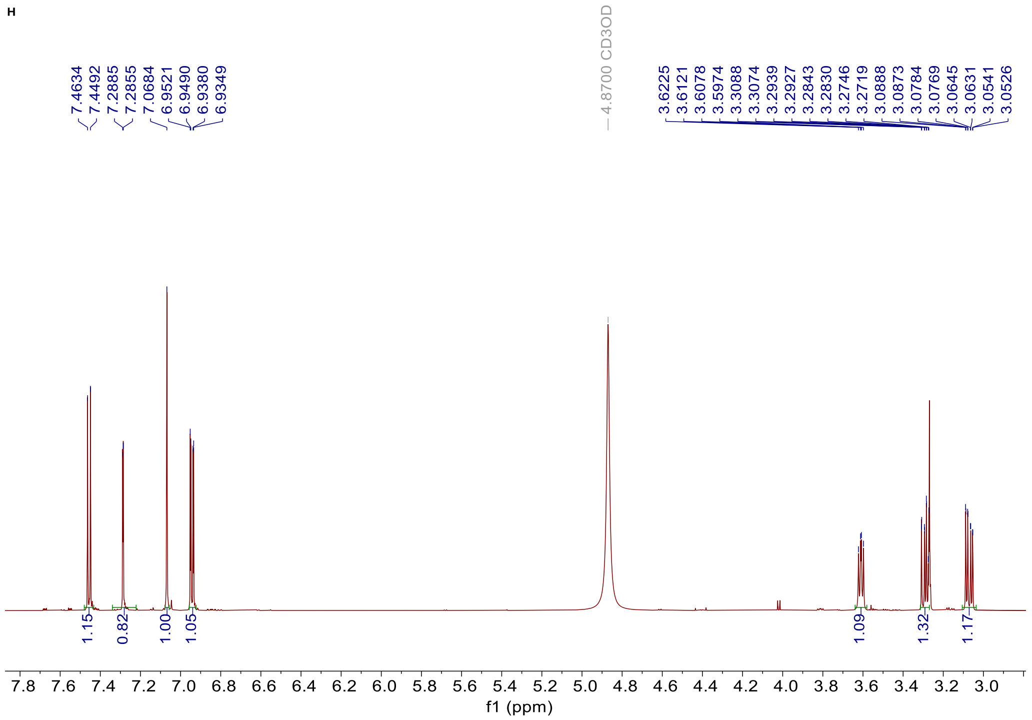

**Supplementary Fig. 21:** The [1]H NMR spectrum of compound **4** in $CD_3OD$.

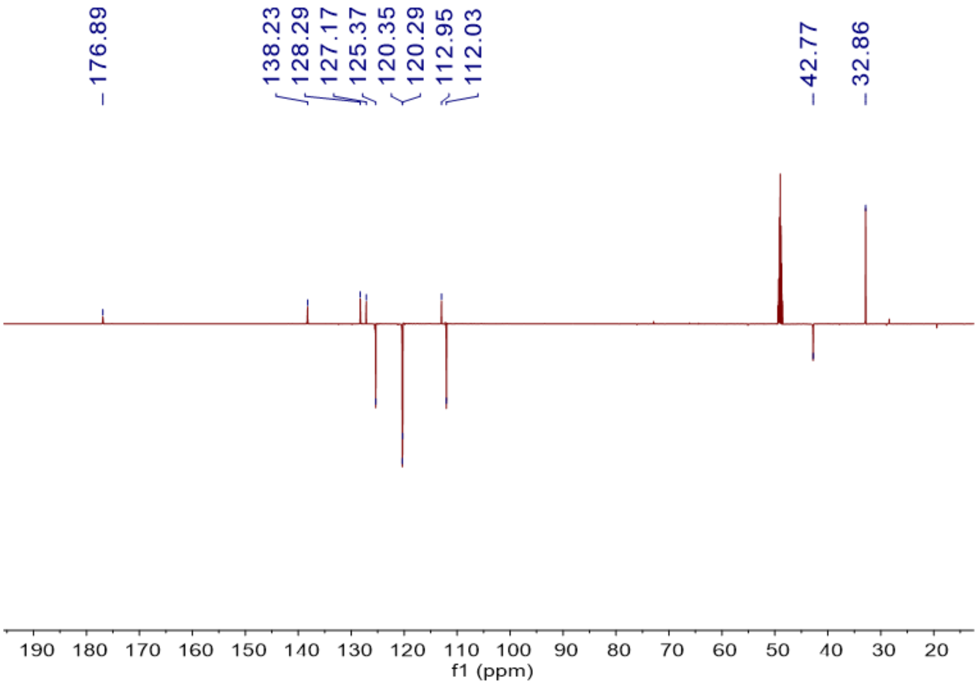

**Supplementary Fig. 22:** The $^{13}C$-NMR spectrum (DEPT-Q) of compound **4** in $CD_3OD$.

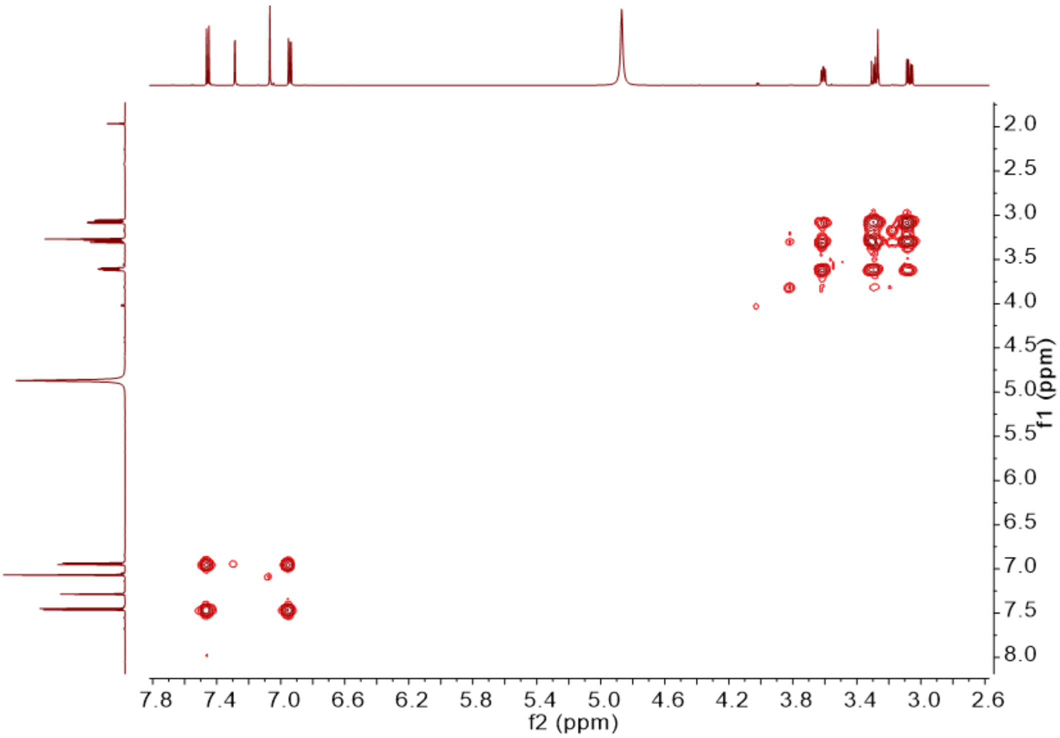

**Supplementary Fig. 23:** The $^{1}H$-$^{1}H$-COSY spectrum of compound **4** in $CD_3OD$.

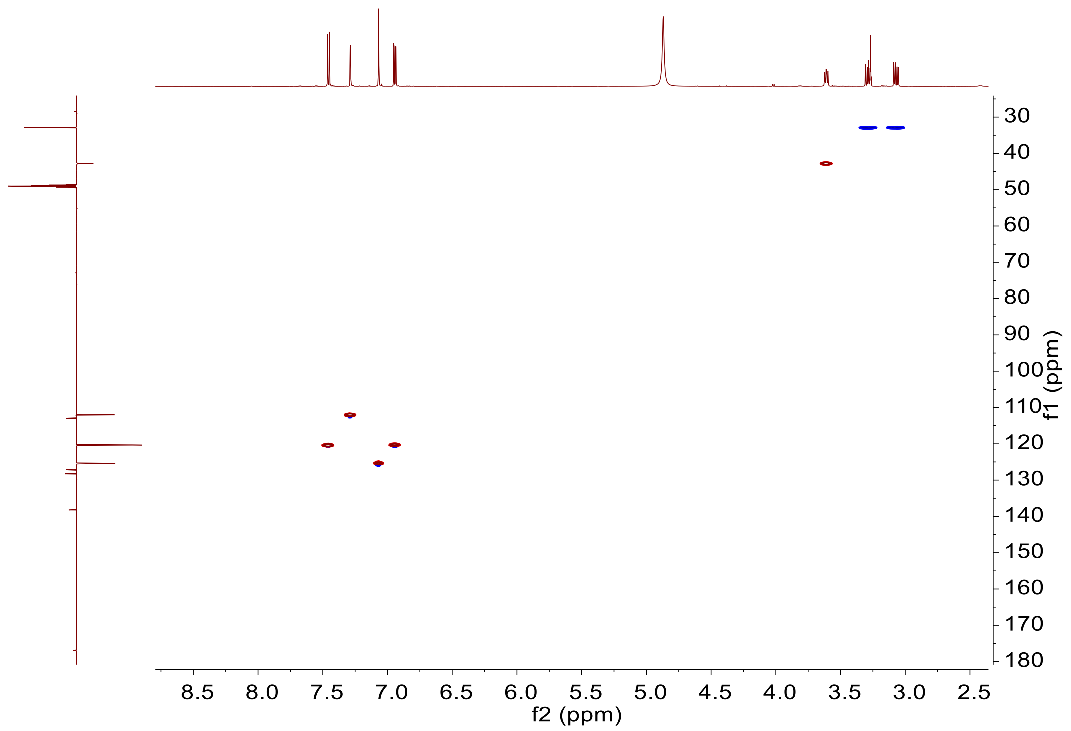

**Supplementary Fig. 24:** The [1]H-[13]C HSQC spectrum of compound **4** in $CD_3OD$.

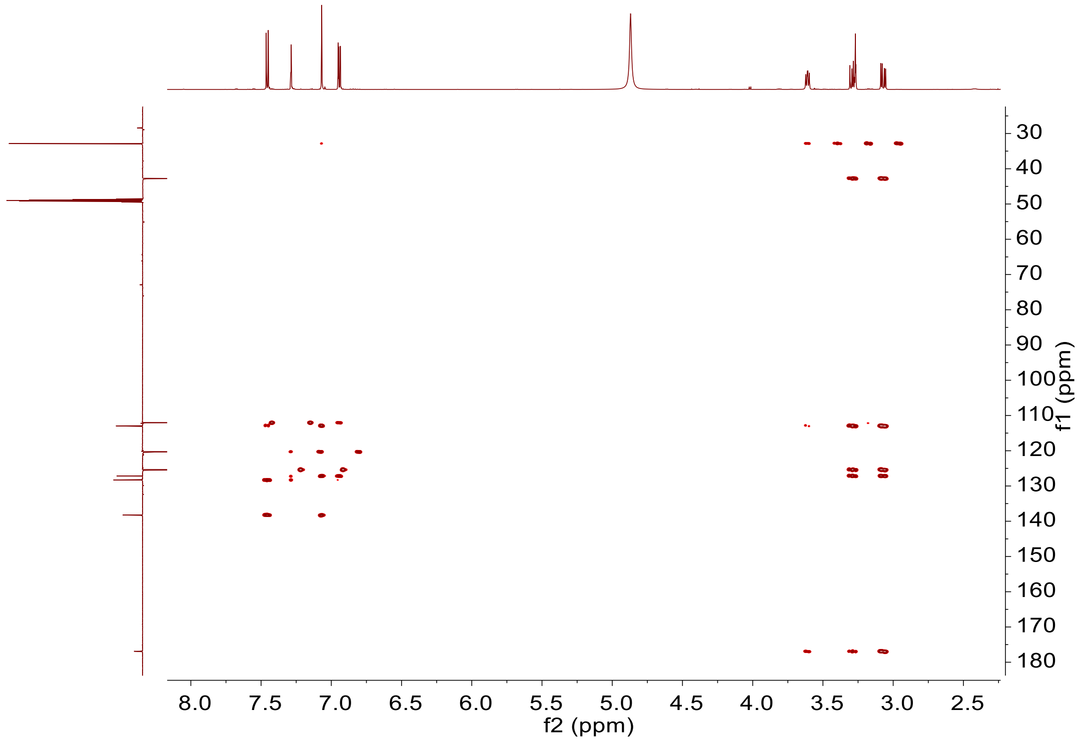

**Supplementary Fig. 25:** The [1]H-[13]C HMBC spectrum of compound **4** in $CD_3OD$.

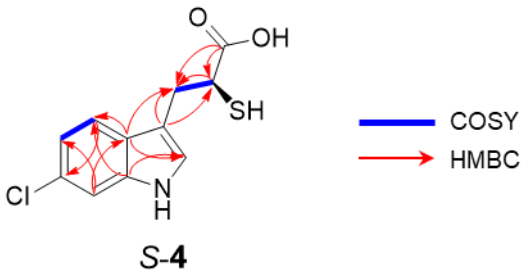

**Supplementary Fig. 26:** Key $^1H$-$^1H$ COSY and $^1H$-$^{13}C$ HMBC correlations of compound **4**.

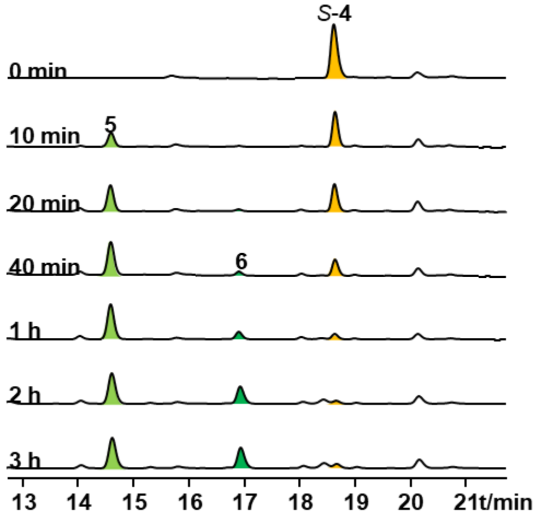

**Supplementary Fig. 27: Time-course analysis of ThnD enzymatic activity.** Reactions containing ThnD (20 µM), substrate *S*-**4** (500 µM), electron transfer system (40 µM *Sel*Fdx1499, 20 µM *Sel*FdR0978), NADPH (10 mM), and DTT (10 mM) were sampled at indicated time points (10, 20, 40 min; 1, 2, 3 h). Samples were immediately quenched with equal volume of methanol, vortex-mixed, and centrifuged (14,000 × g, 10 min). The resulting supernatants were collected for analysis.

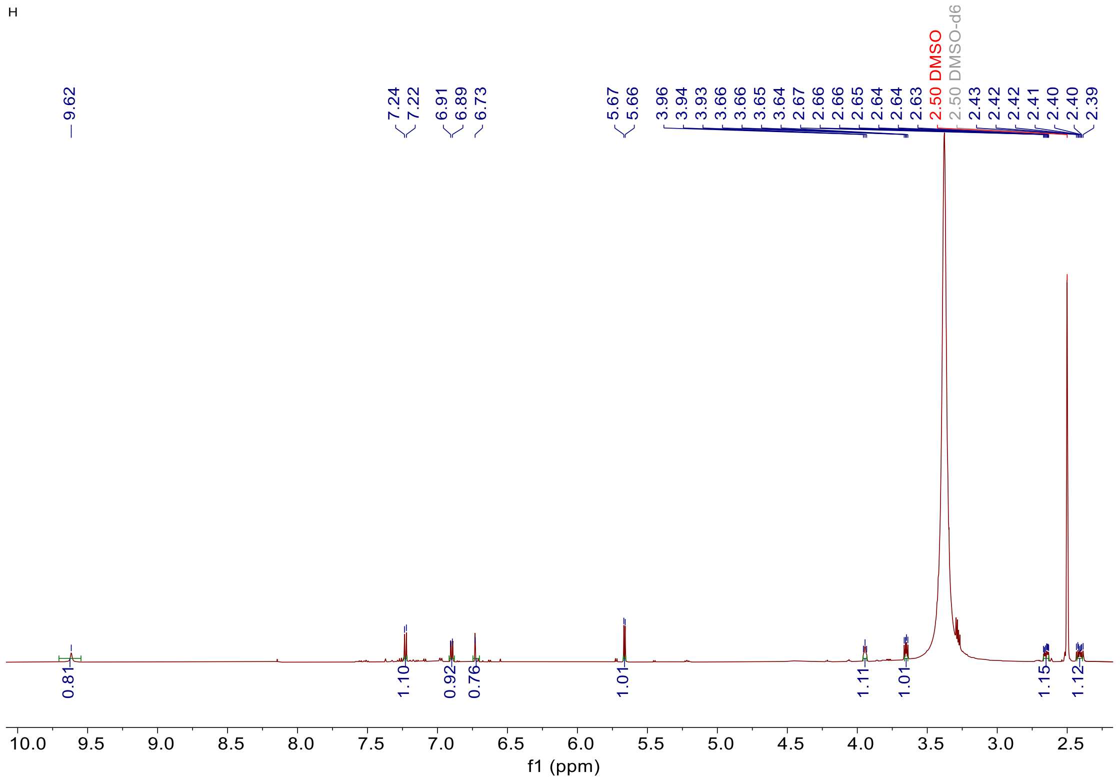

**Supplementary Fig. 28:** The $^{1}H$ NMR spectrum of compound **5** in DMSO-d*6*.

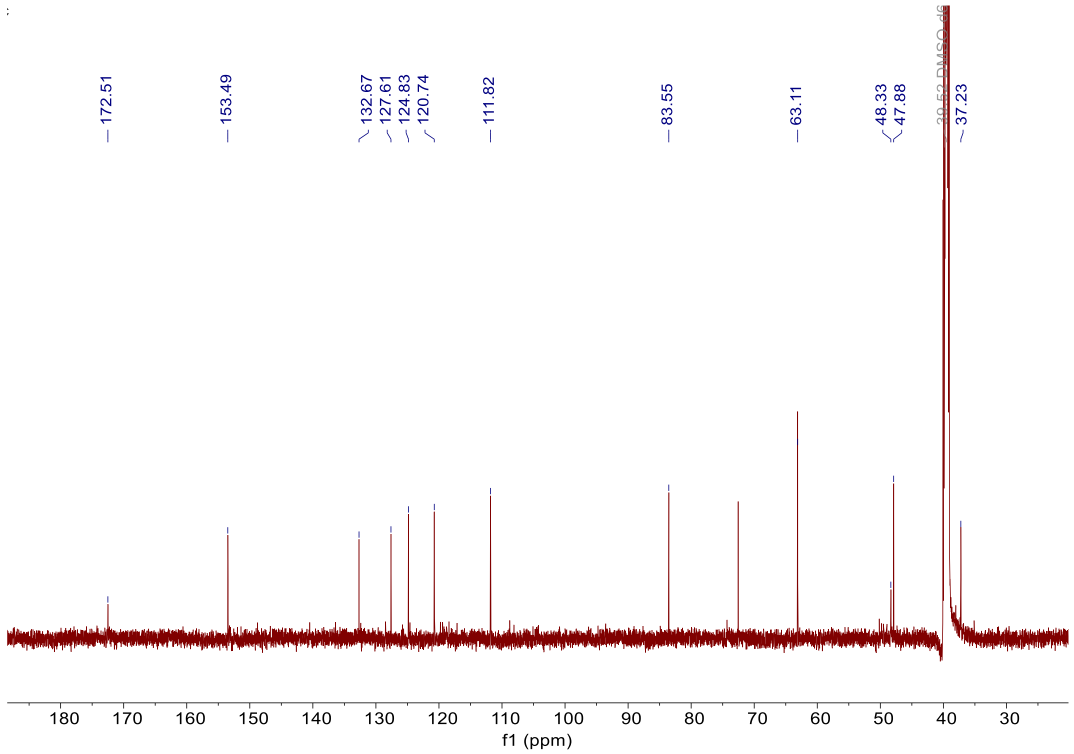

**Supplementary Fig. 29:** The $^{13}C$-NMR spectrum of compound **5** in DMSO-d*6*.

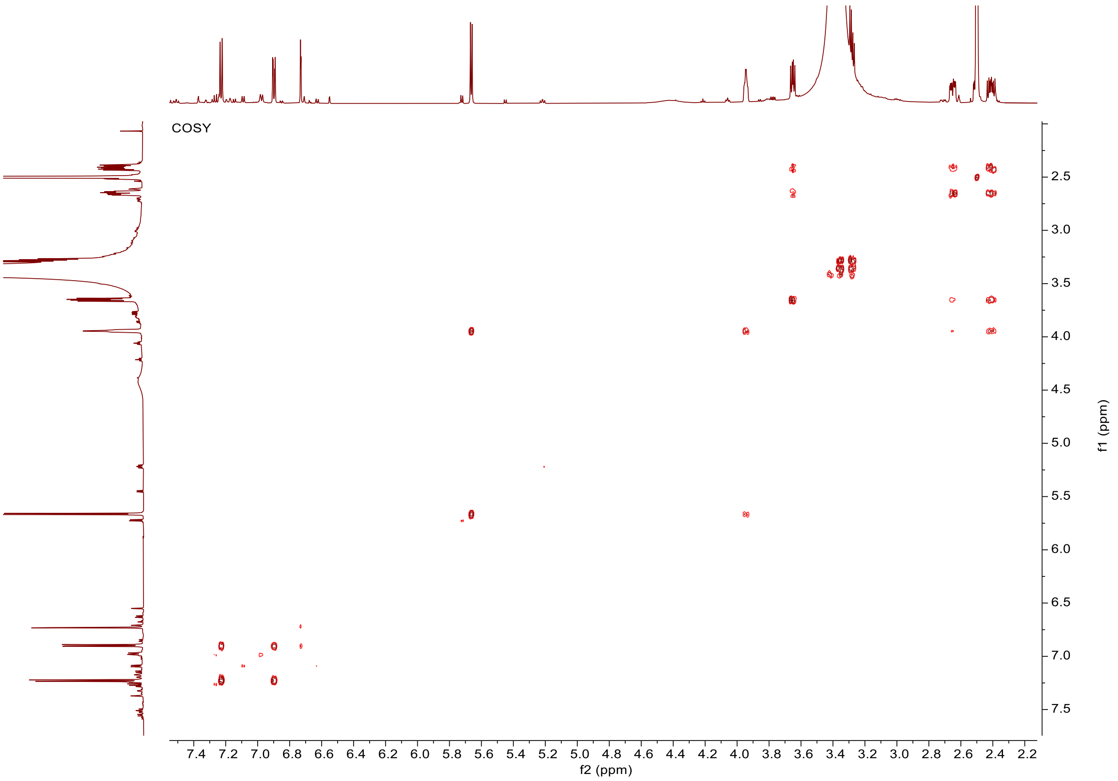

**Supplementary Fig. 30:** The $^{1}H$-$^{1}H$-COSY spectrum of compound **5** in DMSO-d*6*.

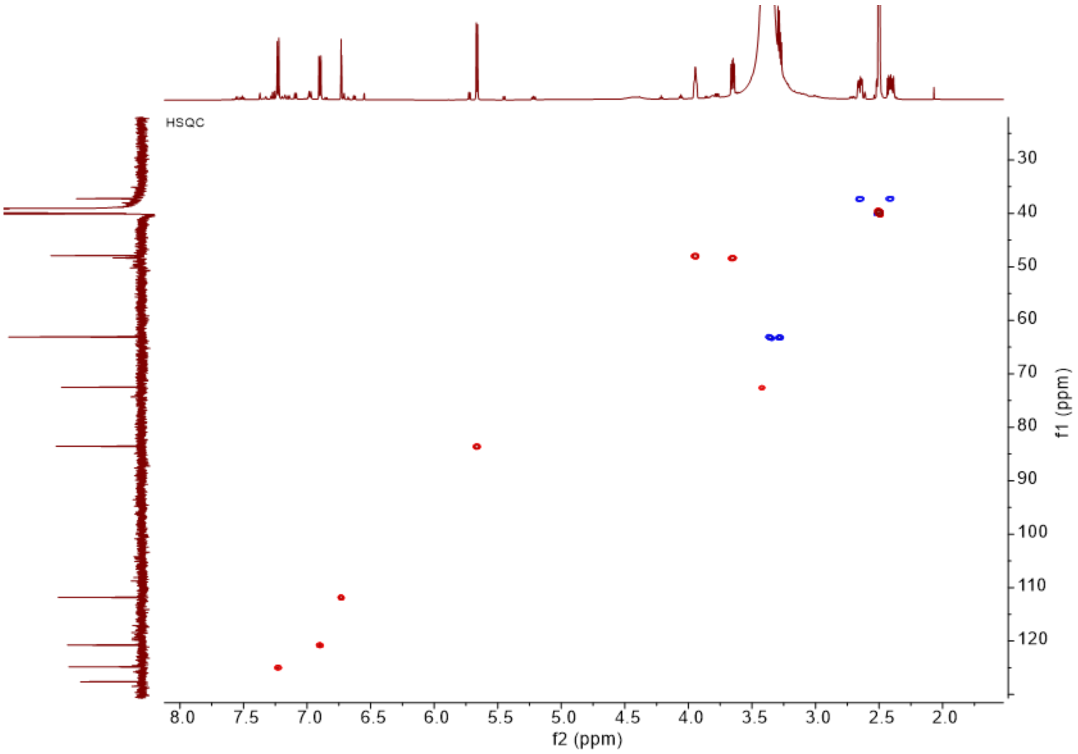

**Supplementary Fig. 31:** The $^{1}H$-$^{13}C$ HSQC spectrum of compound **5** in DMSO-d*6*.

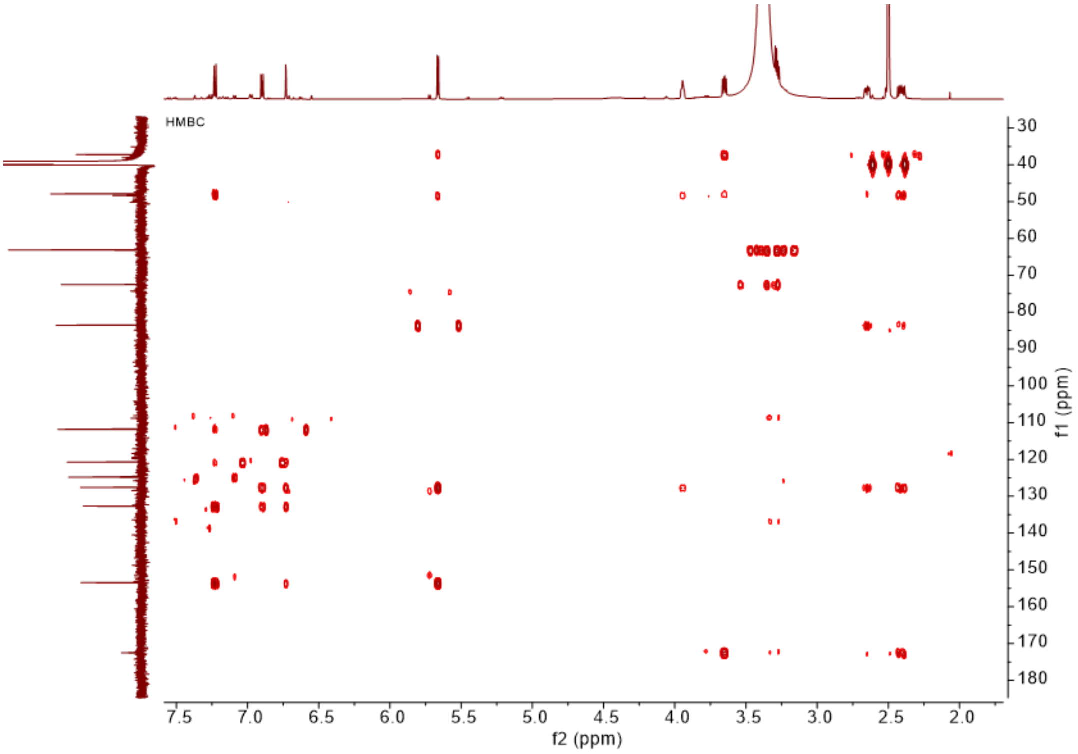

**Supplementary Fig. 32:** The $^{1}$H-$^{13}$C HMBC spectrum of compound **5** in DMSO-d*6*.

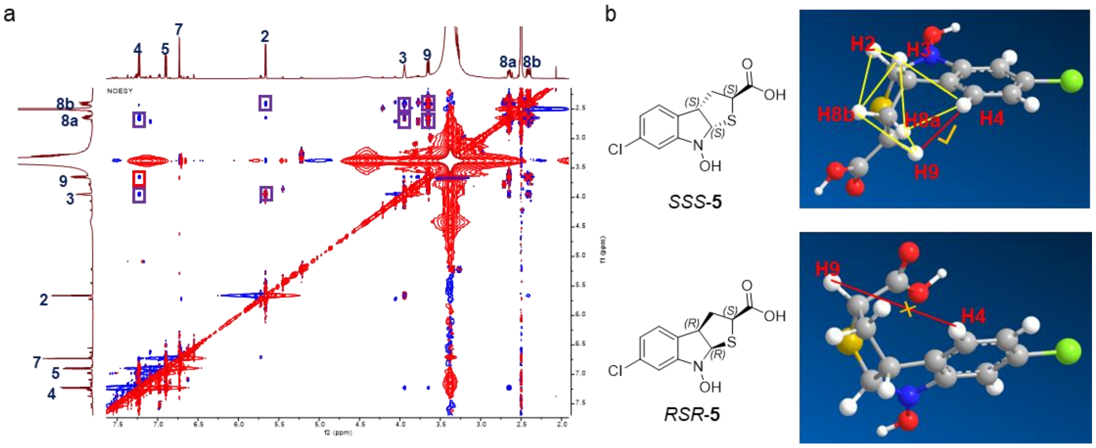

**Supplementary Fig. 33: Stereochemical analysis of compound 5. a**, The $^{1}$H-$^{1}$H-NOESY spectrum of compound **5** in DMSO-d*6*. **b**, Proposed stereochemical assignment. The key NOE correlation between H4 ($\delta$ 7.23) and H9 ($\delta$ 3.65) is consistent with the *S*, *S*, *S*-configuration. Structure generated with Chem3D 20.0.

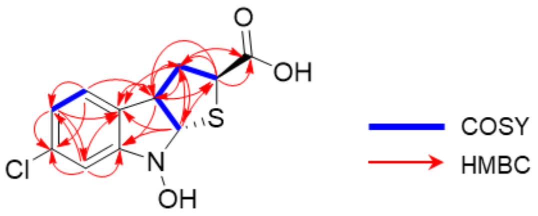

**5**

**Supplementary Fig. 34:** Key $^{1}$H-$^{1}$H COSY, $^{1}$H-$^{1}$H-NOSY and $^{1}$H-$^{13}$C HMBC correlations of compound **5**.

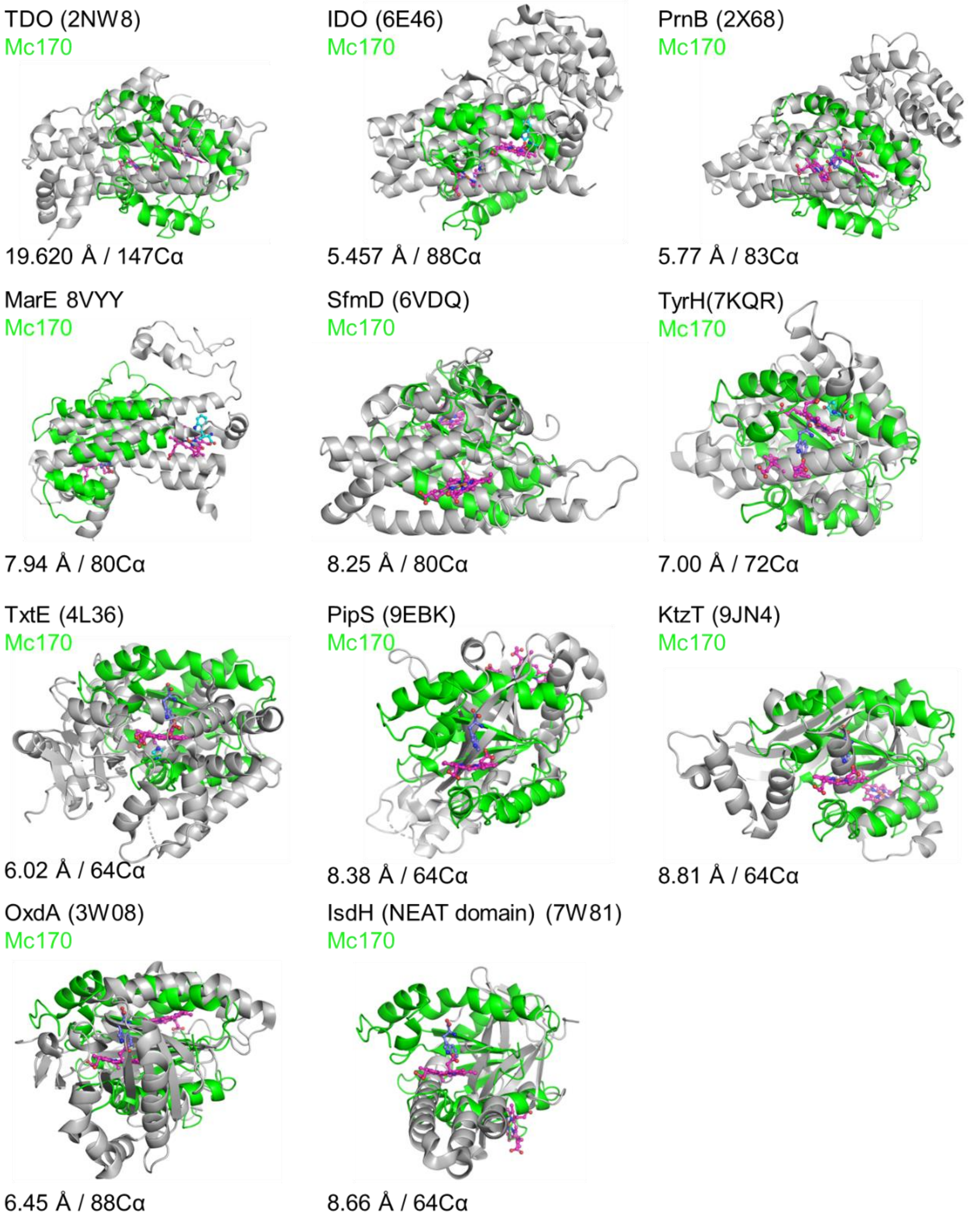

**Supplementary Fig. 35: Structural alignment of Mc170 with heme-dependent tryptophan metabolic enzymes.** Structural superpositions of Mc170 with chain A of reference heme-dependent enzymes (PDB: 2NW8-TDO, 6E46-IDO, 2X68-PrnB, 8VYY-MarE, 6VDQ-SfmD, 7KQR-TyrH, 4L36-TxtE, 9EBK-PipS, 9JN4-KtzT, 3W08-OxdA, 7W81-IsdH). Coloring scheme: heme groups (red), Mc170-bound tryptophan (purple), and cognate ligands in reference proteins (cyan).

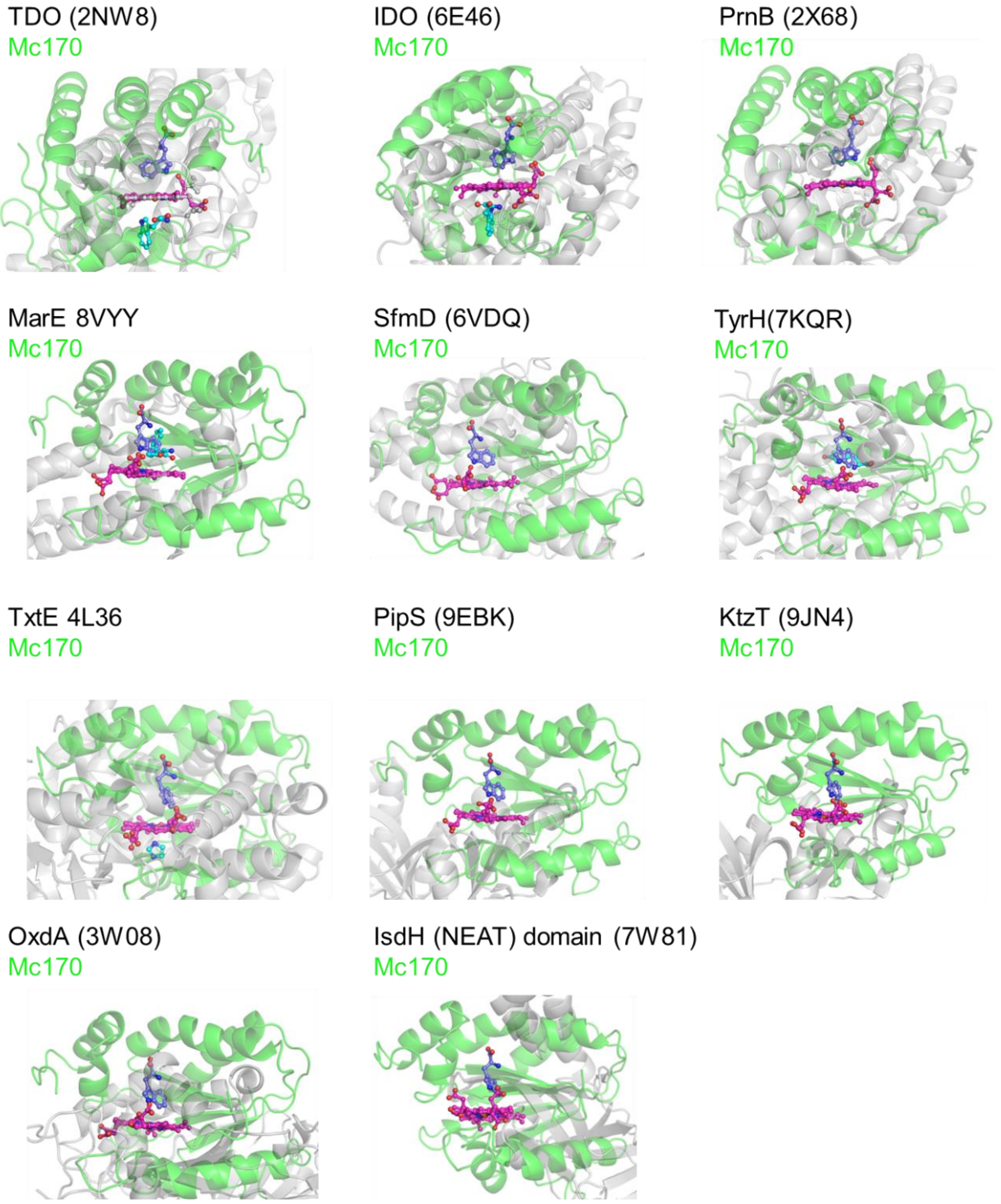

**Supplementary Fig. 36: Heme-plane-aligned structural superposition of Mc170 with tryptophan-metabolizing heme enzymes.** Structural alignments performed by superimposing the heme planes of Mc170 and Chain A of reference enzymes (PDB: 2NW8-TDO, 6E46-IDO, 2X68-PrnB, 8VYY-MarE, 6VDQ-SfmD, 7KQR-TyrH, 4L36-TxtE, 9EBK-PipS, 9JN4-KtzT, 3W08-OxdA, 7W81-IsdH). Coloring: heme cofactors (red), Mc170-bound tryptophan (purple), cognate substrates/products in reference structures (cyan).

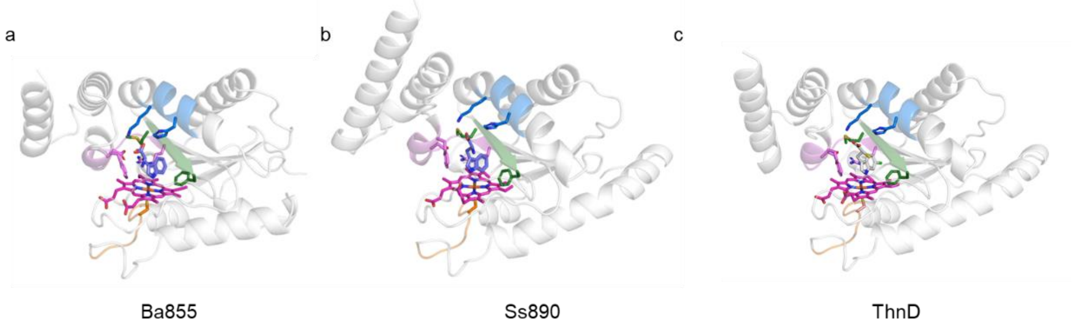

**Supplementary Fig. 37: Predicted ligand-bound structures of cytochrome P422 members.** Computational models of (a) Ba855/L-Trp, (b) Ss890/L-Trp, and (c) ThnD/ *S*-**4** complexes generated by AlphaFold3[4]. Structures shown as cartoon representations with ligands and key amino acid as sticks.

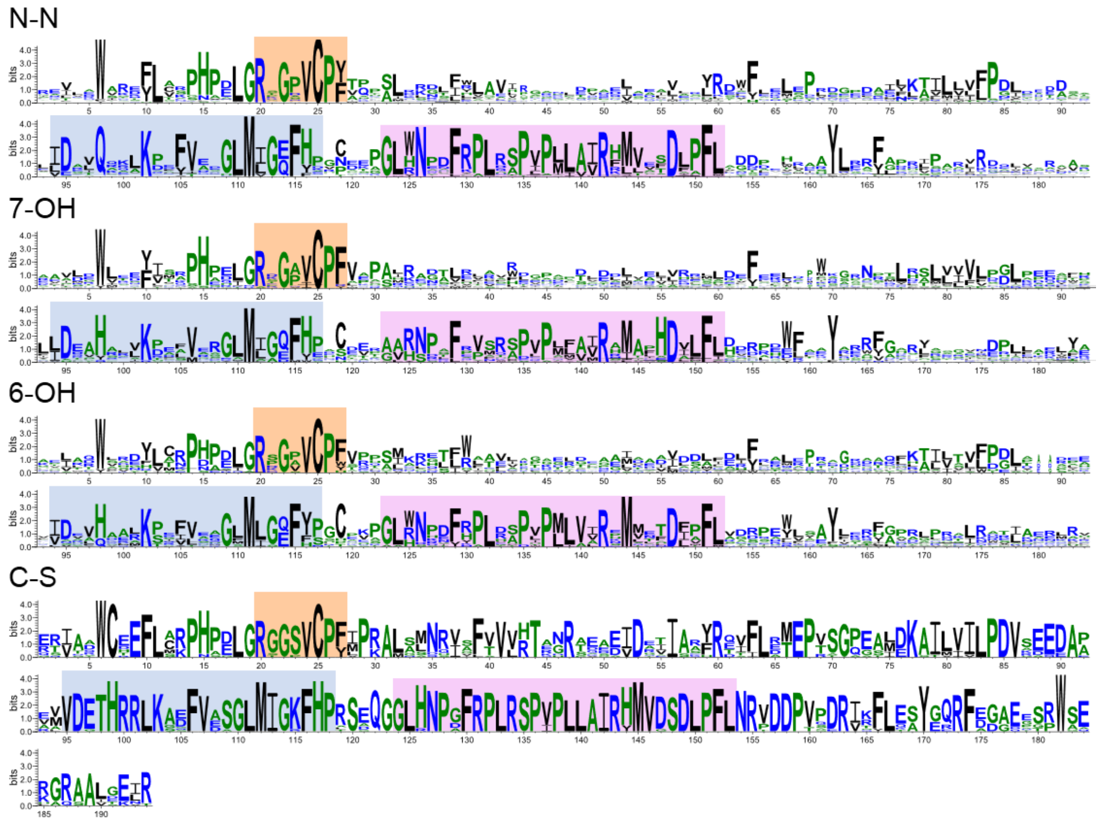

**Supplementary Fig. 38: Functional class-specific sequence conservation in cytochrome P422 enzymes.** Sequence logos constructed for four reaction classes using WebLogo 3[5]: N–N bond formation (N–N, n=41), C7-hydroxylation (7–OH, n=18), C6-hydroxylation (6–OH, n=15), and C–S bond formation (C–S, n=13). Conserved motifs shown: substrate-binding residues (blue/pink) and heme-binding signatures (orange). Sequence Sets:

N–N bond formation: The analysis included 41 sequences with the following UniProt IDs: A0A1Z1FA26, A0A3S2X2H1, A0A158HAL1, A0A149PGL8, A0A1I4PGA9, A0A7Y4Y5E2, A0A0Q8D691, A0A108UBG6, A0A8J6WP09, A0A978SBE7, A0A3S0ZQG6, A0A6P1KUG2, A0A139WVV4, A0AAP5M4T2, A0A6G9SJS3, A0A2H6LNR8, A0A6I3ZX66, A0A848KAF2,

F6EF82, A0A0D8HYK3, A0A917VT16, A0A562WKB5, A0A4R2J5W9, A0A229RLY4, A0A1H0M606, A0A9K3T6T0, A0A1Q5MKF1, A0A384HJB5, A0A2S9PXX9, A0A6G9FAW0, A0A556MU32, A0A410RXA3, A0A918UUE0, A0A542DBL9, A0A495X981, A0A1C4Z9T6, A0A7W7WSA9, D2AY02, A0A7W8EGY6, A0A5M3WAL5, A0A840NWY3.

C7-hydroxylation: The dataset comprised 18 sequences (UniProt IDs: C6WFN5, A0A6C0Q126, A0A853BV61, D6R234, A0A4U5W5J8, K8FE45, A0A3N6HL00, D6R241, A0A2R3ZQ27, K4RBM6, A0A154MRM9, A0A840Q4G5, A0A0G3APM5, A0A2N7WBB0, A0A4R5M467, A0A4R6F500, A0A837XV78, A0A856MJE7).

C6-hydroxylation: This class included 15 sequences (UniProt IDs: A0A7X1IS07, A0A1I2HNG7, A0A1Z2KYE7, A0AA91H005, D3VI28, A0A1I7GRM0, A0A089WT68, A0A0Q8BRF7, A0A543F3X1, A0A231H7J4, A0A562E549, A0A7K1UWC0, A0A6G9YPW7, A0A0Q7NTW8, A0A6G9XXX9).

C–S bond formation: To address the limited number of biochemically characterized enzymes for this reaction, a homology-based approach was used. Thirteen homologs with >65% identity to ThnD were retrieved via BLAST search against the NCBI database and included in the analysis (NCBI accession numbers: AMR44304.1, ANW12114.1, WP_260216142.1, WP_365631381.1, HJZ56405.1, WP_239335159.1, WP_098755158.1, WP_328629986.1, WP_384384459.1, WP_399053243.1, WP_438289717.1, XWD23044.1, WP_211768557.1).

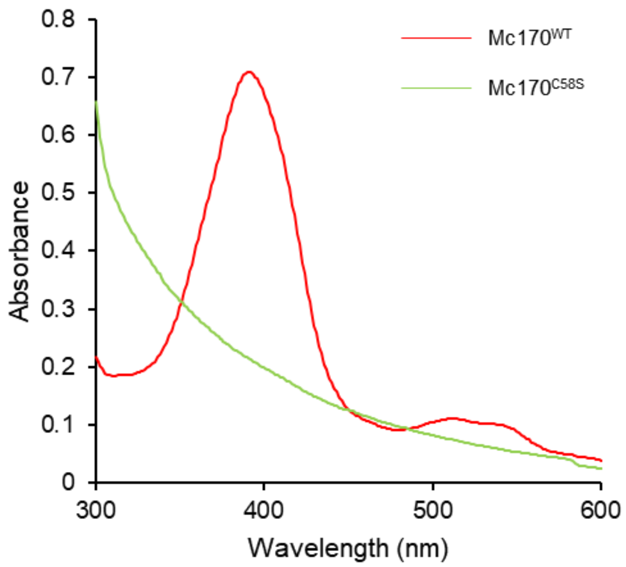

**Supplementary Fig. 39: UV-visible absorption spectra of Mc170 wild type (Mc170$^{WT}$) and the C58S mutant (Mc170$^{C58S}$).** The loss of enzymatic activity in the Mc170$^{C58S}$ variant is consistent with its severely impaired heme binding, as confirmed by UV-visible spectroscopy.

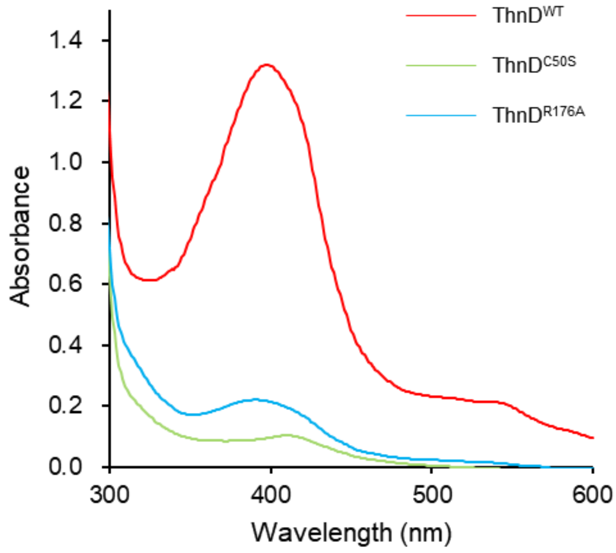

**Supplementary Fig. 40: UV-visible absorption spectra of ThnD wild type (ThnD$^{WT}$) and the C50S and R176A mutants (ThnD$^{C50S}$, ThnD$^{R176A}$).** The loss of enzymatic activity in both the ThnD$^{C50S}$ and ThnD$^{R176A}$ variants is consistent with their severely impaired heme binding, as confirmed by UV-visible spectroscopy.

**References**

1. Oberg, N., Zallot, R. & Gerlt, J. A. EFI-EST, EFI-GNT, and EFI-CGFP: enzyme function initiative (EFI) web resource for genomic enzymology tools. *J. Mol. Biol.* **435**, 168018 (2023).
2. Zallot, R., Oberg, N. & Gerlt, J. A. The EFI web resource for genomic enzymology tools: leveraging protein, genome, and metagenome databases to discover novel enzymes and metabolic pathways. *Biochemistry* **58**, 4169-4182 (2019).
3. Shi, X. *et al.* Hydroxytryptophan biosynthesis by a family of heme-dependent enzymes in bacteria. *Nat. Chem. Biol.* **19**, 1415-1422 (2023).
4. Abramson, J. *et al.* Accurate structure prediction of biomolecular interactions with AlphaFold 3. *Nature* **630**, 493-500 (2024).
5. Crooks, G. E., Hon, G., Chandonia, J. M. & Brenner, S. E. WebLogo: a sequence logo generator. *Genome Res.* **14**, 1188-1190 (2004).
